# P3a site-specific and cassette mutagenesis for seamless protein, RNA and plasmid engineering

**DOI:** 10.1101/2025.02.03.636246

**Authors:** Xiang-Jiao Yang

## Abstract

Site-directed mutagenesis is a basic molecular tool required for protein, RNA and plasmid engineering. For mutagenesis methods, an ideal goal is to reach the efficiency of 100%. Towards this goal, we have recently taken the first step by adopting an innovative strategy via primer pairs with 3′-overhangs and developing P3 site-directed mutagenesis, with an average efficiency of ∼50%. As the second step towards the ideal goal, we report here P3a site-directed mutagenesis with an efficiency reaching ∼100%. We systematically evaluated this new method by engineering >100 point mutations and small deletions (or insertions) on >20 mammalian expression vectors encoding various epigenetic regulators and the spike protein of SARS-CoV-2. As all known mutagenesis methods are limited to point mutations and small deletions/insertions (up to a dozen nucleotides), a technical problem is how to carry out cassette mutagenesis for replacement, deletion or insertion of large DNA fragments. The high efficiency of P3a mutagenesis and the ‘handshaking’ feature of primer pairs with 3′-overhangs inspired us to adapt this new method for seamless cassette mutagenesis, including highly efficient epitope tagging and untagging, deletion of small or large DNA fragments (up to 5 kb) and insertion of gene fragments (up to ∼0.4 kb), LoxP sites and sequences encoding degrons, sgRNA and tigRNA. Thus, this new site-specific and cassette mutagenesis method is highly efficient, fast and versatile, likely resulting in its wide use for typical biomedical research, as well as for engineering and refining synthetic or mutant proteins from AI-assisted design.

## Introduction

Site-directed mutagenesis is a basic method required for research in different branches of biomedical sciences [1]. Whole-genome and exome sequencing have identified numerous germline or somatic mutations in various diseases, as illustrated by genetic disease-linked mutations listed in the recently established ClinVar database [2] and cancer-associated mutations documented in the cBioPortal and COSMIC databases [3,4]. Because many of these mutations are missense and their functional impact is difficult to predict, an important question is to test whether these mutations are causal. For this, it is necessary to utilize site-directed mutagenesis and engineer the mutations for functional analysis *in vitro,* followed by generation and analysis of model organisms with equivalent mutations *in vivo*. Site-directed mutagenesis is also required for modeling evolution of viruses, such as SARS-CoV-2, to assess the impact of their newly acquired mutations [5,6]. This technique is also essential for protein engineering, such as testing mutant protein candidates from machine learning-based design [7] and generating artificial SH2 domains like phospho-tyrosine super-binders for deep proteomics and clinical profiling [8,9]. Other applications of protein engineering include generation of Cas9 variants, construction of super-antibodies and artificial intelligence (AI)-assisted *de novo* design of novel proteins for biomedical research, clinical diagnosis and therapy [10–13].

Furthermore, instead of targeting one or a few residues, ‘cassette’ mutagenesis replaces, deletes or inserts DNA fragments, so it is valuable for introducing small or large deletions, engineering epitope tags, optimizing plasmids, inserting different types of DNA sequences (such as LoxP sites, related DNA-recombination elements, promoters and the coding sequences for nuclear import and export signals) and constructing sgRNA and shRNA expression vectors. So far, the only means to carry out cassette mutagenesis is through subcloning, which is restricted to the availability of suitable restriction sites. Seamless cassette mutagenesis is an ideal choice and should have much wider applications.

Site-directed mutagenesis was initially developed for engineering mutations at specific sites of single-stranded phage or phagmid DNA [1,14]. By contrast, the QuickChange^™^ mutagenesis method utilizes double-stranded plasmids as the templates [15–17]. It is based on Pfu DNA polymerase-mediated PCR with a pair of completely complementary primers containing a given mutation (Fig. 1A), followed by DpnI digestion to destroy parental plasmids methylated at GATC sites. While it has been widely used in different laboratories, this method suffers from at least five limitations: (1) the complementary primers self-anneal and generate primer-primer dimers during PCR, thereby decreasing the amplification efficiency; (2) newly synthesized plasmid DNA is ’nicked’ and thus unsuitable as a template for further amplification; (3) unwanted mutations at the primer sites resulting from impurity of synthesized primers (this limitation applies to all site-directed mutagenesis methods); (4) unwanted mutations at the mutation and primer sites due to the processivity or strand-displacing activity of Pfu [18,19]; and (5) this polymerase synthesizes at a rather slow rate of ∼1 kb/min at 72°C, so it takes over 6 hours to complete a 25-cycle PCR reaction for a 13-kb plasmid. This can reach 10-12 hours if the extension is carried out at 68°C for such a plasmid, with parameters recommended by the manufacturer of Pfu_Ultra. Thus, it is ideal to reduce length of PCR time.

**Fig. 1.**
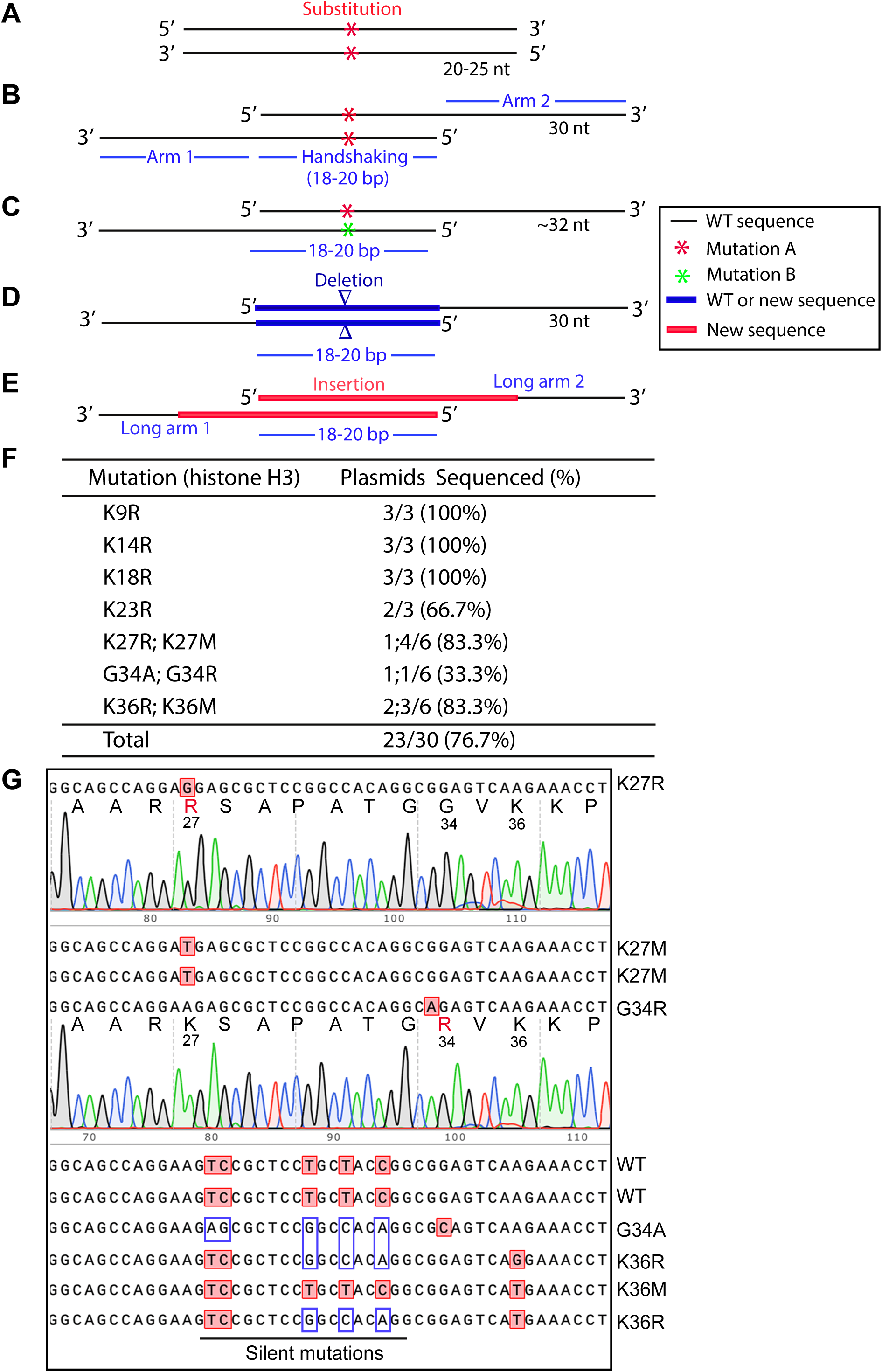
Primer design for P3a mutagenesis and efficient generation of histone H3 mutants. A. Completely complementary primer pair for the QuickChange^TM^ mutagenesis method [15,16]. Two red asterisks mark the mutation sites within the primer pair. The primers are 20-25 nucleotides (nt) in length and completely complementary to each other. B. Partially complementary primer pair with 3′-overhangs. Two red asterisks mark the mutation sites within the primer pair. Liu & Naismith utilized primers with the average length of ∼45 nucleotides (ranging from 35 to 55 nucleotides) in length, with special instructions on primer design [21]. We have simplified the primer design and reduced the length of the two primers to ∼30 nucleotides, with 9-bp complementary regions flanking a single-nucleotide mutation site [18]. Shorter primers reduce cost and also minimize unwanted mutations introduced by the impurity of primers resulting from their chemical synthesis [18,19]. C. The two primers for engineering the mutation can carry a mismatch at the same position or two slightly different positions as indicated with red and green asterisks. In these cases, the primers are slightly longer than 30 nucleotides, with 9-10 bp complementary regions flanking a single-nucleotide mutation site. With such design, two different mutants can be generated with one pair of primers [18]. D. A pair of ‘handshaking’ primers to introduce deletion (denoted with two red triangles) via a pair of primers possessing two arms complementary to the flanking regions. The deletion can be from a few base pairs to several kb. For deletion of large DNA fragments, it is impossible or highly inefficient for all classical site-directed mutagenesis methods. Conceptually, the primer pairs in strategies depicted in panels D-E form cassettes for replacing the corresponding regions in the wild-type backbone. E. A pair of ‘handshaking’ primers to introduce insertion (denoted with red solid lines) through a pair of primers containing two arms complementary to the regions (indicated by solid dark lines) flanking an insertion site. For insertion of large DNA fragments, it is impossible or typically inefficient for all classical site-directed mutagenesis methods. Importantly, this strategy can be used for replacing DNA fragments. Conceptually, deletion and insertion can be considered as two special cases where the respective sequences to be inserted and deleted are zero bp. This is because replacement mutagenesis means conversion of fragment A to fragment B. For deletion, fragment B is zero bp in size, whereas for insertion, fragment A is zero bp in size. F. Efficiency for generating different histone H3 mutants. The notation “K27R;K27M” refers to generating two mutants K27R and K27M, whereas the term “1;4/6” denotes, respectively, one and four K27R and K27M mutants out of 6 colonies sequenced. See Fig. S1 for details on P3a mutagenesis and about how the primers are designed. G. Sequence chromatograms or sequences of 10 representative plasmids analyzed for engineering histone H3 mutants. The parental plasmid encodes *Xenopus* histone H3. During sequencing, we noticed that the plasmid used for mutagenesis contains silent mutations compared to the sequence that we used to design the primers, which is expected to result in more mismatches between the primers and template. Amazingly, despite these mismatches, the primers successfully introduced the designed mutations and corrected some of the silent mutations.

Mitigating limitations (1) and (2), an alternative approach utilizes a pair of partially complementary primers with two 3′-protruding ends (Fig. 1B), which allows the use of newly synthesized strands as templates for subsequent PCR amplification (Fig. S1A) [20,21]. This strategy was initially introduced by Zheng, Baumann & Reymond [20]. One drawback of this previous study was that the 5′ arms from the desired mutation sites are only 3-5 nucleotides long for some primers, which may lead to insufficient annealing to and priming from newly synthesized strands. Moreover, restriction digestion at coupled secondary sites, rather than DNA sequencing, was used to assess the mutagenesis efficiency, so the real success rate remains uncertain [20]. Unwanted mutations at and around the desired mutation sites were not assessed at all. This method was later refined by Liu & Naismith, leading to its broader application [21]. However, the method was developed and tested only for two 5-6 kb plasmids [21], so it is unclear how the method performs with larger plasmids and those with more complex sequences, as site-directed mutagenesis becomes more challenging with larger plasmids [22–24]. Moreover, only 8 point mutations were tested in the previous study and two of them failed [21], so further improvements are required to establish the general applicability of the method.

To address limitation (3) mentioned above, we have shortened the length of primers to ∼30 nucleotides (Fig. 1B) as shorter oligonucleotides tend to be synthesized more easily at high quality. Related to limitation (4), we utilized PfuUltra (an improved version of Pfu) [25]. As <u>p</u>rimer pairs with <u>3</u>′-protruding ends are used, this improved method has been referred to as P3 site-directed mutagenesis [18,19]. Moreover, we have systematically evaluated this method (Fig. 1B-C) with a dozen mammalian expression vectors ranging from 7.0-13.4 kb [18,19]. Compared to the QuickChange^™^ method, the success rate has increased significantly but still varies, with an average efficiency of about 50% [18,19], instead of being almost 100% as reported in the previous study [20]. The difference could be size and sequence differences of the plasmids used in these two studies. In this previous study [20] and also our recent study [18], problems still occurred with certain mutations and/or plasmids. Thus, from the technological point of view, it would be ideal to raise this efficiency near or to 100% for various mutations and plasmids. Another issue is that only point mutations were tested in our recent study [18], so it is unclear whether the method works with deletion and insertion, especially large ones. In the previous study from another group, only mall deletion or insertion (up to about a dozen base pairs) have been tested [21], so it is remains to be investigated whether the method works with large deletion and insertion. In fact, no effective methods have been reported in this regard. This is important as it is known that it much more difficult to engineer large deletion or insertion than small ones.

Related to limitation (5) mentioned above, we utilized Pfu_fly, an improved version of Pfu with a much faster DNA synthesis rate [18,19]. But it tends to introduce undesired mutations [18,19]. Thus, alternatives are urgently needed. To improve the P3 mutagenesis method further [18,19], we sought to overcome limitations (4) & (5) by replacing Pfu DNA polymerase with thermostable DNA polymerases with higher fidelity and faster synthesis rates. As a result, we tested 4 candidates, two of which are: 1) Q5 high-fidelity DNA polymerase, composed of a thermostable DNA polymerase fused to the processivity-enhancing Sso7d DNA-binding domain, thereby improving speed, fidelity and reliability of PCR amplification [26]; and 2) Platinum SuperFi II polymerase, whose fidelity was claimed to be even better than Q5 DNA polymerase. Results from >100 mutations that we have engineered with various plasmids (up to 13.4 kb in size) indicate that these two polymerases are superior to Pfu and its derivatives by elevating the mutagenesis efficiency very close to 100%, reducing PCR length by 2-3 folds and minimizing unwanted mutations at or near the primer sites. Due to the much higher success rate, only 1-2 bacterial colonies need to be sequenced per mutation reaction. Moreover, fewer complete cells and bacterial plates are needed for transformation of the mutagenesis reaction mixtures. All of these reduce labor and reagent costs. To our knowledge, this is the first site-specific mutagenesis method to achieve the efficiency so consistently at or close to the ideal level of 100%, for various plasmids up to 13.4 kb in size.

Subsequently, in light of the unique ‘handshaking’ feature of the primers with 3′-protruding ends (Fig. 1B), we utilized the method to engineer highly efficient cassette mutagenesis (Fig. 1D-E),including seamless epitope tagging or untagging, deletion of small or large fragments (up to 3.2 kb), and insertion of oligonucleotide duplexes with 3′-overhangs (potentially up to 0.4 kb). Notably, the handshaking portions of the primer pairs does not need to be the wild-type but can be completely new sequences even for deletion (Fig. 1D). Conceptually, we consider deletion and insertion as two special cases of DNA conversion (or replacement) where the respective sequences to be inserted and deleted are zero bp (Fig. 1D-E). This is significant because to our knowledge, no such methods have been reported yet. The reagent cost per mutagenesis reaction is only about $0.25 and $0.5 with Q5 and SuperFi II master mixes, respectively. Compared to those based on Pfu and its derivatives [18,20,21], this newly developed mutagenesis method is not only much faster but also more efficient, versatile and economical, likely making it a standard tool in various biomedical laboratories. Thus, the current study nicely complements and extends the previous three studies based on primer pairs with 3′-overhangs [18,20,21].

## Results

### Improving P3 site-directed mutagenesis

We have recently evaluated the P3 site-directed mutagenesis method that relies on PfuUltra (Fig. 1B-D) [18,19]. The method reached an average efficiency of nearly 50% but encountered problems with some vectors [18]. To improve the method further, we considered to replace PfuUltra with other high-fidelity thermostable DNA polymerases comparable with or superior to PfuUltra in terms of PCR performance. We tested 4 candidates: (1) Q5 high-fidelity DNA polymerase, (2) Platinum SuperFi II high-fidelity DNA polymerase, (3) i7^®^ high-fidelity DNA polymerase and (4) KOD high-fidelity DNA polymerase. In terms of mis-incorporation rates, the former three were estimated to be ∼15 times lower than PfuUltra, whereas KOD high-fidelity DNA polymerase is comparable to Pfu. Moreover, commercially available master mixes containing Q5 high-fidelity DNA polymerase, Platinum SuperFi II high-fidelity Polymerase or i7^®^ high-fidelity DNA polymerase contain hot-start and other ingredients to enhance PCR performance, especially for templates with high GC-content and complex DNA sequences. As described below, Q5 and Platinum SuperFi II high-fidelity DNA polymerases outperformed PfuUltra, increasing the mutagenesis efficiency to ∼100%, shortening the PCR amplification time by 2-3 folds and reducing unwanted mutations at or near the primer sites. Compared to Q5, Platinum SuperFi II DNA polymerase yielded more colonies. KOD DNA polymerase was comparable to PfuUltra, whereas i7^®^ high-fidelity DNA polymerase yielded very few or no colonies at all. Compared to PfuUltra [18], Q5 and Platinum SuperFi II high-fidelity DNA polymerases yielded 5-10 more colonies per mutagenesis reaction, which enhances the success rate substantially. While it remains unclear why i7^®^ high-fidelity DNA polymerase did not produce sufficient colonies as expected, the superior performance of Q5 and Platinum SuperFi II high-fidelity DNA polymerases is in agreement with their high processivity, fast synthesis rate and low mis-incorporation rate. In light of these observations, we chose Q5 and Platinum SuperFi II high-fidelity DNA polymerases for the rest of this study. To distinguish it from Pfu-based P3 mutagenesis [18,19], this newly developed method is referred to as P3a site-directed mutagenesis, where the letter “a” denotes an updated version of the P3 method.

### P3a site-specific mutagenesis for efficiently engineering missense mutants of histone H3

Histone H3 is almost invariant from yeasts to humans and plays an important role in organizing the genomes and their access for various DNA-based nuclear processes, such as DNA replication, repair and RNA transcription [27]. At its N-terminal tail, there are multiple lysine residues (i.e., K4, K9, K14, K18, K27 and K36) for acetylation, methylation and other modifications, thereby bringing about various regulatory mechanisms [27]. Moreover, substitutions of K27, G34 and K36 have been linked to oncogenesis, which has led to the onco-histone hypothesis [28]. To test P3a mutagenesis, we sought to engineer different histone H3 mutants as they are important for molecularly investigating the role of these residues by analyzing the impact of their substitutions. As shown in Fig. 1F-G, for the K9R, K14R, K18R and K23R mutants, we utilized 4 different primer pairs, such as K9R-F and K9R-R for engineering the mutation K9R (Fig. S1B). We sequenced plasmids from three of the resulting colonies per mutation and found that 11 of them were mutants, reaching an overall efficiency of 11/12 (91.6%, Fig. 1F). This is impressive as we have not seen any known site-directed mutagenesis methods with such an almost ideal success rate.

One drawback of P3 or P3a site-directed mutagenesis is the need to use a pair of primers for engineering one mutation, which is different from classical methods that utilize only one primer per mutation. Thus, we next utilized the primer design strategies described in Fig. 1C to generate two mutants per primer pair. For example, we employed one primer pair (K27R-F and K27M-R) with a mismatch for generating two mutations (K27R and K27M; Fig. S1B). Similarly, we designed the primers G34R-F and G34A-R with two mismatched bps for generating the mutations G34R and G34A, respectively (Fig. S1C). For this, we analyzed 6 colonies per primer pair and found 2 to 5 colonies containing plasmids with the expected mutations (Fig. 1F), thereby reaching an overall efficiency of 12/18 (66.7%). Typically, for P3a mutagenesis, it is often adequate to sequence 2 colonies per mutation, thereby reducing labor and reagent costs. Thus, this newly developed method is efficient and economical.

### P3a mutagenesis for engineering missense mutants of two epigenetic regulators

To assess the versatility of the method, it is necessary to test it with a range of vectors, including those much larger than the one described above for histone H3 (only ∼5.7 kb). In addition to histones, there are hundreds of epigenetic regulators that modify or bind to histones for regulating genome organization during different nuclear processes. To study molecular functions of these epigenetic regulators, it is necessary to engineer various mutants. We thus sought to apply P3a mutagenesis for rapidly generating such mutants. Another practical reason is that the expression vector for histone H3 is ∼5.7 kb, which is small compared to many expression vectors for epigenetic regulators. Typically, it becomes more difficult for PCR-based mutagenesis as the vector becomes larger. One epigenetic regulator is BRPF1 (bromodomain- and PHD finger-containing protein 1) [29]. It was originally identified as a 140 kDa protein with a bromodomain [29], so its expression vector reaches ∼9.2 kb and is significantly larger than that for histone H3.

In addition to the bromodomain, BRPF1 possesses multiple other domains for epigenetic regulation, including the N-terminal part for interacting with lysine acetyltransferase 6A (KAT6A) and its paralog KAT6B [30]. The interaction is required for activation of these acetyltransferases to acetylate and propionylate nucleosomal histone H3 [31,32]. In addition, BRPF1 possesses two EPC (Enhancer of Polycomb)-like motifs, the second of which interacts with ING4 (or its paralog ING5) and MEAF6 [33]. Moreover, BRPF1 has two PHD fingers flanking a zinc knuckle, with the first PHD finger for recognition of the unmodified N-terminus of nucleosomal histone H3 [34]. Recent studies have linked *BRPF1* mutations to a neurodevelopmental disorder [31,35–38]. According to the ClinVar database [2], additional *BRPF1* mutations have been identified in patients. While some of the identified mutations have been molecularly characterized [31,35,36], many more remain to be analyzed for firmly establishing their pathogenicity. For molecular characterization of different BRPF1 domains, we also needed to engineer more mutants (Fig. 2A). We sought to employ P3a mutagenesis to engineer new BRPF1 mutants. As shown in Fig. 2A, among 31 bacterial colonies analyzed, 28 contained the mutant plasmids, so the efficiency was 28/31 (90.3%).

**Fig. 2.**
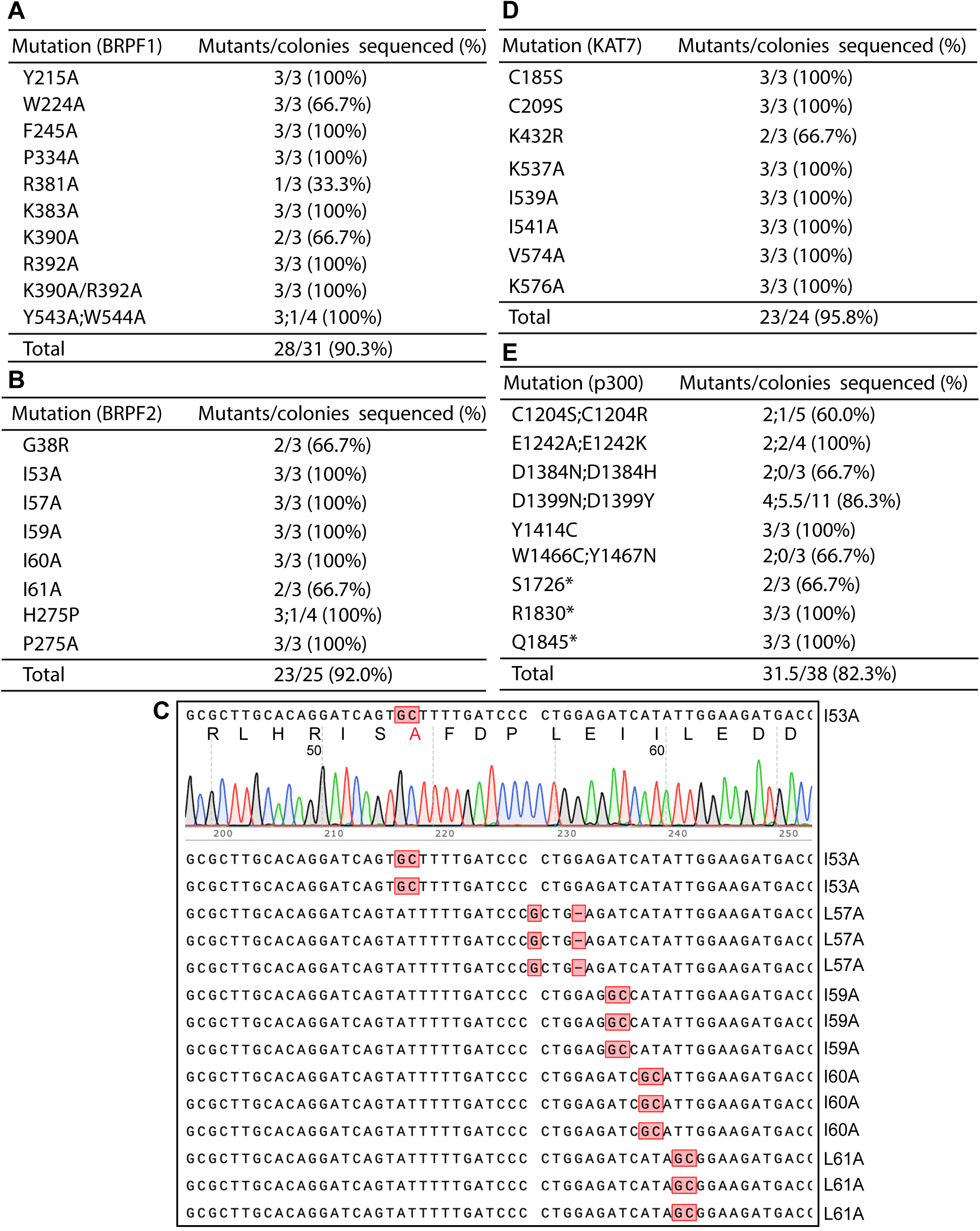
Engineering missense or nonsense epigenetic regulator mutants by P3a site-specific mutagenesis. A. Efficiency in engineering eleven BRPF1 mutants. The notation ‘Y543A;W544A’ refers to generating the two mutants Y543A and W544A with just one pair of primers (see Fig. 1C), whereas the term ‘K390A/R392A’ denotes the double mutant containing both substitutions. B. Efficiency in engineering eight BRPF2 mutants. Among them, G38R is to investigate a polymorphic substitution, whereas H275P is to repair an unexpected substitution in an expression plasmid that we have in the lab. The remaining mutants are located at a KAT7-binding site mapped by structural analysis [42]. C. A representative sequence chromatogram and the sequences of 15 plasmids analyzed for engineering BRPF2 mutations. The plasmids were subject to Sanger sequencing and the resulting data were analyzed via SnapGene. D. Efficiency in generating eight KAT7 mutants. The first two alter a zinc finger, whereas the remaining 6 mutants are located at a BRPF2-binding site mapped by crystal structural analysis [42]. E. Efficiency in engineering fourteen p300 mutants. The notation ‘C1204S;C1204R’ refers to generating two mutants Y543A and W544A with just one pair of primers (see Fig. 1C). The asterisks in S1726*, R1830* and Q1845* denote stop codons. See Fig. S2 for mutagenesis efficiency for four other epigenetic regulators.

At the sequence and domain-organization levels, BRPF1 is highly homologous to BRPF2, but KAT7 is a preferred partner of BRPF2 for acetylating nucleosomal histone H3 [33,39–41]. Functionally different from BRPF2, BRPF1 activates KAT6A and KAT6B [31,35]. Crystal structural analysis has identified a hydrophobic interface for interaction between BRPF2 and KAT7 [42]. This involves residues I53, L57, I59, I60 and L61 of BRPF2 [42]. No functional validation has been reported on the functional importance of this physical interaction. One approach is to engineer BRPF2 mutations at this interface. We thus utilized P3a mutagenesis to generate such mutants, including I53A, L57A, I59A, I60A and L61A (Fig. 2B). Moreover, we engineered G38R, H275P and P275A, to investigate the importance of a polymorphic substitution, correct an unexpected substitution identified in an expression plasmid that we have, and investigate the functional importance of P275, respectively. Amazingly, 23 out of 25 colonies that we analyzed were the correct mutants, reaching the efficiency of 92% (Fig. 2B). Representative sequence analysis results are shown in Fig. 2C. In another experiment, we engineered 11 point mutants of BRPF2 using 10 pairs of primers. We sequenced plasmids from 2-3 colonies per mutagenesis reaction. 22 out of 23 of the resulting colonies were correct, so the mutagenesis efficiency was 95.7%. The BRPF2 expression vector is ∼9.2 kb and similar to that for BRPF1 in size, so these results indicate that P3a mutagenesis is highly efficient for vectors up to 9.2 kb. If an empty vector itself is 5-6 kb, an insert can be 4-5 kb and encode a protein up to 150 kDa. This is important because a majority of proteins encoded by various genomes range from 30-150 kDa, so the method is expected to be applicable to many proteins.

### P3a mutagenesis for engineering mutants of small and large histone acetyltransferases

To evaluate the versatility of this new method, we also tested it with various expression vectors of different sizes. For this, we targeted histone acetyltransferases as they are our main research subjects [43]. The human genome encodes over a dozen of such enzymes [44]. As mentioned above, one such enzyme, KAT7, interacts with (and is activated by) BRPF2. Crystal structural analysis has identified a surface for interaction with a hydrophobic core of BRPF2 [42]. This involves five residues (K537, I539, I541, V574 and K576) for interacting with I53, L57, I59, I60 and L61 within the hydrophobic core of BRPF2, [42]. With P3a mutagenesis, we carried out alanine substitutions of these five residues of KAT7 [42]. Moreover, we sought to substitute two zinc-chelating residues within an uncharacterized zinc finger of KAT7 and replace its K432, a potential autoacetylation site that has never been characterized [45]. As shown in Fig. 2D, 23 of 24 colonies that we analyzed were mutants, thus reaching a high efficiency of 95.8%, further attesting to effectiveness of the method.

To assess the general applicability of the method further, we next tested it with six different mammalian expression vectors encoding five different lysine acetyltransferases, including KAT6A, KAT6B, KAT8, p300 and CBP. The expression vectors range from 6.6-13.4 kb in size: human KAT6A (full-length), 11.9 kb; its HAT domain, 6.6 kb; the HAT domain of KAT6B, 6.8 kb; KAT8, 7.0 kb; p300, 12.8 kb and mouse CBP, 13.4 kb. Among them, the expression vectors for full-length KAT6A, p300 and CBP are quite large, ranging from 11.9-13.4 kb, so they are good candidates for testing the versatility of P3a mutagenesis with large vectors. This is important as PCR-based mutagenesis methods are more efficient with small plasmids and tend to encounter problems with large plasmids, especially those close to or over 12 kb.

KAT6A and KAT6B interact with BRPF1 for acylating histone H3 [31,32]. For full-length KAT6A, we engineered two missense mutants and three nonsense mutants (Fig. S2A). In addition, we generated five missense mutants of its HAT domain (Fig. S2A). While K604R possesses an arginine substitution of K604, the sole autoacetylation site required for the acyltransferase activity of KAT6A [31], the other mutants of KAT6A and its HAT domain are clinical variants identified in patients with a neurodevelopmental disorder [2]. For example, L1061* is derived from a patient with intellectual disability [31]. These clinical variants have not been adequately analyzed for their pathogenicity, so it is necessary to engineer them for functional analysis *in vitro*. We thus applied P3a mutagenesis to engineering these mutants. For the five mutations on the backbone for full-length KAT6A, we analyzed plasmids from 23 bacterial colonies by Sanger sequencing and identified 19 as correct mutants, reaching efficiency of 82.6% (Fig. S2A). This result is significant as the vector is quite large, 11.9 kb. For the five mutations engineered on the vector for the HAT domain of KAT6A, 9 out of 12 colonies analyzed possessed the mutant plasmids (Fig. S2A), yielding an efficiency of 75%. The overall mutagenesis efficiency for these two KAT6A expression vectors was 80.0% (Fig. S2A).

KAT6B is paralogous to KAT6A. For KAT6B, we engineered 7 missense mutants on an expression vector for the HAT domain (Fig. S2B). These missense mutants are based on clinical variants identified in patients with a neurodevelopmental disorder [2]. These clinical variants have not been analyzed for their pathogenicity, so it is important to engineer them for functional analysis *in vitro*. Moreover, to define the boundary of the HAT domain, it is necessary to generate two deletion mutants (Fig. S2B). We utilized P3a mutagenesis to engineer these 9 missense or deletion mutants. As shown in Fig. S2B, plasmids from 26 out of 27 colonies analyzed by Sanger sequencing were the mutants. Thus, the efficiency was 26/27 (96.3%), very close to the ideal level of 100%.

We next tested KAT8, which shares its catalytic domain with KAT6A, KAT6B and KAT7 [43]. Importantly, *KAT8* mutations have recently been linked to a new neurodevelopmental disorder [46]. Moreover, additional missense mutants of KAT8 have been identified as clinical variants in patients with this disorder [2]. These clinical variants have not been analyzed for their pathogenicity, so it is important to engineer them for functional analysis *in vitro*. We thus applied P3a mutagenesis to engineer 13 missense mutants on a mammalian expression vector encoding FLAG-tagged KAT8. The size of this vector is 7.0 kb. As shown in Fig. S2C, plasmids from 24 out of 31 colonies analyzed by Sanger sequencing were the mutants. Thus, the mutagenesis efficiency was 24/31 (77.4%).

In the superfamily of lysine acetyltransferases encoded by the human genome, p300 and CBP are the largest, each possessing ∼2,400 residues. Their mammalian expression vectors reach 12.8 and 13.4 kb in size. They are among the largest plasmids that have been constructed. Thus, they are good candidates for testing the versatility of the P3a mutagenesis method. For p300, we sought to engineer 11 missense mutants and 3 C-terminal truncation mutants (Fig. 2E). Some of these mutants are derived from somatic mutations identified in cancer and deposited in the COSMIC and cBioPortal databases [3,4]. For example, D1399N, D1399Y, Y1414C, W1466C and Y1467N are five hotspot mutations [3,4]. The others (such as C1204S/R, D1384N/Y and the three truncation mutants) are to understand functions of different domains of p300. As shown in Fig. 2E, plasmids from 31.5 out of 38 colonies analyzed by Sanger sequencing carried the expected mutations, resulting in the mutagenesis efficiency of 82.3%.

For CBP, we sought to engineer 7 missense mutants and 2 C-terminal truncation mutants. The first four mutants (i.e., R1446C, R1466H, Y1503D and Y1503H; Fig. S2D) are derived from somatic hotspot mutations identified in cancer and deposited in the COSMIC and cBioPortal databases [3,4]. The remaining three missense mutants (K1832E, R1869Q and R1968W) and the two truncation mutants are to understand functions of different domains of CBP. As shown in Fig. S2D & S2E plasmids from 17 out of 20 colonies analyzed by Sanger sequencing carried the expected mutations. Thus, the efficiency was 85.0%. Together with the high efficiency to engineer full-length KAT6A and p300 mutants, these results support that P3a mutagenesis is also highly efficient for engineering mutants encoded by large plasmids.

### P3a mutagenesis for engineering missense spike variants of SARS-CoV-2

The coronavirus disease 2019 (COVID-19) pandemic resulted in tragic loss of life, affected health care and crippled economies around the world. Severe acute respiratory syndrome coronavirus 2 (SARS-CoV-2) is the underlying viral agent whose sequence was determined in January 2020 [5]. This virus has evolved rapidly, gained various mutations and yielded many variants and subvariants [6]. Notably, only some of these mutations act as drivers and shape up the evolutionary trajectories. Derived from Omicron variant, JN.1 and its descendants are currently contributing to active infections around the world. One major gene of variations encodes the spike protein, also the target of vaccines that have been or are being developed.

Compared to Omicron variant, JN.1 and its descendants have gained additional substitutions in the spike protein, including R346T, L455S, F456L, Q493E and V1104L [47]. To elucidate the underlying pathological mechanisms, it is necessary to engineer such substitutions in the wild-type spike protein for analysis of their functional impact using molecular and cell-based assays *in vitro*. We thus investigated whether P3a mutagenesis can be employed for engineering these mutations in the backbone of spike proteins of the D614G and Omicron variants. The D614G variant is a SARS-CoV-2 derivative that appeared in the early part of 2020 and led to development of many major variants that drove the pandemic. We also engineered L981F in the backbone of the spike protein of the D614G variant as this substitution is present in Omicron variant [6]. For L455S and F456L, we only utilized two primers, with one encoding F456L and another for both L455S and F456L, using the strategy illustrated in Fig. 1C. As shown in Fig. 3, among 32 colonies that we analyzed, 29 carried the correct mutant plasmids and one was a mixed clone of the wild-type and a mutant, so the overall mutagenesis efficiency was 92.2%. Once the primers were available, the initial mutagenesis experiment was carried out on day 1 and the colonies were inoculated on day 2. On day 3, the plasmids were prepared and sent for Sanger sequencing. Typically, 1-3 days later, the sequencing results were available for analysis. Thus, the mutant plasmids could be obtained easily within a week. Thus, this method is applicable for rapidly generating and molecularly modeling new mutations that JN.1 and its descendants will gain during evolution.

**Fig. 3.**
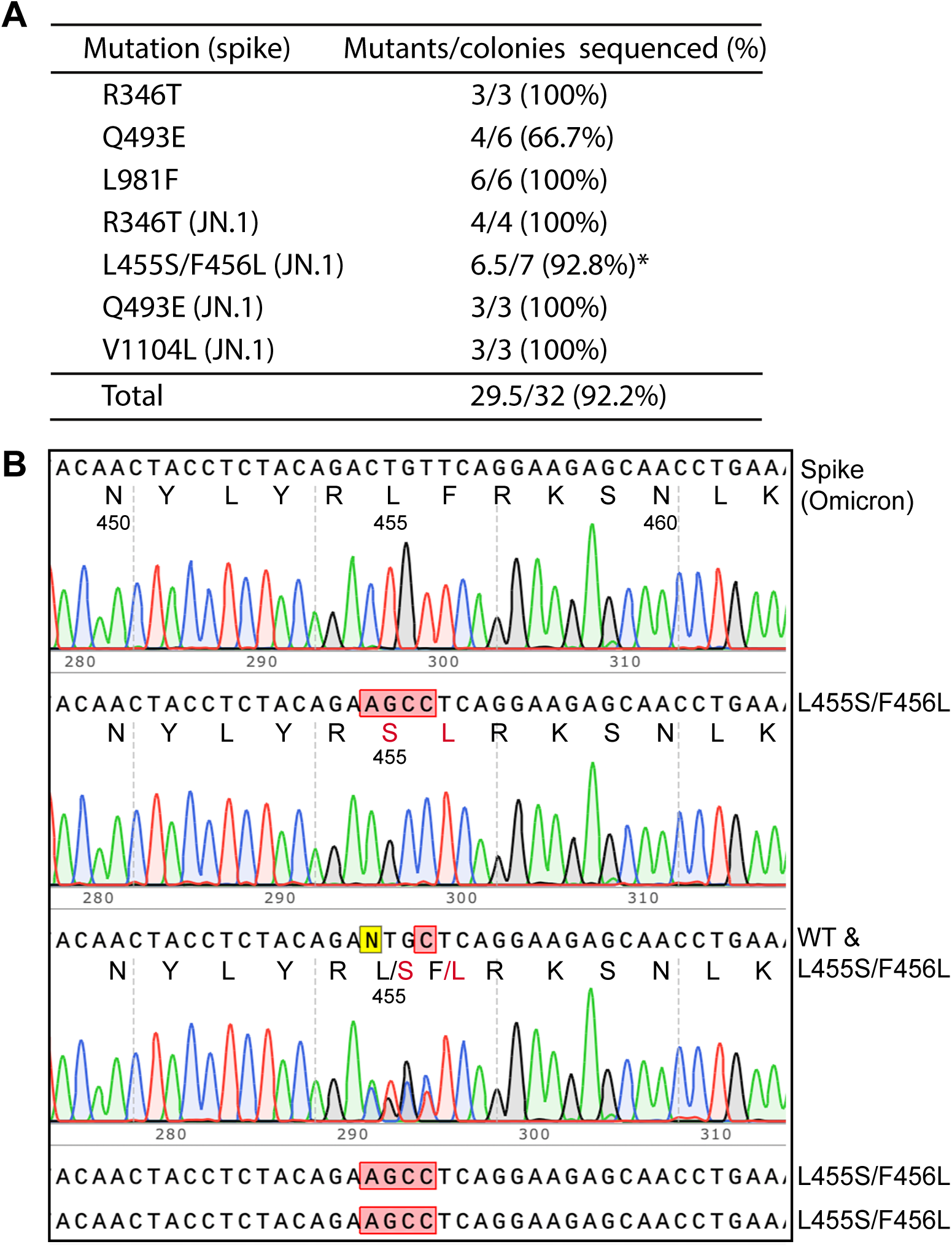
Rapid construction of SARS-CoV-2 spike mutants by P3a site-directed mutagenesis. A. Efficiency for generating seven different spike mutants. The first three were generated with the coding sequence for the D614G spike protein, whereas the last four were generated with the coding sequence for the Omicron spike protein. The asterisk indicates that among the 7 colonies analyzed, the L455S/F456L mutant plasmid is predominant. We sequenced plasmids from 8 more colonies; among them, five are for the L455S/F456L mutant plasmid, one carries the double mutation but with deletion of an A just upstream from the mutation, another is wild-type and the 8th could not be sequenced. We reasoned that primer imbalance might have caused the problem (due to different quality), so we adjusted their ratio so that the concentration of the primer for the F456L mutation is four times of that for the L455S/F456L double mutation. The mutagenesis experiment was repeated under the adjusted primer condition and 4 of the resulting colonies were sequenced. One of them carries the F456L mutant plasmid, another is for L455S/F456L and the remaining two are wild-type. Notably, perhaps due to 3′->5′ exonuclease activity of the polymerase used, mutagenesis was also successful even though one primer for the R436R substitution has a 2-bp mismatch at the 3*′*-end with the expression plasmid for the Omicron spike protein. B. Sequence chromatograms of representative plasmids sequenced for engineering SARS-CoV-2 spike mutants. The top chromatogram is the wild-type as the control and the remaining four are candidates from P3a site-directed mutagenesis. The sequencing results show that three are the L455S/F456L mutant and one contains a mixture of the wild-type and this mutant.

### P3a cassette mutagenesis to introduce scarless large deletion

Completely complementary primers (Fig. 1A) can be used to introduce small deletions (such as removal of a few base pairs), but it is quite challenging to introduce large deletions (such as those from a few hundred base pairs to several kb). This is a common problem for all known mutagenesis methods. One approach is to amplify by PCR and subclone a DNA fragment with deletion, but this is time-consuming. The fragment functions as a cassette, so such an approach is also known as ‘cassette’ mutagenesis. The ‘handshaking’ feature of the primer pair with 3′-overhangs inspired us to postulate that such a primer pair can be leveraged for introducing large deletions (Fig. 1D). Related to this, the ideal efficiency of 100% in generating the two mutants del364-394 and del364-427 of the HAT domain of KAT6B (deletion of 93 and 182 bp in the respective coding sequences; Fig. S2B) made us reason that P3a mutagenesis can indeed be used to delete large DNA fragments. To explore this concept further, we tested it with additional vectors, some of which are much larger than that for the HAT domain of KAT6B (Fig. S2B). We first assessed the concept for generating two N-terminal truncation mutants of BRPF1 (Fig. 4A). Shown in Fig. 4B-C is the primer design strategy to delete the 393-bp fragment encoding the N-terminal 131 residues. A similar strategy was used to design the primer pair for engineering dN204, the truncation mutant lack of the N-terminal 204 residues (Fig. 4A). All six colonies analyzed by Sanger sequencing carry the expected deletions (Fig. 4A), thereby reaching the ideal efficiency of 100%. We also generated a mutant with an internal deletion of 11 amino acids (d594-604) and obtained two corrected colonies out of three, yielding an efficiency of 66.7% (Fig. 4A).

**Fig. 4.**
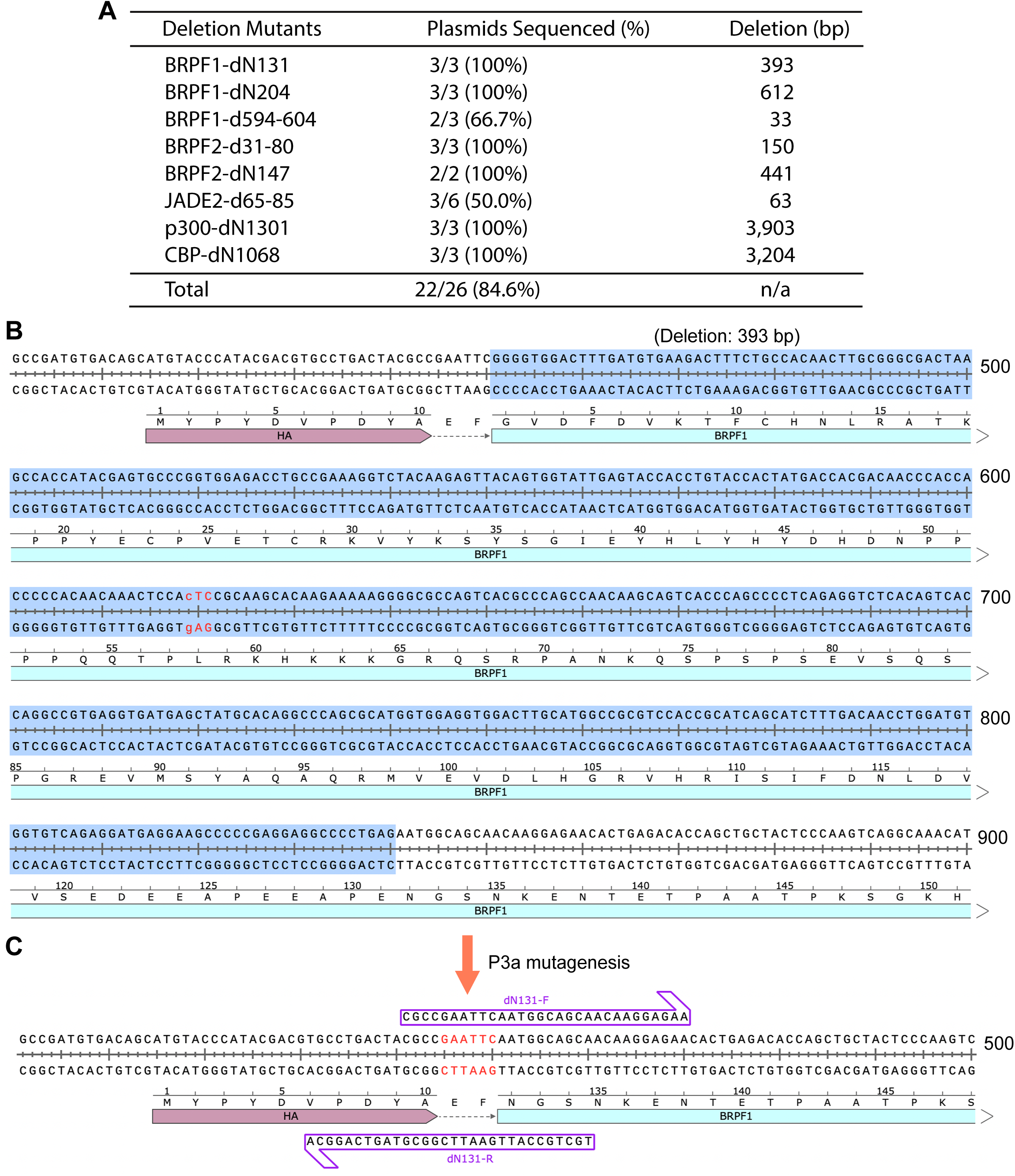
Seamless deletion of small and large DNA fragments by P3a cassette mutagenesis. A. Efficiency of generating three BRPF1 deletion mutants and five other truncation or internal deletion mutants. The primer design for the other mutants is similar to what is illustrated for the dN131 mutant of BRPF1 in panels B-C. B. Sequence of the region encoding the N-terminal 131 residues of human BRPF1. The 393-bp DNA fragment to be deleted is highlighted in blue. C. Sequence of the deletion mutant and the two partially complementary primers (dN131-F and dN131-R) that were used for P3a cassette mutagenesis. The notation ‘dN131’ refers to the deletion of the N-terminal 131 residues.

Success with these three BRPF1 deletion mutants led us to apply a similar strategy to generate larger deletion mutants of a paralog, BRPF2, and a related protein, JADE2, as we needed such mutants to investigate their interaction with KAT7. As shown in Fig. 4A, for two BRPF2 deletion mutants and one JADE2 deletion mutant, we analyzed 11 colonies and found that 8 were mutants, so the efficiency was 8/11 (72.3%). We next tested the strategy with the expression vectors for p300 and CBP, which are 12.8 and 13.4 kb in size, respectively. We aimed to engineer two mutants to delete the coding sequences for the N-terminal 1031 residues of p300 and the N-terminal 1068 residues of CBP (deletion of 3.1 and 3.2 kb in the respective coding sequences; Fig. 4A). As shown in Fig. 4A, all six colonies analyzed by Sanger sequencing have the expected mutations, thereby reaching the ideal efficiency of 100%. These results attest to the notion that P3a cassette mutagenesis is highly efficient for introducing small or large deletion mutants.

### Seamless epitope tagging and untagging by P3a cassette mutagenesis

The high efficiency in engineering deletions via P3a mutagenesis led us to investigate the possibility to use this strategy to introduce insertions (Fig. 1E). We first tested a mammalian expression vector for human p300, which does not possess a FLAG epitope for affinity purification on M2 agarose, so we sought to insert such a tag at the N-terminus of p300 (Fig. 5A). In the design, we also deleted 92 bp between the CMV promoter and the p300 coding sequence (Fig. 5B-D). As shown in Fig. 5D, all three colonies analyzed by Sanger sequencing carried the deletion of the 92-bp fragment and the insertion of the FLAG-coding sequence, thereby reaching the ideal efficiency of 100%. This supports that P3a cassette mutagenesis is highly efficient for epitope tagging.

**Fig. 5.**
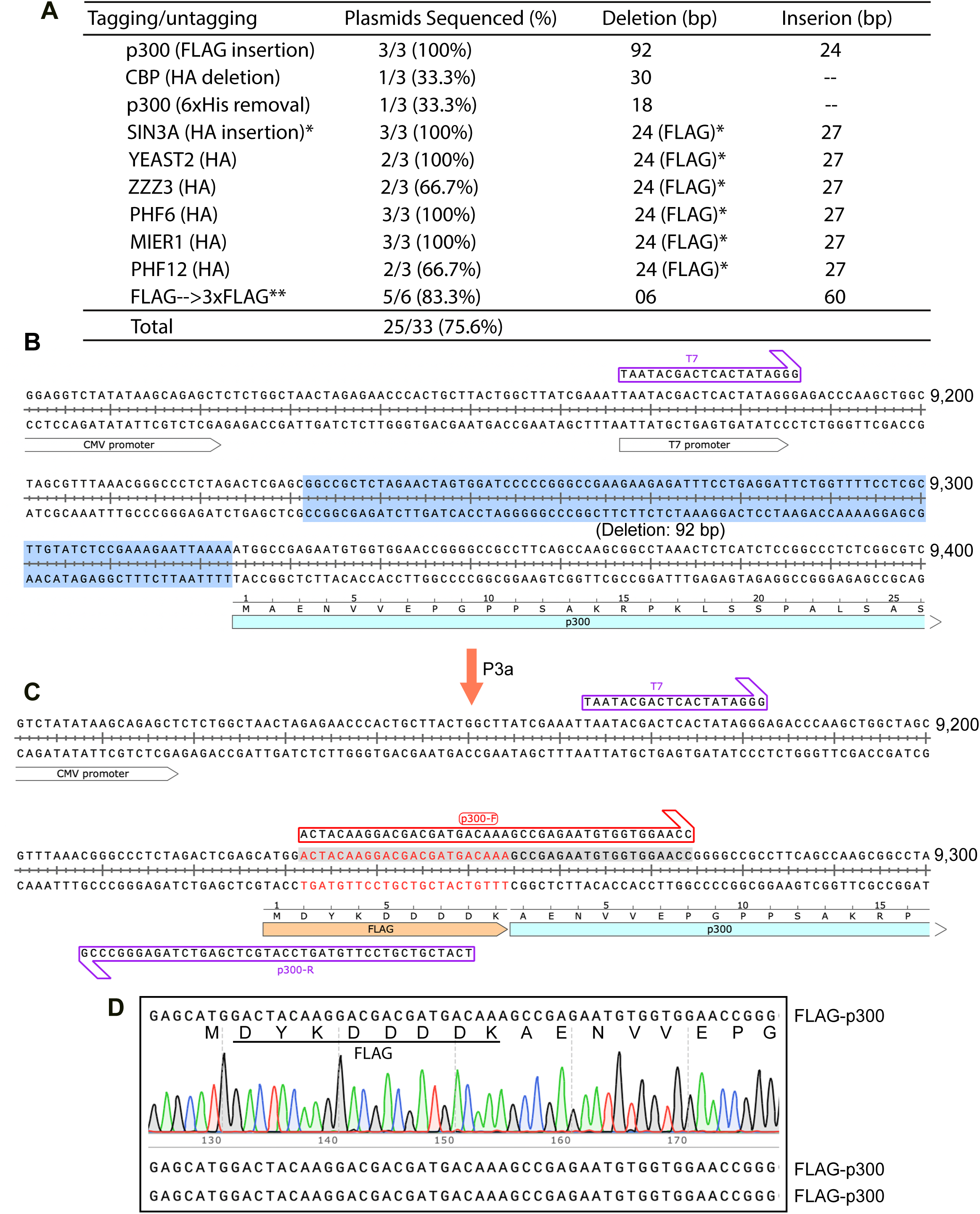
Scarless epitope tagging and untagging via P3a cassette mutagenesis. A. Efficiency of epitope tagging into mammalian expression vectors for 8 epigenetic regulators and epitope removal for two expression vectors (deletion of an HA tag from the CBP expression plasmid encoding both FLAG and HA tags, and removal of a 6xHis tag from the p300 expression vector). A single asterisk denotes deletion of the coding sequence of a FLAG tag, where the double asterisks refer to insertion of the coding sequence for tandem FLAG tags into a vector that already carries one such tag. B. DNA sequence encoding the N-terminal region of human p300. A 92-bp fragment to be deleted during P3a mutagenesis for introducing the coding sequence for a FLAG tag is highlighted in blue. C. Sequence of the resulting mutant and the two partially complementary primers (p300-F and p300-R) that were used for P3a site-directed mutagenesis. The two primers were designed to delete the 92-bp fragment (highlighted in panel B) and insert the coding sequence for the FLAG tag. D. Sanger sequence chromatogram of one mutant plasmid analyzed for the FLAG-coding region (top). Sequences of two other mutant candidates are at the bottom two lines. The results indicate that all three plasmids carry the FLAG coding sequence properly inserted.

We also tested the strategy to modify additional vectors. One such expression vector for CBP encodes two epitope tags, FLAG and HA, initially designed in another lab for tandem affinity purification. The HA tag interferes with detection of CBP partners in co-immunoprecipitation experiments, as they are also expressed as HA-tagged fusion proteins. We thus sought to remove the HA tag from the CBP expression vector using the strategy illustrated in Fig. S3A-B. As shown in Fig. 5A, one of three colonies analyzed by Sanger sequencing carries the expected deletion. Similarly, the p300 expression vector has a 6xHis tag, which may cause non-specific interaction. We employed a strategy analogous to that depicted in Fig. S3A-B to remove this tag. One of three colonies analyzed by sequencing carries the expected deletion. These results support that P3a cassette mutagenesis is efficient for epitope untagging.

We next investigated epitope tag conversion, i.e. involving replacement of DNA fragments (just as ‘cassettes’). Related to this, we had purchased expression vectors for multiple epigenetic regulators, including SIN3A, YEATS2, ZZZ3, MIER1, PHF6 and PH12 (Fig. 5A). These vectors were initially designed for expression of FLAG-tagged proteins, but for co-immunoprecipitation with partners that are also expressed as FLAG-tagged proteins, it is necessary to replace the FLAG tag with another tag, such as the HA tag. For the SIN3A vector, we utilized a strategy depicted in Fig. S4A-B to achieve tag conversion. As shown in Figs. 5A and S4C, all three colonies analyzed by Sanger sequencing were the expected mutants, thereby reaching the ideal efficiency of 100%. Similar strategies were employed to replace the FLAG tag on the expression vectors for YEATS2, ZZZ3, MIER1, PHF6 and PH12. As shown in Figs. 5A and S4D, 12 of 15 colonies analyzed by Sanger sequencing were the expected deletion mutants, thereby reaching the efficiency of 75%. Using this strategy (Fig. S3C), we converted a FLAG expression vector to a 3xFLAG vector. Notably, the ‘handshaking’ region is only 18 bp, allowing 60-bp insertion with a pair of primers only 58 nucleotides long (Figs. 5A & S3C). This is based on the general strategy illustrated in Fig. 1E. As shown in Fig. 5A, 5 of 6 colonies analyzed by Sanger sequencing harbor the expected plasmids, thereby reaching an efficiency of 83.3%. The overall efficiency for manipulating epitope tags on the 10 vectors is 75.6% (Fig. 5A), attesting to the high efficiency of P3a cassette mutagenesis for epitope tagging and untagging.

### Scarless engineering of restriction and LoxP sites by P3a site-specific mutagenesis

High efficiency in deletion of large DNA fragments encoding different epigenetic regulators by P3a mutagenesis (Fig. 4) led us to consider the use of this method for plasmid engineering. For example, the mammalian expression vector for p300 is almost 12.8 kb (Fig. S5A) and its preparation yield from bacterial cultures is much lower than other plasmids of similar size. To enhance the yield, we considered to delete some unnecessary sequences. Related to this, the Fi origin that was designed decades ago for preparation of phagmid forms is obsolete and no longer needed (Fig. S5A). Moreover, the Neo gene marker and its promoters are not necessary either. Thus, P3a mutagenesis was carried out to delete 1,928-bp fragment encompassing the F1 origin and the Neo elements. The resulting plasmid was expected to be 10.7 kb (Fig. S5B). The two primers were designed as illustrated in Fig. S5C. As shown in Fig. S5D, 2 of 4 colonies analyzed by Sanger sequencing are the expected deletion mutants. The preparation yield of the mutant plasmids was still similar to the parental plasmid, indicating that sequences other than the deleted region are responsible for the low yield. Among the remaining two colonies, one was the original plasmid and the other contained a much larger deletion. Thus, the mutagenesis efficiency was 50%. Due to its reduced size, the resulting plasmid with the 1,928-bp deletion should allow easy insertion of other sequences, such as those of KAT6A and KAT6B for engineering related leukemic fusion proteins [43].

We next sought to engineer simultaneous double deletions for plasmid optimization. Like the p300 expression plasmid, many plasmids in use now contain unnecessary sequences that can be deleted for optimization, such as the F1 origin mentioned above. Thus, we sought to optimize a commonly used 5.4-kb vector in our laboratory. This vector, known as pAW51 (Fig. S6A), was initially derived from a pcDNA3.1 vector and encodes an HA tag. pAW51 contains two regions that are no longer needed. One of them is a 430-bp fragment containing the F1 origin, and the other is a 390-bp fragment that encompasses a CAP-binding site and a Lac operator. To optimize this plasmid, we sought to delete these two fragments. The resulting plasmid is smaller (4.6 kb, Fig. S6B). Two pairs of primers were designed (Fig. S6C-D). These four primers were used in the same mutagenesis reaction to introduce both deletions simultaneously. In the first trial, we used 15 ng of the parental plasmid as the PCR template (condition A, Fig. S6E). We analyzed plasmids from 9 colonies by restriction digestion (Fig. S6E). Two plasmids possess the 390-bp deletion and 4 contain the 430-bp deletion. Among these 6, one carries both deletions. As the efficiency to obtain the mutant plasmid with both deletions is low, 1/9 (11.1%), we tested two additional conditions by using 5 ng of the parental plasmid as the PCR template (conditions B-C, Fig. S6E). For each of these two conditions, we analyzed plasmids from 8 colonies by restriction digestion (Fig. S6E). For both conditions, 7 colonies contain one of the two deletions. For condition B, three colonies harbor the mutant plasmid with both deletions (Fig. S6E). For condition C, none of the 8 colonies possess the mutant plasmid with both deletions (Fig. S6E). For these three trials, the overall efficiency to obtain the mutant plasmid with both deletions is only 4/25 (16%). These results suggest that while P3a mutagenesis is efficient in engineering single-site deletions, it is feasible for P3a mutagenesis to introduce two mutations simultaneously. Thus, alternative approaches are needed to enhance the efficiency. One possibility is to engineer two deletions with sequential mutagenesis reactions.

For plasmid engineering, it is often necessary to introduce or remove restriction sites for subcloning. Related to this, we needed to remove one of two NheI site that pAW51 harbors, to make the remaining site unique. The site to be deleted is located upstream from the coding sequence for the HA tag (Fig. S6A). This was achieved easily with the primers NheI-F and NheI-R, at the mutagenesis efficiency of 100% (Fig. S7A). Into the resulting vector (pAW51a), we then introduced an XhoI site and an NheI site with another pair of primers, XhoI-F and XhoI-R (Fig. S7B). Again, this was easily carried out, at the mutagenesis efficiency of 100% (Fig. S7A). Similarly, we introduced an HindIII site into pCL36, a vector almost the same as pAW51, but with a different poly-linker downstream from the coding sequence for the HA tag (Fig. S7A). As shown at the top of Fig. S8, we designed one pair of primers to introduce the HindIII site following two different reading frames downstream from the HA coding sequence. We analyzed 4 colonies by Sanger sequencing. All of them were the expected mutants, with 3 for one reading frame and the 4th for the other reading frame (Fig. S8), reaching the ideal efficiency of 100%. For these cases, restriction digestion could be used as the initial screening for the resulting colonies, so the results could be obtained within 3 days after primers were synthesized. The candidates from the initial screens could then be validated by Sanger sequencing.

To produce mammalian expression vectors for human SIN3B, USP10 and histone H4, we obtained their coding sequences from the ORFeome Collaboration Collection [48]. We initially tried PCR to amplify the coding sequences for subcloning, but due to some unknown technical reasons, the amplification failed. To overcome this technical problem, we sought to utilize P3a mutagenesis to introduce necessary restriction sites for subcloning. Into the vectors for SIN3B and USP10, we introduced an HindIII site upstream from their coding sequences. The resulting plasmids were analyzed by restriction digestion, and Sanger sequencing was performed for verification. The analyses revealed that the efficiency was 44.4% and 75% for introducing two HindIII sites into the SIN3B and USP10 expression vectors, respectively. The large size of the SIN3B insert (∼3.6 kb) might have contributed to the lower efficiency. For the histone H4 expression vector, we needed to introduce XhoI and BamHI sites upstream and downstream from the coding sequence, respectively. We designed two pairs of primers and used both pairs in one mutagenesis reaction. We analyzed 4 colonies by digestion. Two had the correct digestion pattern and were verified by Sanger sequencing, so the efficiency to obtain both restriction sites on the same vector is 2/4 (50%), supporting the feasibility of P3a mutagenesis to introduce two distant mutations simultaneously.

We also utilized P3a mutagenesis to insert a LoxP site, which is widely used for inducing Cre-dependent gene deletion in cells and animals. Typically, such sites are introduced by standard subcloning techniques relying on restriction sites, but a seamless mutagenesis method to insert such sites at ease is advantageous. As a proof-of-concept experiment, we engineered a 34-bp LoxP site on a BRPF1 expression vector (Fig. S9A). Out of 6 colonies analyzed, all contained the insertion and all of them possessed the correct insertion, leading to the ideal efficiency of 6/6 (100%), supporting that P3a mutagenesis is efficient to introduce LoxP sites. Therefore, together with the results on deletion of large fragments from the p300 expression vector (Fig. S5) and pAW51 (Fig. S6), the above experiments on introduction and deletion of restriction sites (Figs S7-S8) and LoxP sites (Fig. S9) indicate that P3a mutagenesis is efficient and valuable for seamless plasmid engineering.

### Deletion and insertion by P3 cassette mutagenesis

Previously, we systematically analyzed P3 site-directed mutagenesis [18,19], but it remains unclear whether the method is applicable to introduction of deletion and insertion mutations using the new strategies (Fig. 1D-E) as described above for P3a mutagenesis. Thus, it is important to compare the two methods in this regard. For these, we utilized P3 site-directed mutagenesis and Pfu_Ultra for generating 6 deletion mutants (Fig. S9B) and one insertion mutant (insertion of a LoxP site, Fig. S9A-B). For generation the 6 deletion mutants, the efficiency ranged from 33.3% to 85.7%, with the average level at 56-57% (Fig. S9B). This is much lower than what we obtained with the P3a method (Fig. 4). As for introduction the LoxP site, we analyzed plasmids from 6 colonies and four contained the correct inserts, with the remaining two possessing unwanted insertions (Fig. S9A-B). Thus, the efficiency was 4/6 (66.7%), lower than the 100% efficiency what we obtained with the P3a method (Fig. S9A). Thus, P3 mutagenesis was reliable for introducing deletion and insertion but less efficient than the P3a method. In addition to higher efficiency than the original P3 method, P3a mutagenesis offers four other advantages: 1) PCR length of about 2 hours, whereas P3 mutagenesis takes 4-10 hours dependent on the vector size and the choice of the extension temperature at 68°C or 72°C; 2) many more colonies (Fig. S9C); 3) fewer competent cells needed to be transformed; and 4) lower frequency of unwanted mutations. Therefore, P3a mutagenesis is much more advantageous than the P3 method itself.

*Engineering different* g*enome-editing vectors by P3a site-specific mutagenesis*

CRISPR has become a standard tool for genome editing [49]. An important aspect is to generate guide RNA-expressing vectors and modify Cas9 and other editing enzymes for biomedical research and eventual clinical evaluation. An efficient and seamless mutagenesis method should facilitate this process. As such, we applied P3a mutagenesis for modifying Cas9 expression vectors. As a proof-of-concept experiment, we first deleted a 5-kb fragment encompassing the Cas9 sequence and its promoter from pX459 and an pX330-derivative (Fig. 6A). pX330 and pX459 are two widely used Cas9 expression vectors due to their description in one of the first CRISPR-Cas9 papers [50]. For each mutagenesis reaction, 4 colonies were analyzed and all of them possessed the expected deletion, leading to the ideal efficiency of 100% (Fig. 6A). During the process, we noticed two 44-bp repeats shared by the sgRNA scaffold and its downstream region (Figs. 6B & S10A). This downstream repeat hinders P3a mutagenesis of the scaffold and insertion of gRNA-coding sequence upstream from the scaffold, so the second repeat was deleted along with its downstream TTT, leading to removal of a 47-bp fragment (Fig. S10B). For the mutagenesis reaction, plasmids from 4 colonies were analyzed and all of them possessed the expected deletion, leading to the ideal efficiency of 100% (Fig. 6A). Based on this vector, we engineered an sgRNA to edit the codon for R318 of human BRPF1. From the mutagenesis reaction, plasmids from 4 colonies were analyzed and three of them possessed the expected deletion, leading to an efficiency of 75% (Figs. 6A & S10C). Optimization of sgRNA is important for enhancing the editing efficiency. One possibility is to replace the premature termination signal TTTT within the scaffold with TTTC (Fig. S10C) [51]. Accordingly, the downstream AAAA needs to be replaced with GAAA [51]. Another possibility is to introduce a stable hairpin, referred to as Gold t-lock, involving a 11-bp insertion [52]. We carried out both mutagenesis reactions and analyzed three colonies from each (Fig. 6A). All of them possessed plasmids with the designed insertion, so the mutagenesis efficiency was 100%.

**Fig. 6.**
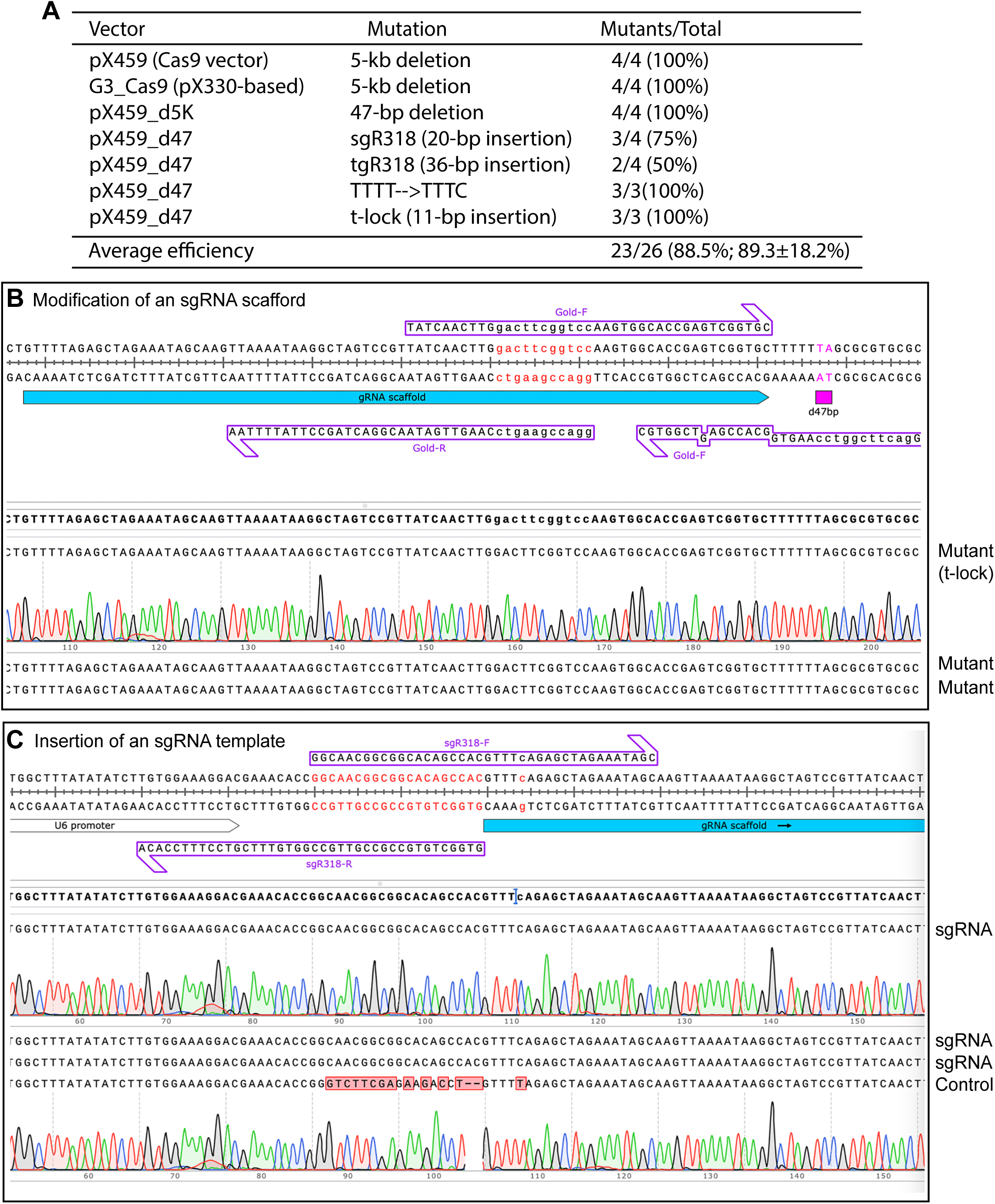
P3a site-directed mutagenesis of Cas9 and guide RNA vectors for genome editing. A. Summary of the efficiency from P3a site-directed mutagenesis to engineer deletion, insertion and point mutations in two different Cas9 vectors (pX459 and a related plasmid) and two guide RNA vectors (pX459_d5K and pX459_d47). tgR318 refers to a vector expressing a tigRNA [53] targeting the coding sequence around R318 of human BRPF1. B. The Cas9 expression vectors pX330, pX459 and some of their derivatives possess two 44-bp repeats in the sgRNA scaffold and its downstream region. The repeats are problematic for correct annealing of PCR primers designed to mutate the scaffold itself or insert sgRNA-coding sequences just upstream from the scaffold. C. Sequence chromatograms of four representative plasmids sequenced for engineering an sgRNA for editing the codon of R318 of human BRPF1. The two primers (sgR318-F and sgR318-R) were designed to express the sgRNA site from U6 promoter. The first three clones contained the insertion, whereas the 4^th^ was the vector itself, resulting in an efficiency of 3/4 (75%).

A recent study discovered and developed a new genome-editing TIGR-Tas system that is independent of any PAM sequences, so it is advantageous over CRISPR-Cas9 and any other known genome-editing systems [53]. The new system involves a new type of guide RNA, referred to as tigRNA. To introduce this system for editing the codon for R318 of human BRPF1, we engineered a tigRNA vector based the sequence encompassing the codon for R318 of human BRPF1. From the mutagenesis reaction, plasmids from 4 colonies were analyzed and two of them possessed the expected deletion, leading to an efficiency of 50% (Fig. 6A). Perhaps due to the primer quality or because of unique sequence at and round the deletion region, this efficiency was lower than the other mutations. Overall, the average mutagenesis efficiency reaches ∼90% for engineering the 7 genome-editing vectors (Fig. 6A).

### Restriction site-independent insertion or replacement of gene fragments via P3a mutagenesis

In the above experiments, we utilized regular oligonucleotides as primers. As such primers are limited to 60 nucleotides, the insert size is restricted to 60 bp. This is because the two primers need 20 nucleotides for each of the two arms to anneal to the template and also 20 nucleotides for the overlapping region of the two primers. To circumvent this limitation, we sought to utilize Ultramer primers, which can be up to 200 nucleotides, so the insert size can be increased to 340 bp (Fig. 7A). To implement this strategy, we tested three cases. In the first one, we sought to repair an incomplete ORF clone that we had for human HAT1. The clone encodes residues 85-419, with the N-terminal 84 residues missing (Fig. S11A). To complete the ORF, we designed two Ultramers, HAT1-uF and -uR (Fig. S11B), for P3a cassette mutagenesis. Plasmids from a dozen bacterial colonies were analyzed by restriction digestion and 6 of them were found to carry the expected insertion of 0.25 kb. Three of them were sequenced and all were correct (Fig. S11C). Thus, the overall efficiency was 6/12 (50%).

**Fig. 7.**
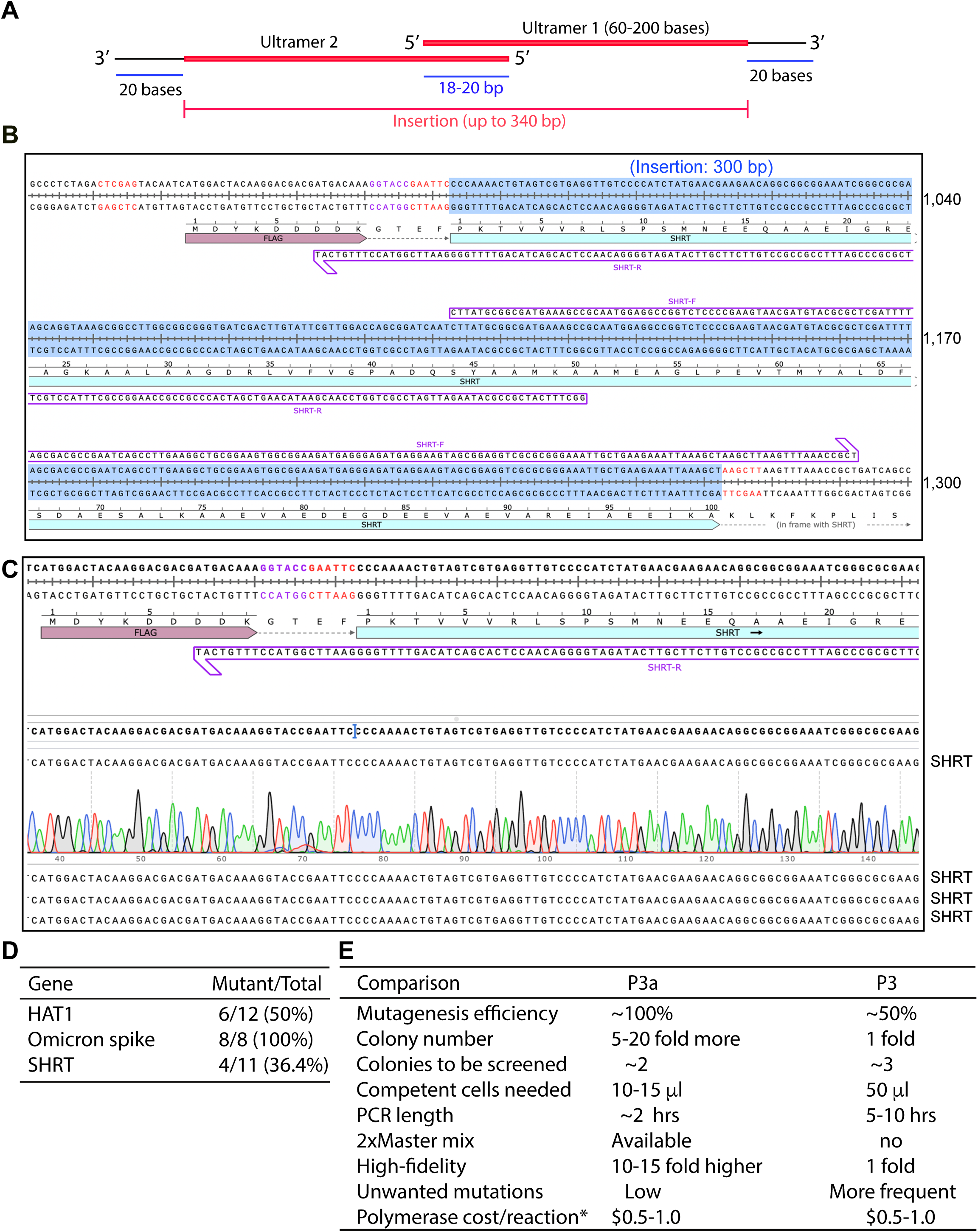
P3a cassette mutagenesis directed by Ultramer primer pairs. A. Cartoon showing a pair of Ultramer oligo primers for introducing insertion up to 0.36 kb. The current up limit of an Ultramer is 0.2 kb. B. Insertion of the 300-bp coding sequence for SHRT (an artificial protein that neutralize a snake toxin) into a mammalian expression vector C. Sequence chromatograms of four representative plasmids sequenced for engineering the coding sequence for SHRT. Eleven colonies were analyzed by restriction digestion and 4 were found to contain the correct insertion (as shown here), resulting in an efficiency of 4/11 (36.4%). D. Efficiency of P3a cassette mutagenesis to insert gene fragments via Ultramer primers. E. Summary of different parameters for direct comparison of P3 and P3a mutagenesis methods, with the latter being much faster, more convenient and efficient.

In the second case, we sought to clone the ORF of an artificial protein known as SHRT, which was AI-designed to bind and neutralize a snake venom toxin [54]. SHRT is composed of 100 residues and its ORF is 0.3 kb. We utilized two Ultramers, SHRT-F and SHRT-R (Fig. 7B). From the mutagenesis reaction, plasmids from 11 bacterial colonies were analyzed by restriction digestion and 4 of them were found to carry the expected insertion of 0.3 kb. All of them were sequenced and two were correct (Fig. 7C). For the remaining two, one possessed a T insertion and the other carried a G deletion. The synthesis yield for these two Ultramers was lower than those for HAT1, and one-nucleotide insertion or deletion is expected to be frequent for long primers like Ultramers. It is very likely that these two mutations were due to the quality of the Ultramers. Thus, the overall efficiency was 4/11 (36.4%). As mutations from incorrect synthesis are expected to be frequent for long primers like Ultramers, we sought to correct the T deletion and G deletion by regular P3a site-specific mutagenesis. We analyzed 4 colonies each, and three or four of them were correct, supporting the usefulness of the method to correct unwanted mutations resulting from DNA synthesis.

In the last case, we replaced the coding sequence for the receptor-binding domain of the spike protein encoded by the D614G SARS-CoV19 variant with the counterpart of a new Omicron subvariant, which possesses 13 extra mutations compared to the original D614G variant (Fig. S12A) *[6,47]*. For this, a crossover strategy was used (Fig. S12A), with the two Ultramers Omi430-F and Omi430-R (Fig. S12A-B). From the mutagenesis reaction, plasmids from 8 bacterial colonies were analyzed by sequencing and all of them were found to carry the expected mutations (Fig. S12C), resulting in the ideal mutagenesis efficiency of 100%. Thus, as summarized in Fig. 7D, the above three cases based on Ultramers illustrate that P3a cassette mutagenesis is efficient for inserting and replacing gene fragments in a seamless manner.

## Discussion

Site-directed mutagenesis is a fundamental technique for biomedical research, used to analyze disease-associated germline or somatic mutations [2–4], assess the impact of viral genetic alterations during evolution [5,6], generate specific mutants of natural proteins [7] and engineer artificial proteins from natural backbones [8,9] or AI-assisted *de novo* design [11–13]. An innovative strategy involves the use of partially complementary primers with 3′-protruding ends (Fig. 1B) [18,20,21]. These primers anneal to nicked regions of newly synthesized DNA strands, enabling their use as new templates for PCR amplification (Fig. S1A). In the previous study [18], we evaluated this method with a dozen mammalian expression plasmids. While it is generally reliable for many cases, the efficiency was close to 50% and can be improved further. Moreover, other issues, such as 1) unwanted mutations at the primer sites, 2) relatively low fidelity of Pfu polymerase and 3) slow synthesis rate of this enzyme (Fig. 7E), require further improvement of the method [18]. In the current study, we have developed a new approach by replacing Pfu and its derivatives with Q5 and Platinum SuperFi II high-fidelity DNA polymerases, both of which elevate the efficiency close to the ideal level of 100%, speed up PCR amplification and eliminate unwanted mutations at or close to the primer sites (Figs 1-3 & 7E). Typically, the new method requires fewer PCR cycles than what was described previously [18]. We have evaluated the new method with over ten mammalian expression vectors encoding various epigenetic regulators (Figs 1-2, 4 & S1-S2) and two vectors for the spike protein of SARS-CoV-2 (Fig. 3).

In light of the unique handshaking feature of the primers with 3′-overhangs, we have extended the method for seamless cassette mutagenesis to engineer deletion (Fig. 1D) and insertion (Fig. 1E), as exemplified by introduction of large deletion, up to 5 kb (Figs. 4A & 6), epitope tagging/untagging (Figs. 5 & S3-S4), plasmid optimization (Figs S5-S8) and modification of genome-editing vectors (Figs 6 & S10). The largest plasmid that we tested is the 13.4-kb CBP expression vector (Figs. 4A & S2D). For seamless plasmid minimization, we tested the method with four plasmids (Figs. S5-S7 & 6A). But it should be applicable to many other plasmids, especially for those (such as viral vectors for preclinical research and gene therapy), where the size limit is a critical bottleneck. Notably, for deletion and insertion, the P3a method was more efficient than P3 mutagenesis (Fig. S9B).

For scarless epitope tagging and untagging, we tested the method with four tags, including FLAG, 3xFLAG, HA and 6xHis tags (Fig. 5A), but the method should be applicable to many other tags, such as a modified tandem affinity purification tag [55] and a novel degron [56]. Notably, the size of insertion is limited to the two primers that can be synthesized chemically. The largest insertion that we have made was the conversion of a FLAG tag to a 3xFLAG tag, where the inserted sequence is 60 bp (Fig. 5A). Moreover, as an Ultramer DNA primer can be up to 200 nucleotides long, the insertion could be up to 0.4 kb in size (2×200 bp, according to the strategy shown in Fig. 7A). We have successfully implemented this strategy (Figs. 7 & S11-S12). This is not something that the strategy depicted in the previous study could achieve [21]. Moreover, sequential mutagenesis should help circumvent the size limit (Fig. S13E). As shown for introducing epitope tags (Figs S3-S4), this method should also allow efficient and seamless insertion of oligonucleotide duplexes for engineering gRNA and shRNA vectors. Similarly, this method shall also be useful for seamless insertion of LoxP and other recombination sites for construction of targeting vectors for homologous recombination and conditional knockouts. Theoretically, the method can also be used for seamless insertion of the coding sequences for nuclear localization and export signals.

The strategy utilizing primer pairs with two 3′-overhangs was initially implemented by Zheng, Baumann & Reymond [20] and subsequently refined by Liu & Naismith [21]. As the first step towards the ideal goal to reach the mutagenesis efficiency of 100%, we have recently optimized the method and systematically tested it with over a dozen plasmids, ranging in 7.0-13.4 kb in size [18]. The success rate varies, with an average efficiency of about 50% for many plasmids [18], thereby leaving space for further improvement. As the second step towards the aforementioned ideal goal, the current study fills this void and raises the mutagenesis efficiency to or near 100%, making the new method a routine choice for site-directed mutagenesis in molecular biology experiments. Even more importantly, this current study provides an effective remedy to address the challenges to insert or delete large fragments, up to 5 kb (Fig. 7). Related to this, only small deletion and insertion (up to 18 nucleotides) were tested in the previous study (Fig. S13A) [21]. Even for such small deletion, the primer size ranges from 38-46 nucleotides [21]. In comparison, we used primers ∼30 nucleotides long (Fig. 1D), thereby reducing costs and minimizing the chance of unwanted mutations from primers. Different from that used in the previous study (Fig. S13A) [21], the overlapping region of the primer pair can be a new sequence, e.g. a codon optimized one (Fig. S13B). Moreover, our insertion strategy (Fig. 1E) is completely different from (and also superior to) the one used in this previous study, thereby allowing much larger insertion (Fig. S13C-D) [21]. In comparison, from the previous studies [18,20,21], it was unclear if the methods work with large deletion and insertion. To our knowledge, no effective methods have been reported in the literature. The current study offers a simple method for this purpose.

In light of the unique ‘handshaking’ feature of primer pairs with 3′ overhangs, the novel strategy depicted in (Fig. 1E) was successfully to replace DNA fragments (Fig. 5A). Conceptually, deletion and insertion can be considered as two special cases where the respective sequences to be inserted and deleted are zero bp (Fig. S13F; also see the legend to Fig. 1E). Thus, the insertion strategy is also applicable to deletion and replacement of DNA fragments, as exemplified in Fig. 5. Notably, the unique ‘handshaking’ feature of primer pairs with 3′ overhangs is lacking in QuickChange primer pairs (Fig. 1A), thereby limiting this previous method to small deletion, insertion or replacement. Thus, the P3 and P3a methods are also advantageous over the QuickChange strategy in this regard.

One common challenge with site-directed mutagenesis is occurrence of unwanted mutations due to three sources: (1) impurity in primers, (2) unwanted recombination at the mutation or primer sites, and (3) misincorporation during synthesis mediated by DNA polymerase. Related to source (3), Q5 and SuperFi II DNA polymerases are high-fidelity enzymes with misincorporation rates 10-15-fold lower than PfuUltra that we used in the previous study [18] and 30-35-fold lower than regular Pfu. Related to this, results from over 100 Sanger sequencing reactions performed for the current study have not yielded any such mutations away from the primer sites. Related to source (2), the previous study frequently uncovered unwanted mutations at the region at or close to the primer sites when PfuUltra was used, perhaps due to its strand-replacement activity or unwanted synthesis when reaching the 5′ end of an annealed primer [18]. By contrast, we have found only two such cases in the current study (Figs. S4D & S5D). One of them (Fig. S4D) is right at a primer site, so it could also be due to source (1). Instead of unwanted recombination at the primer sites or misincorporation during PCR, impurity in primers is a major source of unwanted mutations, so it is important to verify the sequence at the primer sites. The results from the current study and those from the previous study [18] also indicate that Q5 and SuperFi II polymerases replicate much more faithfully than Pfu and its derivatives when DNA synthesis reaches the primer sites.

The current study focuses on engineering single-site mutations. For pAW51 (Fig. S6) and the histone H4 vector (Fig. S7), we tested the feasibility to engineer double mutations simultaneously. The results show that the dual-site mutagenesis efficiency was 50% for the histone H4 vector (Fig. S7A) and only 16% for pAW51(Fig. S6). Theoretically, PCR amplification between the forward primer for site 1 and reverse primer for site 2, as well as PCR amplification between the reverse primer for site 1 and forward primer for site 2, becomes dominant and may overshadow the expected two PCR amplifications from two pairs of primers at sites 1 and 2 for P3a mutagenesis. This problem is expected to get worse for multi-site mutagenesis. This is an intrinsic flaw of all mutagenesis methods based on primer pairs, including the QuickChange^™^ and P3 site-directed mutagenesis methods (Fig. 1A-B). One potential solution is to carry out sequential mutagenesis, but this is time-consuming, especially for multi-site mutagenesis where the site number is 5 or more. Therefore, alternative approaches are needed to generate multi-site mutations fast and efficiently.

In summary, we have developed a fast, economical and highly efficient site-specific mutagenesis method based on Q5 and SuperFi II high-fidelity DNA polymerases, along with primer pairs with 3′-overhangs (Fig. 1C-E). We have evaluated this method systematically by engineering >100 mutations on >20 mammalian expression vectors, up to 13.4 kb in size. The results indicate that the method is superior to those based on Pfu and its derivatives [18,20,21], by elevating the efficiency to or close to 100%, speeding up PCR amplification and reducing unwanted mutations at the primer sites. Even more importantly, the high efficiency and the ‘handshaking’ feature of the primer pairs with 3′-protruding ends have led us to extend the method for seamless cassette mutagenesis, which allows highly efficient epitope tagging or untagging, deletion of small or large DNA fragments, optimization of plasmid constructs and insertion of oligonucleotide duplexes with 3′-overhangs. Given its versatility, P3a site-specific and cassette mutagenesis is expected to be of wide use for biomedical research.

## ACKNOWLEDGEMENT

We thank Ms Paulina Varela-Castillo and Arezousadat Razavi for critically reading the manuscript. This work was supported by funds from Canadian Institutes of Health Research (CIHR), Natural Sciences and Engineering Research Council of Canada (NSERC) and Compute Canada (to X.J.Y.).

## Materials and Methods

### Plasmid vectors

The mammalian expression vector for wild-type *Xenopus* histone H3 with its C-terminus fused to a FLAG tag was modified from a pcDNA3.1 derivative containing the coding sequence for this epitope tag. A bacterial expression vector encoding *Xenopus* histone H3 was generously provided by Dr. Karolin Luger [57]. Mammalian expression vectors for HA-tagged BRPF1, BRPF2 (also known as BRD1) and BRPF3 were described earlier [31,41,58]. The mammalian expression vector for FLAG-KAT6A was kindly provided by Dr. Issay Kitabayashi [59]. The mammalian expression vector for the catalytic domain of KAT6A was described previously [31,58]. The mammalian expression vectors for FLAG-KAT6B and its catalytic domain have been reported [31,58,60]. The expression plasmids for untagged p300 and FLAG/HA-tagged mouse CBP were obtained from Addgene (Cat. 23252 and 32908, respectively). Mammalian expression vectors for untagged D614G and Omicron spike proteins of SARS-CoV-2 were purchased from SinoBiological (Cat. VG40589-UT and VG40835-UT, respectively). Vectors containing the coding sequences for human SIN3B (BC172412), histone H4 (EU446978) and USP10 (DQ892553), initially produced as a part of the ORFeome Collaboration Collection [48], were obtained from the Platform for Cellular Perturbation at McGill University, Montreal, Canada. The mammalian expression vectors for human SIN3A, YEATS2, ZZZ3, MIER1, PHF6 and PHF12 were purchased from GeneScript. They were expressed as fusion proteins with a FLAG tag fused to the C-termini, upon transfection into mammalian cells. pX459_HypaCas9 and G3_Cas9 (eSpCas9(1.1)_No_FLAG_ATP1A1_G3 [61], a pX330-derived vector) were purchased from Addgene (#108294 and #86611, respectively).

### Primers for site-directed mutagenesis

Primers were designed with the aid of SnapGene software package (version 7.2.1). They were synthesized at Integrated DNA Technologies (IDT) as small scale, standard and desalted DNA oligos, without further purification. Primers used to generate *Xenopus* histone H3 mutants for expression in mammalian cells as fusion proteins with a FLAG tag at the C-terminus were as listed below, where the mutated nucleotides are shown in lowercase: K9R-F, CCGCCCGGAgATCCACCGGAGGGAAGGCTCCCCGC; K9R, CCGGTGGATcTCCGGGCGGTCTGCTTAGTACGAGC; K14R-F, CCGGAGGGAgGGCTCCCCGCAAGCAGCTGGCCACC; K14R-R, CGGGGAGCCcTCCCTCCGGTGGATTTCCGGGCGGT; K18R-F CTCCCCGCAgGCAGCTGGCCACCAAGGCAGCCAGG, K18R-R, GCCAGCTGCcTGCGGGGAGCCTTCCCTCCGGTGGA; K23R-F, TGGCCACCAgGGCAGCCAGGAAGAGCGCTCCGGCC; K23R-R, CTGGCTGCCcTGGTGGCCAGCTGCTTGCGGGGAGC; K27M-F, CAGCCAGGAtGAGCGCTCCGGCCACAGGCGGAGTC; 27R-R, GGAGCGCTCcTCCTGGCTGCCTTGGTGGCCAGCTG; G34R-F, GCCACAGGCaGAGTCAAGAAACCTCACCGTTACCG; G34A-R, TTCTTGACTgCGCCTGTGGCCGGAGCGCTCTTCCT, K36M-F, GCGGAGTCAtGAAACCTCACCGTTACCGGCCCGGC; and K36R-R TGAGGTTTCcTGACTCCGCCTGTGGCCGGAGCGCT.

The primers for engineering plasmids to express deletion mutants of human BRPF1 were dN131-F, CGCCGAATTCaATGGCAGCAACAAGGAGAA; dN131-R, TGCTGCCATtGAATTCGGCGTAGTCAGGCA; dN204-F, CGCCGAATTCtCTGCAGAGGAGCTGGACGA; dN204-R, CCTCTGCAGaGAATTCGGCGTAGTCAGGCA; d594-604-F, CCTGGCAGCGGGTGGAATTGATCCGCAAGCG; d594-604-R, GATCAATTCCACCCGCTGCCAGGACTTGAGCT; Y215A-F, GAAGTAGAGgcTGACATGGACGAGGAGGACT; Y215A-R TCCATGTCAgcCTCTACTTCCTCGTCCAGCT; W224A-F GACTACATCgcGCTGGATATCATGAATGAGC; W224A-R ATATCCAGCgcGATGTAGTCCTCCTCGTCCA; F245A-F CAGGAGATCgcTGAGTACCTAATGGACCGAC; F245A-R AGGTACTCAgcGATCTCCTGCGGGATGGGAC; R381A-F CCACCAGCTgcCTGGAAGCTCACCTGCTACA; R381A-R AGCTTCCAGgcAGCTGGTGGGATGTGCTCAA; K390A-F TACATTTGCgcACAACGGGGCTCAGGGGCCT; K390A-R CCCCGTTGTgcGCAAATGTAGCAGGTGAGCT; R392A-F TGCAAACAAgcGGGCTCAGGGGCCTGCATCC; and R392A-R CCTGAGCCCgcTTGTTTGCAAATGTAGCAGG, where the mutation is denoted with lowercase letters.

The primers to engineer the expression plasmids for human BRD1 mutants were as follows: G38R-F, CAAGCTCAAaGGATGGTAGAGATAGAAATT; G38R-R CTACCATCCtTTGAGCTTGAGCGTAGGTCA; I53A-F, ACAGGATCAGTgcTTTTGATCCCCTGGAGATCA; I53A-R, GGGATCAAAAgcACTGATCCTGTGCAAGCGCC; L57A-F, TTTTTGATCCCgctGAGATCATATTGGAAGATGA; L57A-R, AATATGATCTCagcGGGATCAAAAATACTGATCC; I59A-F, TCCCCTGGAGgcCATATTGGAAGATGACCTCA; I59A-R, CTTCCAATATGgcCTCCAGGGGATCAAAAATA; I60A-F, CCTGGAGATCgcATTGGAAGATGACCTCACTG; I60A-R, ATCTTCCAATgcGATCTCCAGGGGATCAAAAA; L61A-F, GGAGATCATAgcGGAAGATGACCTCACTGC; L61A-R, GTCATCTTCCgcTATGATCTCCAGGGGATCAA; P275A-F, GTGCTGTGCgCCAACAAGGGTGGTGCCTTC; P275A-R, CCTTGTTGGcGCACAGCACACAGTCGGCGG; H275P-F, TGCTGTGCCCCAACAAGGGTGGTGCCTTCA; H275P-R, CCCTTGTTGGGGCACAGCACACAGTCGGCG; d31-80-F, GCGAGAAACggAGCGGCCTCCTGTCTGCTT; d31-80-R1, GAGGCCGCTccGTTTCTCGCGTAGGTGAGTGTTTA; dN147-F, CGCCGAATTCGAGGAACTGGACAACGAGGT; and dN147-R, CCAGTTCCTCGAATTCGGCGTAGTCAGGCA.

The primers for engineering plasmids toFLAG-tagged human KAT6A mutants were as follows: K604R-F, TTGACCACAgAACCCTCTATTACGATGTGG; K604R-R, TAGAGGGTTcTGTGGTCAAGAAACAACTTT; K350Kfs-F, ACGGTATCAAAGGTCCCTTCAGCAAAGTTCGAACTGGCCCTGGAA; K350Kfs-R TGAAGGGACCTTTGATACCGTGTTTTGTTTCTTTAACCTGTTTTT; R469*-F, AATGAGGAGtGACTTTTTGGGAGCCAGGAA; L668F-F CAGTTATTTcTTATCAAAGCGTGAAGGCCAAGCAG; L668F-R, CTTTGATAAgAAATAACTGAAATCGATGAGAAACC; P682Q-F, CAGAGAAACaGTTATCTGATCTGGGTCGTCTTTCC; P682L-R, TCAGATAACaGTTTCTCTGGAGACCCTGCTTGGCC; Q743R-F, GTAGTGACCgATTTGTGATTATCCGCCGGGAAAAA; Q743R-R, ATCACAAATcGGTCACTACGGAAGTCCAGCATTCG; R740H-F, TGGACTTCCaTAGTGACCAATTTGTGATTATCCGC; R740L-R, TGGTCACTAaGGAAGTCCAGCATTCGTAGGTGGTG; R469*-R, CAAAAAGTCaCTCCTCATTTTCTTGTTTGC; G814*-F, ATCAGTGTGtGAAAGTCTGTGTCTCATGAG; G814*-R, CAGACTTTCaCACACTGATCTCTAATTCTC; L1061*-F, ATGCCAAGATaAGAACCCACGTTTGAGATCGATGAAGAAGAGGAG; L1061*-R, CGTGGGTTCTtATCTTGGCATTGGCCTCTCGGAGTCAGAATCTTC; E1393*-F, GCTGGGTCTtAGGACGACCACGAAGAAGAC; and E1393*-R, GGTCGTCCTaAGACCCAGCCATCTGCTCTG.

The primers for engineering mammalian vectors to express human KAT6B mutants as a FLAG-tagged fusion proteins were as follows: F551L-F, GTTGGATACCTCTCTAAGGAAAAGCTTTGC; F551L-R, CCTTAGAGAGGTATCCAACCAGATGACAGC; Q863H-F, CATGCCCCAcCACCAAAGGCAAGGATTTGGACGGT; Q863H-R, CCTTTGGTGgTGGGGCATGATCATTATGCAGGAGA; E884G-F, CTAGAAGAGgAGGCCAAGCAGGGTCTCCTGAAAAG; E884G-R, GCTTGGCCTcCTCTTCTAGAAAGCAAATAGCTGAA; S889C-F, AAGCAGGGTgTCCTGAAAAGCCTCTCTCCGATCTG; S889C-R, TTTTCAGGAcACCCTGCTTGGCCTTCTCTTCTAGA; D896N-F, CCTCTCTCCaATCTGGGCCGTCTCTCCTACCTGGC; D896N-R, GGCCCAGATtGGAGAGAGGCTTTTCAGGAGACCCT; R899H-F, ATCTGGGCCaTCTCTCCTACCTGGCATATTGGAAG; R899H-R, TAGGAGAGAtGGCCCAGATCGGAGAGAGGCTTTTC; E912K-F, GTCATCTTGaAGTATCTCTACCACCACCATGAGAG; E912K-R, AGAGATACTtCAAGATGACGCTCTTCCAATATGCC; d364-394-F, ATTCGGATCCGTCGTTACTGAAGAGGATTT; d364-394-R, CAGTAACGACGGATCCGAATTCGGTACCTT; d364-427-F, ATTCGGATCCCCTTCTGTGATTGAATTTGG; and d364-427-R, TCACAGAAGGGGATCCGAATTCGGTACCTT.

The primers for engineering plasmids to express human KAT7 mutants were as follows: C185S-F, ATATGAAGTcTCCTACACCAGGCTGTAACTCT; C185S-R, GGTGTAGGAgACTTCATATTGAAGTTGTAGCT; C209S-F, TCTCAGGATcCCCACTGTATCATAACCTCTCA; C209S-R, TACAGTGGGgATCCTGAGATGGAGAAATGTCT; K432R-F, CTGGACCACcgtACATTATATTATGATGTGGAGCCCT; K432R-R, AATATAATGTacgGTGGTCCAGAAAcAGTTTGGCCAA; I537A-F, TTTTCAAGGCgcAGAGATTTCTATCAAAGAAA; I537A-R, AGAAATCTCTgcGCCTTGAAAATTATGCAGGT; I539A-F, AGGCAAAGAGgcTTCTATCAAAGAAATCAGTC; I539A-R, TTTGATAGAAgcCTCTTTGCCTTGAAAATTAT; I541A-F, AAGAGATTTCTgcCAAAGAAATCAGTCAGGAGAC; I541A-R, GATTTCTTTGgcAGAAATCTCTTTGCCTTGAAA; V574A-F, AACACCTAGcTTTAAAGAGACAGGACCTGA; V574A-R, CTCTTTAAAgCTAGGTGTTTTCCCTTCCAG; K576A-F, ACCTAGTTTTAgccAGACAGGACCTGATTGATGA; and K576A-R, AGGTCCTGTCTggcTAAAACTAGGTGTTTTCCCT.

The primers for engineering plasmids to express human KAT8 mutants were as follows: E87D-F, GAGGGCCGAGAtGAATTCTATGTACACTACGT; E87Q-R, ATAGAATTCCTgTCGGCCCTCCTGGTCGTTCA; V91I-F, GAATTCTATaTACACTACGTGGGCTTTAAC; V91I-R, GGAAGTCCCaCAGGATCTCTAGCAGCACCC; R136C-F, GCAGCCTGAGtGCAAGATCACTCGCAACCAA, R136H-R, AGTGATCTTGtGCTCAGGCTGCTCTGCGAGC; R140H-F, AAGATCACTCaCAACCAAAAGCGCAAGCATG; R140G-R, TTTTGGTTGCcAGTGATCTTGCGCTCAGGCT; D190G-F, ACGAAATTGgTGCCTGGTATTTCTCACCAT; D190G-R, CATCGGGAACTACGAAATTGgTGCCTGGTA; R224H-F, AAGAGCTACCaCTTCCACTTGGGTCAGTGCC; R224G-R, AAGTGGAAGCcGTAGCTCTTCTCATATTTCA; Y241C-F, AAGAGATCTgCCGCAAGAGCAACATCTCCG; Y241C-R, TCTTGCGGcAGATCTCTTTCCCGGGtGGCT; P321H-F, TGACCTTGCaCCCCTACCAACGCCGCGGCTA; P322L-R, GCGTTGGTAGaGGGGCAAGGTCAGGATGCAG; R374W-F, GAGATCCTGtGGGACTTCCGGGGCACACTG; and R374W-R, AGGGCCGAGAGGAATTCTATaTACACTACG.

The primers for engineering plasmids to express human p300 mutants were as follows: p300-F, ACTACAAGGACGACGATGACAAAGCCGAGAATGTGGTGGAACC; p300-R, TCATCGTCGTCCTTGTAGTCCATGCTCGAGTCTAGAGGGCCCG; dN1031-F, ACTACAAGGACGACGATGACAAAAGTACTTCAGCTACCCAGTC; d6His-F, CTAGACATATAGTGATACTAAGCTTAAGT; d6His-R, GTATCACTATATGTCTAGTGTACTCTGTG; d2K-F, GGTATCCCCTCATGTCTGTATACCGTCGA; d2K-R, CAGACATGAGGGGATACCCCCTAGAGCCC; C1204S-F, TGTGAGAAGTcTTTCAATGAGATCCAAGGG ; C1204R-R, TCATTGAAACgCTTCTCACAGAAATGATACC; E1242K-F, ACTGGATCCTaAACTGTTTGTTGAATGTACA; E1242A-R; ACAAACAGTgCAGGATCCAGTGTGTCATTT; D1384N-F, TATGGCTCTaACTGCCCTCCACCCAACCAG; D1384H-R, GAGGGCAGTgAGAGCCATACTCTTGAACAT; D1399Y-F, ATCTTACCTCtATAGTGTTCATTTCTTCCGT; D1399N-F, GCCTGGGCTaCTGGGTGGCTGCAGCCTGCTY1414C-F, CTGCAGTCTgTCATGAAATCCTAATTGGAT; Y1414C-R, ATGAACACTATtGAGGTAAGATATGTATACT; W1466C-F, GCAGGAATGcTACAAgAAAATGCTTGACAAGGCTGT; Y1467N-R, ATACACAAGgCATAATCCTCACAGACAGTA; S1726*-F, GCCACCCAGtAGCCCAGGCGATTCTCGCCG; S1726*-R, ATTTCATGAcAGACTGCAGTCCTCAAGCAT; R1830*-F, ATGCTTCGCtAGGAGGATGGCCAGCATGCA; R1830*-R, CATCCTCCTaGCGAAGCATTTGGGCCTGCT; Q1845*-F GGGCAGCAAtAGGGCCTCCCTTCCCCCACT; and Q1845*-R, GGAGGCCCTaTTGCTGCCCAACCACACCAG.

The primers for engineering plasmids to express mouse CBP mutants were as follows:dN1068-F, CGACGACAAAGACACAGCCTCACAATCAAC; dN1068-R, AGGCTGTGTCTTTGTCGTCGTCGTCCTTGT; R1446C-F CCGCTGCCTCtgtACAGCTGTTTACCATGAGAT; R1446H-R, AACAGCTGTatgGAGGCAGCGGGGCCGGAAGA; Y1503D-F, ACAGGAGTGGgACAAGAAGATGCTGGACAAG; Y1503H-R, GGATTCCTGtGGGCTCTaGGACTGTGGCTCACCCT; R1868*-F, GCTCATGCGCtaaCGAATGGCAACCATGAACAC; R1868*-R, TTGCCATTCGttaGCGCATGAGCTGAGCCTGCT; Q1881*-F, GTGCCTCAGtAGAGTTTGCCTTCTCCTACC; and Q1881*-R, GCAAACTCTaCTGAGGCACATTGCGGGTGT.

The primers for generating vectors to express mutants of the D614G and Omicron spike proteins were previously described [25]. The backward primers for engineering an HA tag fused to the C-terminus of MIER1, SIN3A, PHF6, PHF12, ZZZ3 and YEATS2 were HA-MIER1 (TAGTCAGGCACGTCGTATGGGtagTCATCTGTGTTTTCAAGTT), HA-SIN3A (TAGTCAGGCACGTCGTATGGGtaaGGGGCTTTGAATACTGTGC), HA-PHF6 (TAGTCAGGCACGTCGTATGGGtaGTTTCCATTAAGTTGCTGCT), HA-PHF12 (TAGTCAGGCACGTCGTATGGGtaaGGAACAGAGTTGGAGCGCA), HA-ZZZ3 (TAGTCAGGCACGTCGTATGGGtaTCTGTTTGCTGGAAAGTAGT), HA-YEATS2 (TAGTCAGGCACGTCGTATGGGtaCTGGTCCTCATTCAATATTC), respectively. The forward primer was the same: HA-F4, CCATACGACGTGCCTGACTACGCCTGATAAACCCGCTGATCAGC. The original plasmids, purchased from Genescript, were based on a pCDNA3.1 derivative encoding a FLAG tag fused to the C-terminus of a target protein. To convert the coding sequence for a FLAG tag on pAW48 (a pCDNA3.1 derivative) to that for a 3xFLAG tag, the two primers used were as follows: 3Flag-F, TGATAAGGGCAGCGATTATAAGGATGACGACGATAAGGAATTCCACCACACTGGACTA; and 3Flag-R, TAATCGCTGCCCTTATCATCATCGTCTTTATAATCTGCACCTTTGTCATCGTCGTCCT.

Primers for modifying pAW51, a pcDNA3.1 derivative encoding an HA tag, were as follows: NheI-F, AGCTGGCTAcgGTTTAAACGGGCCCTCTAGA; NheI-R, CGTTTAAACcgTAGCCAGCTTGGGTCTCCCT; XhoI-F, CTCGAGGATATCGCTAgcCACCACACTGGACTAGTGGA; and XhoI-R, GGCTAGCGATATCCTCGAGGAATTCGGCGTAGTCAGGCA. Primers for introducing an HindIII site into pCL36, another pcDNA3.1 derivative encoding an HA tag, were 36H-F GGAATTCTGaagcttTCCAGCACAGTGGCGGCCGC and 36H1-R, TGTGCTGGAaagctttCAGAATTCCGGCGTAGTCAG. These two primers led to two constructs with different reading frames downstream from the coding sequence for the HA tag.

Primers to introduce an HindIII site into the 5′ region of the encoding sequence for human Sin3B were SIN3B-HF, CCCACCATGaagcttGCGCACGCTGGCGGTGGCAGC and SIN3B-HR, AGCGTGCGCaagcttCATGGTGGGCCTTGACGGCC. Primers to introduce an XhoI site and a BamHI site flanking the encoding sequence of human histone H4 were HisH4-XF1, AGCAGGCTCgagCACCATGTCTGGTCGCGGCA; HisH4-XR1, GACATGGTGctcGAGCCTGCTTgTTTGTACAAAG; HisH4-BF1, CTTCGGCGGggatccTTGGACCCAGCTTTCTTGTA; and HisH4-BR1, TGGGTCCAAggatccCCGCCGAAGCCGTAAAGAGT.

The primers to introduce a LoxP site were LoxP-F, CGTATAATGTATGCTATACGAAGTTATCTAGCaccatgtacccatac; and LoxP-R, CGTATAGCATACATTATACGAAGTTATCCAGCTTGGGTCTCCCTATA. The primers used for engineering genome-editing vectors were: d47bp-F, GTGCTTTTTTAGCGCGTGCGCCAATTCTGC; d47bp-R, CGCACGCGCTAAAAAAGCACCGACTCGGTG; d5K-F TGGCTCTAGAGAATTCCTAGAGCTCGCTGA; d5K-R, CTAGGAATTCTCTAGAGCCATTTGTCTGCA; TTTC-F, ACCTGTTTcAGAGCTAGAAATAGCAAGTTgAAATAAGGCTAGTCCGTTAT; TTTC-R, GCCTTATTTcAACTTGCTATTTCTAGCTCTgAAACAGGTCTTCTCGAAGA; Gold-F, TATCAACTTGgacttcggtccAAGTGGCACCGAGTCGGTGC; Gold-R, ggaccgaagtcCAAGTTGATAACGGACTAGCCTTATTTTAA; tgR318-F, CTGTGtgaaacccaCAACGGCGGtgcgTTTTTTGTTTTAGAGCTAGA; tgR318-R, CGTTGtgggtttcaCACAGCCACtggctCGGTGTTTCGTCCTTTCCAC; sgR318-F, GGCAACGGCGGCACAGCCACgtttcagagctagaaatagc; sgR318-R GTGGCTGTGCCGCCGTTGCCggtgtttcgtcctttccaca.

Ultramer primers were ordered from IDT at the PAGE-purified grade: HAT1-uF, gGAAGATCTTGAAAATGACATTAGAACTTTCTTTCCTGAGTATACCCATCAACTCTTT GGtGATGATGAAACTGCTTTTGGTTACAAGGGTCTAAAGATCCTGTTATACTATATTG CTGGTAGCCTGTCAACAATGTTCCGTGTTGAATATGCATCT; HAT1-uR, TCATTTTCAAGATCTTCcGGgAAACGAACTAATTTTAGTTCAATTGCTGTGTTGGTGTT ACATTTGTACTCTGCCAGTTTCTTCTCCACTGCACTCTTATATTCTACCAAgAATTTCT CCATAGCACCAAATCCCGCGAATTCGGTACCTTTGTCAT; Omi430-F1, ACAGAGATTTACCAGGCTGGCAACAAGCCATGTAATGGAGTGGCCGGCTTCAACTG TTACTTTCCACTCgaATCCTATAGCTTCAGACCAACCTACGGAGTGGGCCACCAACCA TACAGGGTGGTG; Omi430-R1, AGCCTGGTAAATCTCTGTGCTGATGTCCCTCTCAAATGGTTTCAGGTTGCTCTTCCTG AggctTCTGTAGAGGTAGTTGTAGTTGCCGCTCACCTTGCTGTCCAGCTTGTTGCTGTT CCAGGCAAT; SHRT-F, CTTATGCGGCGATGAAAGCCGCAATGGAGGCCGGTCTCCCCGAAGTAACGATGTAC GCGCTCGATTTTAGCGACGCCGAATCAGCCTTGAAGGCTGCGGAAGTGGCGGAAGA TGAGGGAGATGAGGAAGTAGCGGAGGTCGCGCGGGAAATTGCTGAAGAAATTAAA GCTAAGCTTAAGTTTAAACCGCT; and SHRT-R GGCTTTCATCGCCGCATAAGATTGATCCGCTGGTCCAACGAATACAAGTCGATCACC CGCCGCCAAGGCCGCTTTACCTGCTTCGCGCCCGATTTCCGCCGCCTGTTCTTCGTTC ATAGATGGGGACAACCTCACGACTACAGTTTTGGGGAATTCGGTACCTTTGTCAT.

The primers to correct a G deletion and a T insertion in the SHRT coding sequence were: SHRT-iG-F, AAGGCTGCGGAAGTGGCGGAAGATGAGG; SHRT-iG-R, CCGCCACTTCCGCAGCCTTCAAGGCTGATT; SHRT-dT-F, AGGCCGGTCTCCCtGAAGTAACGATGTACGCGC; and SHRT-dT-R, CGTTACTTCaGGGAGACCGGCCTCCATTGCGGC.

### P3 mutagenesis via Q5 and SuperFi II high-fidelity DNA polymerases

For a mutagenesis reaction, the template plasmid needed to be isolated from DH5α or any other Dam^+^ *E. coli* hosts, as required for subsequent DpnI digestion. Primers were designed with the aid of the SnapGene software package and purchased from IDT as small-scale desalted oligos, without further purification. Their sequences were listed above. If primer sequences contain long stretches of A and/or G (typically more than 5), a silent mutation was introduced to break such stretches. This was also done with long stretches of T or C (typically more than 5). All oligonucleotides were used without polyacrylamide gel or HPLC purification, except for Ultramers. Upon receipt from IDT, autoclaved Nanopure water was used to dissolve lyophilized oligonucleotides to prepare 100 mM (i.e. 100 pmol/μl) stocks for further dilution to 10 mM (i.e. 10 pmol/μl) working solutions.

10 μl PCR reactions were set up in 0.2 ml 8-strip thin-wall PCR tubes (Diamed Cat. DIATEC420-1378), with each reaction containing 0.1-0.15 μl of plasmid DNA (0.1 μg/μl), 0.5 μl forward/reverse primer mixture (2.5 pmol/μl for each primer), 4.35-4.40 μl autoclaved Nanopure water and 5 μl 2x Q5 hot-start master mix (New England Biolabs, Cat. M0494S), 2x Platinum SuperFi II PCR master mix (Thermo Fisher Scientific, Cat. 12368010), 2x i7^®^ high-fidelity DNA polymerase master mix (Intact Genomics, Cat. 3257 or 3257S) or 2x KOD hot-start master mix (Sigma-Aldrich, Cat. 71842). The amplification was carried out in a Bio-Rad PCR T100 Thermal Cycler using the following parameters: 96°C for three minutes as the initial step to denature plasmid DNA, followed by 19-25 cycles to amplify the DNA. Each amplification cycle was composed of 93°C for 15 seconds as the denaturation step, 50-52°C for 20 seconds as the annealing step and 72°C for 3-5 minutes as the extension step, where the extension time was calculated according to the plasmid size (20-30 seconds per kb). The amplification cycle number varied slightly from one plasmid to another, with the initial number set at 19-20. Typically, for large plasmids such as those p300 and CBP, 25 cycles were used, while for the plasmids 6-7 kb in size, 19-20 cycles were sufficient. After amplification, an additional extension step of 72°C for 7.5-10 minutes was added to the PCR program. After that, the reaction was pre-programmed for short-term storage at 4°C. For DpnI digestion, 0.25 μl (20 units/μl; New England Biolabs, Cat. R0176) was added to each reaction mixture, which was then pipetted into a clean PCR tube for incubation at 37°C for 90 min in a Bio-Rad T100 Thermal Cycler. The clean PCR tube at this step helped eliminate contamination of undigested plasmid from the inner wall of the old PCR tube used for PCR amplification. The reagent per mutagenesis reaction was $0.5-1. To reduce the cost further, the reaction volume could be reduced to 5 μl, with 1-2.5 μl used for transformation (see below).

### Preparation and transformation of competent *E. coli* cells

DH5α competent cells were prepared and stored at -80°C or in a liquid nitrogen tank as described previously [18,19]. Briefly, 1.0-2.5 μl of the digested PCR mixture, preferred to be on ice, was gently mixed with 10-25 μl of DH5α competent cells on ice (just thaw from -80°C or a liquid nitrogen) in a 0.5 ml sterile Eppendorf tube (pre-chilled on ice) and incubated on ice for 30 min. Notably, even transformation of 1.0 μl of the digested PCR mixture into 10 μl of DH5α competent cells yielded sufficient colonies for analysis, which could save materials further. The remaining DpnI-digested PCR mixture was frozen at -20°C as the backup. After the 30 min-incubation on ice, the tube containing the DH5α competent cells and DpnI-digested PCR mixture was heat-shocked at 42°C for 45 seconds and immediately transferred back onto ice. 30-75 μl of cold SOC, NYZ^+^ or SON^+^ medium [18,19] was then added to the tube and after gentle tapping with a finger, the tube was incubated at a 37°C water bath for ∼30 min (without shaking, as the cells are fragile at this stage). Occasionally, the SOC, NYZ^+^ or SON^+^ medium was forgotten to add to some tubes, but the transformations still worked (albeit with fewer colonies), indicating that the medium helped but was not essential. Afterwards, the entire cell mixture was pipetted onto a LB-agar plate (containing the appropriate antibiotic as required for the plasmid to be mutated) as 4-5 drops, which were further spread out relatively evenly with the outside surface at the bottom half of an autoclaved 1.5 ml Eppendorf tube. After that, the plate was placed in a 37°C incubator for 22-24 h to allow large bacterial colonies to appear and grow.

To reduce the usage of bacterial plates by 50%, two reaction mixtures could be plated out on the two sides of a single LB-agar plate (as shown in Fig. S9D). For this, the back of the plate was marked with a straight dark line using a marker pen. The two mixtures were spotted onto the two sides of the plate while avoiding the area just flanking the dark central line. The two cell mixtures were spread out relatively evenly with the outside surfaces at the bottom half of two separate autoclaved 1.5 ml Eppendorf tubes. It was important to avoid spreading to the central part flanking the dark central line. For this, it was also necessary that the surface of the plate did not contain any visible liquid and that the cell mixture would not be more than 75 μl. It might also help avoid cross-contamination when the plate was allowed to stay and dry at room temperature for 15 min, but this was optional. The plate was then carefully transferred to a 37°C incubator and then flipped over gently, in the same direction of the central dark line, before putting it in the incubator for 22-24 h to allow large bacterial colonies to appear and grow. For a few mutagenesis reactions, this might not make a huge difference, but when dozens or hundreds of mutations were engineered, this practice did help cut the labor and reagent costs. This practice was mainly carried pout for the P3a but not P3 method, as transformation of 5 μl from a typical mutagenesis reaction mixture into 50 μl of DH5α competent cells was required for the latter [18].

### Analysis of plasmids from bacterial colonies

Plasmids were isolated from 2-3 bacterial colonies per mutation for Sanger sequencing as described previously [18,19]. The resulting DNA sequences were compared with that of the wild-type template for the presence of the desired mutations with the aid of the sequence alignment tools in the SnapGene software package. The corresponding chromatograms were manually inspected for sequencing quality. The resulting DNA sequences and their corresponding chromatograms were also examined manually for potential unwanted mutations.

### Plasmid sequencing and analysis

Plasmid isolation, sequencing and sequence analysis were performed as described above [18]. Briefly, after isolation with a Qiagen QIAprep® Spin Miniprep Kit, plasmids were sequenced at Genome Quebec Technological Service Center. For sequencing reactions, plasmids were denatured at 96°C, after which primer annealing was carried out at lower temperatures not exceeding 50°C to allow hybridization of sequencing primers. Samples from the sequencing reactions were analyzed on 96-capillary array DNA Sequencer 3730XL (Applied Biosystems). The resulting sequences and chromatograms were transferred to Snapgene 8.0.1 for alignment analysis and manual inspection.

### Estimation of misincorporation rate

DNA sequence alignment tools in the SnapGene software package were used to analyze Sanger DNA sequencing data. Sequence data quality was manually assessed from the corresponding sequencing chromatograms, and the aligned sequences were visually inspected to identify unwanted mutations. To calculate the misincorporation rate, the number of unwanted mutations was divided by the total number of the sequenced nucleotides whose corresponding sequencing chromatograms were inspected and determined to be of high quality.

## References

1. C. A. Hutchison, 3rd, S. Phillips, M. H. Edgell, S. Gillam, P. Jahnke and M. Smith (1978) Mutagenesis at a specific position in a DNA sequence. J Biol Chem 253, 6551–6560.

2. A. Henrie, S. E. Hemphill, N. Ruiz-Schultz, B. Cushman, M. T. DiStefano, D. Azzariti, S. M. Harrison, H. L. Rehm and K. Eilbeck (2018) ClinVar Miner: Demonstrating utility of a Web-based tool for viewing and filtering ClinVar data. Hum Mutat 39, 1051–1060.

3. E. Cerami, J. Gao, U. Dogrusoz, B. E. Gross, S. O. Sumer, B. A. Aksoy, A. Jacobsen, C. J. Byrne, M. L. Heuer, E. Larsson, Y. Antipin, B. Reva, A. P. Goldberg, C. Sander and N. Schultz (2012) The cBio cancer genomics portal: an open platform for exploring multidimensional cancer genomics data. Cancer Discov 2, 401–404.

4. J. G. Tate, S. Bamford, H. C. Jubb, Z. Sondka, D. M. Beare, N. Bindal, H. Boutselakis, C. G. Cole, C. Creatore, E. Dawson, P. Fish, B. Harsha, C. Hathaway, S. C. Jupe, C. Y. Kok, K. Noble, L. Ponting, C. C. Ramshaw, C. E. Rye, H. E. Speedy, R. Stefancsik, S. L. Thompson, S. Wang, S. Ward, P. J. Campbell and S. A. Forbes (2019) COSMIC: the Catalogue Of Somatic Mutations In Cancer. Nucleic Acids Res 47, D941–D947.

5. B. Hu, H. Guo, P. Zhou and Z. L. Shi (2021) Characteristics of SARS-CoV-2 and COVID-19. Nat Rev Microbiol 19, 141–154.

6. P. V. Markov, M. Ghafari, M. Beer, K. Lythgoe, P. Simmonds, N. I. Stilianakis and A. Katzourakis (2023) The evolution of SARS-CoV-2. Nat Rev Microbiol 21, 361–379.

7. D. G. L. Thean, H. Y. Chu, J. H. C. Fong, B. K. C. Chan, P. Zhou, C. C. S. Kwok, Y. M. Chan, S. Y. L. Mak, G. C. G. Choi, J. W. K. Ho, Z. Zheng and A. S. L. Wong (2022) Machine learning-coupled combinatorial mutagenesis enables resource-efficient engineering of CRISPR-Cas9 genome editor activities. Nat Commun 13, 2219.

8. G. D. Martyn, G. Veggiani, U. Kusebauch, S. R. Morrone, B. P. Yates, A. U. Singer, J. Tong, N. Manczyk, G. Gish, Z. Sun, I. Kurinov, F. Sicheri, M. F. Moran, R. L. Moritz and S. S. Sidhu (2022) Engineered SH2 Domains for Targeted Phosphoproteomics. ACS Chem Biol 17, 1472–1484.

9. Y. Bian, L. Li, M. Dong, X. Liu, T. Kaneko, K. Cheng, H. Liu, C. Voss, X. Cao, Y. Wang, D. Litchfield, M. Ye, S. S. Li and H. Zou (2016) Ultra-deep tyrosine phosphoproteomics enabled by a phosphotyrosine superbinder. Nat Chem Biol 12, 959–966.

10. R. A. Silverstein, N. Kim, A. S. Kroell, R. T. Walton, J. Delano, R. M. Butcher, M. Pacesa, B. K. Smith, K. A. Christie, L. L. Ha, R. J. Meis, A. B. Clark, A. D. Spinner, C. R. Lazzarotto, Y. Li, A. Matsubara, E. O. Urbina, G. A. Dahl, B. E. Correia, D. S. Marks, S. Q. Tsai, L. Pinello, S. S. De Ravin, Q. Liu and B. P. Kleinstiver (2025) Custom CRISPR-Cas9 PAM variants via scalable engineering and machine learning. Nature 10.1038/s41586-025-09021-y.

11. T. N. Starr, N. Czudnochowski, Z. Liu, F. Zatta, Y. J. Park, A. Addetia, D. Pinto, M. Beltramello, P. Hernandez, A. J. Greaney, R. Marzi, W. G. Glass, I. Zhang, A. S. Dingens, J. E. Bowen, M. A. Tortorici, A. C. Walls, J. A. Wojcechowskyj, A. De Marco, L. E. Rosen, J. Zhou, M. Montiel-Ruiz, H. Kaiser, J. R. Dillen, H. Tucker, J. Bassi, C. Silacci-Fregni, M. P. Housley, J. di Iulio, G. Lombardo, M. Agostini, N. Sprugasci, K. Culap, S. Jaconi, M. Meury, E. Dellota, Jr., R. Abdelnabi, S. C. Foo, E. Cameroni, S. Stumpf, T. I. Croll, J. C. Nix, C. Havenar-Daughton, L. Piccoli, F. Benigni, J. Neyts, A. Telenti, F. A. Lempp, M. S. Pizzuto, J. D. Chodera, C. M. Hebner, H. W. Virgin, S. P. J. Whelan, D. Veesler, D. Corti, J. D. Bloom and G. Snell (2021) SARS-CoV-2 RBD antibodies that maximize breadth and resistance to escape. Nature 597, 97–102.

12. S. L. Lisanza, J. M. Gershon, S. W. K. Tipps, J. N. Sims, L. Arnoldt, S. J. Hendel, M. K. Simma, G. Liu, M. Yase, H. Wu, C. D. Tharp, X. Li, A. Kang, E. Brackenbrough, A. K. Bera, S. Gerben, B. J. Wittmann, A. C. McShan and D. Baker (2024) Multistate and functional protein design using RoseTTAFold sequence space diffusion. Nat Biotechnol 10.1038/s41587-024-02395-w.

13. T. Kortemme (2024) De novo protein design-From new structures to programmable functions. Cell 187, 526–544.

14. T. A. Kunkel (1985) Rapid and efficient site-specific mutagenesis without phenotypic selection. Proc Natl Acad Sci U S A 82, 488–492.

15. M. P. Weiner, G. L. Costa, W. Schoettlin, J. Cline, E. Mathur and J. C. Bauer (1994) Site-directed mutagenesis of double-stranded DNA by the polymerase chain reaction. Gene 151, 119–123.

16. C. L. Fisher and G. K. Pei (1997) Modification of a PCR-based site-directed mutagenesis method. Biotechniques 23, 570–571, 574.

17. S. Li and M. F. Wilkinson (1997) Site-directed mutagenesis: a two-step method using PCR and DpnI. Biotechniques 23, 588–590.

18. N. Mousavi, Zhou, E., Razavi, A., Ebrahimi, E., Varela-Castillo, P. and Yang, X. J. (2025) P3 site-directed mutagenesis: An efficient method based on primer pairs with 3’-overhangs. J. Biol. Chem. 301, article # 108219 (also avaliable at bioRxiv: https://www.biorxiv.org/content/108210.101101/102024.108210.108218.615663v1082 12).

19. N. Mousavi, Zhou, E., Razavi, A., Ebrahimi, E., Varela-Castillo, P. and Yang, X. J. (2025) Efficient site-directed mutagenesis mediated by primer pairs with 3’-overhangs. Current Protcols in Protein Sciences 5, e70104.

20. L. Zheng, U. Baumann and J. L. Reymond (2004) An efficient one-step site-directed and site-saturation mutagenesis protocol. Nucleic Acids Res 32, e115.

21. H. Liu and J. H. Naismith (2008) An efficient one-step site-directed deletion, insertion, single and multiple-site plasmid mutagenesis protocol. BMC Biotechnol 8, 91.

22. F. Zeng, S. Zhang, Z. Hao, S. Duan, Y. Meng, P. Li, J. Dong and Y. Lin (2018) Efficient strategy for introducing large and multiple changes in plasmid DNA. Sci Rep 8, 1714.

23. K. Zhang, X. Yin, K. Shi, S. Zhang, J. Wang, S. Zhao, H. Deng, C. Zhang, Z. Wu, Y. Li, X. Zhou and W. Deng (2021) A high-efficiency method for site-directed mutagenesis of large plasmids based on large DNA fragment amplification and recombinational ligation. Sci Rep 11, 10454.

24. H. Jia, R. Couto-Rodriguez, S. Johnson, S. Medina, B. Novillo, P. Huynh, M. Kim, C. Cooper, M. Michalik, B. Siew, E. Turesson and J. A. Maupin-Furlow (2022) Highly efficient and simple SSPER and rrPCR approaches for the accurate site-directed mutagenesis of large and small plasmids. N Biotechnol 72, 22–28.

25. H. H. Hogrefe, C. J. Hansen, B. R. Scott and K. B. Nielson (2002) Archaeal dUTPase enhances PCR amplifications with archaeal DNA polymerases by preventing dUTP incorporation. Proc Natl Acad Sci U S A 99, 596–601.

26. Y. Wang, D. E. Prosen, L. Mei, J. C. Sullivan, M. Finney and P. B. Vander Horn (2004) A novel strategy to engineer DNA polymerases for enhanced processivity and improved performance in vitro. Nucleic Acids Res 32, 1197–1207.

27. B. D. Strahl and C. D. Allis (2000) The language of covalent histone modifications. Nature 403, 41–45.

28. B. A. Nacev, L. Feng, J. D. Bagert, A. E. Lemiesz, J. Gao, A. A. Soshnev, R. Kundra, N. Schultz, T. W. Muir and C. D. Allis (2019) The expanding landscape of ’oncohistone’ mutations in human cancers. Nature 567, 473–478.

29. K. A. Thompson, B. Wang, W. S. Argraves, F. G. Giancotti, D. P. Schranck and E. Ruoslahti (1994) BR140, a novel zinc-finger protein with homology to the TAF250 subunit of TFIID. Biochem. Biophys. Res. Commun. 198, 1143–1152.

30. X. J. Yang (2015) MOZ and MORF acetyltransferases: molecular interaction, animal development and human disease. Biochim Biophys Acta 1853, 1818–1826.

31. K. Yan, J. Rousseau, K. Machol, L. A. Cross, K. E. Agre, C. F. Gibson, A. Goverde, K. L. Engleman, H. Verdin, E. Baere, L. Potocki, D. Zhou, M. Cadieux-Dion, G. A. Bellus, M. D. Wagner, R. J. Hale, N. Esber, A. F. Riley, B. D. Solomon, M. T. Cho, H. Porter, B. C. Lanpher, A. M. Lewis, J. Savatt, I. Thiffault, B. Callewaert, P. M. Campeau and X. J. Yang (2020) Deficient histone H3 propionylation by BRPF1-KAT6 complexes in neurodevelopmental disorders and cancer. Sci. Adv. 6, eaax0021.

32. F. Huang, A. Paulson, A. Dutta, S. Venkatesh, M. Smolle, S. M. Abmayr and J. L. Workman (2014) Histone acetyltransferase Enok regulates oocyte polarization by promoting expression of the actin nucleation factor spire. Genes Dev 28, 2750–2763.

33. Y. Doyon, C. Cayrou, M. Ullah, A. J. Landry, V. Cote, W. Selleck, W. S. Lane, S. Tan, X. J. Yang and J. Cote (2006) ING tumor suppressors are critical regulators of chromatin acetylation required for genome expression and perpetuation. Mol. Cell 21, 51–64.

34. B. J. Klein, K. L. Cox, S. M. Jang, J. Cote, M. G. Poirier and T. G. Kutateladze (2020) Molecular Basis for the PZP Domain of BRPF1 Association with Chromatin. Structure 28, 105–110 e103.

35. K. Yan, J. Rousseau, R. O. Littlejohn, C. Kiss, A. Lehman, J. R. Rosenfeld, C. T. R. Stumpel, A. P. Stegmann, L. Robak, F. Scaglia, T. T. Nguyen, H. Fu, N. F. Ajeawung, M. V. Camurri, L. Li, A. Gardham, B. Panis, M. Almannai, M. J. Sacoto, B. Baskin, C. Ruivenkamp, D. Study, C. Study, M. T. Cho, T. Potjer, G. W. Santen, M. J. Parker, N. Canham, M. McKinnon, L. Potocki, J. MacKenzie, E. R. Roeder, P. M. Campeau and X. J. Yang (2017) Mutations in the chromatin regulator gene BRPF1 causes syndromic intellectual disability and deficient histone acetylation. Am. J. Hum. Genet. 100, 91–104.

36. F. Mattioli, E. Schaefer, A. Magee, P. Mark, G. M. Mancini, K. Dieterich, G. Von Allmen, M. Alders, C. Coutton, M. van Slegtenhorst, G. Vieville, M. Engelen, J. M. Cobben, J. Juusola, A. Pujol, J. L. Mandel and A. Piton (2017) Mutations in Histone Acetylase Modifier BRPF1 Cause an Autosomal-Dominant Form of Intellectual Disability with Associated Ptosis. Am J Hum Genet 100, 105–116.

37. S. Baker, J. Murrell, A. Nesbitt, K. Pechter, J. Balciuniene, X. Zhao, Z. Yu, E. Denenberg, E. DeChene, A. Wilkens, E. Bhoj, Q. Guan, M. Dulik, L. Conlin, A. Abou Tayoun, M. Luo, C. Wu, K. Cao, M. Sarmady, E. Bedoukian, J. Tarpinian, L. Medne, C. Skraban, M. Deardorff, I. Krantz, B. Krock and A. Santani (2019) Automated clinical exome reanalysis reveals novel diagnoses. J Mol Diagn 21, 38–48.

38. N. Pode-Shakked, O. Barel, B. Pode-Shakked, A. Eliyahu, A. Singer, O. Nayshool, N. Kol, A. Raas-Rothschild, E. Pras and M. Shohat (2019) BRPF1-associated intellectual disability, ptosis, and facial dysmorphism in a multiplex family. Mol Genet Genomic Med 10.1002/mgg3.665, e665.

39. Y. Mishima, S. Miyagi, A. Saraya, M. Negishi, M. Endoh, T. A. Endo, T. Toyoda, J. Shinga, T. Katsumoto, T. Chiba, N. Yamaguchi, I. Kitabayashi, H. Koseki and A. Iwama (2011) The Hbo1-Brd1/Brpf2 complex is responsible for global acetylation of H3K14 and required for fetal liver erythropoiesis. Blood 118, 2443–2453.

40. Y. Feng, A. Vlassis, C. Roques, M. E. Lalonde, C. Gonzalez-Aguilera, J. P. Lambert, S. B. Lee, X. Zhao, C. Alabert, J. V. Johansen, E. Paquet, X. J. Yang, A. C. Gingras, J. Cote and A. Groth (2016) BRPF3-HBO1 regulates replication origin activation and histone H3K14 acetylation. EMBO J 35, 176–192.

41. K. Yan, L. You, C. Degerny, M. Ghorbani, X. Liu, L. Chen, L. Li, D. Miao and X. J. Yang (2016) The chromatin regulator BRPF3 preferentially activates the HBO1 acetyltransferase but is dispensable for mouse development and survival. J Biol Chem 291, 2647–2663.

42. Y. Tao, C. Zhong, J. Zhu, S. Xu and J. Ding (2017) Structural and mechanistic insights into regulation of HBO1 histone acetyltransferase activity by BRPF2. Nucleic Acids Res 45, 5707–5719.

43. X. J. Yang (2004) The diverse superfamily of lysine acetyltransferases and their roles in leukemia and other diseases. Nucleic Acids Res. 32, 959–976.

44. C. D. Allis, S. L. Berger, J. Cote, S. Dent, T. Jenuwien, T. Kouzarides, L. Pillus, D. Reinberg, Y. Shi, R. Shiekhattar, A. Shilatifard, J. Workman and Y. Zhang (2007) New nomenclature for chromatin-modifying enzymes. Cell 131, 633–636.

45. Y. Yan, N. A. Barlev, R. H. Haley, S. L. Berger and R. Marmorstein (2000) Crystal structure of yeast Esa1 suggests a unified mechanism for catalysis and substrate binding by histone acetyltransferases. Mol. Cell 6, 1195–1205.

46. L. Li, M. Ghorbani, M. W. Hubshman, J. Rousseau, I. Thiffault, R. E. Schnur, C. Breen, R. Oegema, J. M. Weiss, Q. Waisfisz, S. Welner, H. Kingston, E. M. Boon, L. Basel-Salmon, O. Konen, H. Goldberg, L. Bazaq, S. Tzur, X. Bi, M. Bruccoleri, K. McWalter, M. Cho, M. Scarano, S. S. Brooks, S. S. Hughes, K. L. I. van Gassen, J. M. van Hagen, T. K. Pandita, P. B. Agrawal, P. M. Campeau and X. J. Yang (2020) Lysine acetyltransferase 8 is involved in cerebral development and syndromic intellectual disability. J. Clin. Investig. 130, 1431–1445.

47. S. Yang, Y. Yu, Y. Xu, F. Jian, W. Song, A. Yisimayi, P. Wang, J. Wang, J. Liu, L. Yu, X. Niu, J. Wang, Y. Wang, F. Shao, R. Jin, Y. Wang and Y. Cao (2024) Fast evolution of SARS-CoV-2 BA.2.86 to JN.1 under heavy immune pressure. Lancet Infect Dis 24, e70–e72.

48. O. R. Collaboration (2016) The ORFeome Collaboration: a genome-scale human ORF-clone resource. Nat Methods 13, 191–192.

49. M. Pacesa, O. Pelea and M. Jinek (2024) Past, present, and future of CRISPR genome editing technologies. Cell 187, 1076–1100.

50. L. Cong, F. A. Ran, D. Cox, S. Lin, R. Barretto, N. Habib, P. D. Hsu, X. Wu, W. Jiang, L. A. Marraffini and F. Zhang (2013) Multiplex genome engineering using CRISPR/Cas systems. Science 339, 819–823.

51. Y. C. J. Chey, L. Gierus, C. Lushington, J. C. Arudkumar, B. G. A, L. G. Staker, L. J. Robertson, C. Pfitzner, J. G. Kennedy, R. H. B. Lee, G. I. Godahewa, F. Adikusuma and P. Q. Thomas (2025) Optimal SpCas9- and SaCas9-mediated gene editing by enhancing gRNA transcript levels through scaffold poly-T tract reduction. BMC Genomics 26, 138.

52. S. Riesenberg, N. Helmbrecht, P. Kanis, T. Maricic and S. Paabo (2022) Improved gRNA secondary structures allow editing of target sites resistant to CRISPR-Cas9 cleavage. Nat Commun 13, 489.

53. G. Faure, M. Saito, M. E. Wilkinson, N. Quinones-Olvera, P. Xu, D. Flam-Shepherd, S. Kim, N. Reddy, S. Zhu, L. Evgeniou, E. V. Koonin, R. K. Macrae and F. Zhang (2025) TIGR-Tas: A family of modular RNA-guided DNA-targeting systems in prokaryotes and their viruses. Science 10.1126/science.adv9789, eadv9789.

54. S. Vazquez Torres, M. Benard Valle, S. P. Mackessy, S. K. Menzies, N. R. Casewell, S. Ahmadi, N. J. Burlet, E. Muratspahic, I. Sappington, M. D. Overath, E. Rivera-de-Torre, J. Ledergerber, A. H. Laustsen, K. Boddum, A. K. Bera, A. Kang, E. Brackenbrough, I. A. Cardoso, E. P. Crittenden, R. J. Edge, J. Decarreau, R. J. Ragotte, A. S. Pillai, M. Abedi, H. L. Han, S. R. Gerben, A. Murray, R. Skotheim, L. Stuart, L. Stewart, T. J. A. Fryer, T. P. Jenkins and D. Baker (2025) De novo designed proteins neutralize lethal snake venom toxins. Nature 639, 225–231.

55. M. Dalvai, J. Loehr, K. Jacquet, C. C. Huard, C. Roques, P. Herst, J. Cote and Y. Doyon (2015) A Scalable Genome-Editing-Based Approach for Mapping Multiprotein Complexes in Human Cells. Cell Rep 13, 621–633.

56. J. A. M. Mercer, S. J. DeCarlo, S. S. Roy Burman, V. Sreekanth, A. T. Nelson, M. Hunkeler, P. J. Chen, K. A. Donovan, P. Kokkonda, P. K. Tiwari, V. M. Shoba, A. Deb, A. Choudhary, E. S. Fischer and D. R. Liu (2024) Continuous evolution of compact protein degradation tags regulated by selective molecular glues. Science 383, eadk4422.

57. K. Luger, A. W. Mader, R. K. Richmond, D. F. Sargent and T. J. Richmond (1997) Crystal structure of the nucleosome core particle at 2.8 A resolution. Nature 389, 251–260.

58. M. Ullah, N. Pelletier, L. Xiao, S. P. Zhao, K. Wang, C. Degerny, S. Tahmasebi, C. Cayrou, Y. Doyon, S. L. Goh, N. Champagne, J. Cote and X. J. Yang (2008) Molecular architecture of quartet MOZ/MORF histone acetyltransferase complexes. Mol Cell Biol 28, 6828–6843.

59. I. Kitabayashi, Y. Aikawa, L. A. Nguyen, A. Yokoyama and M. Ohki (2001) Activation of AML1-mediated transcription by MOZ and inhibition by the MOZ-CBP fusion protein. EMBO J. 20, 7184–7196.

60. N. Champagne, N. R. Bertos, N. Pelletier, A. H. Wang, M. Vezmar, Y. Yang, H. H. Heng and X. J. Yang (1999) Identification of a human histone acetyltransferase related to monocytic leukemia zinc finger protein. J. Biol. Chem. 274, 28528–28536.

61. D. Agudelo, A. Duringer, L. Bozoyan, C. C. Huard, S. Carter, J. Loehr, D. Synodinou, M. Drouin, J. Salsman, G. Dellaire, J. Laganiere and Y. Doyon (2017) Marker-free coselection for CRISPR-driven genome editing in human cells. Nat Methods 14, 615–620.

