## Supplementary material for "P3a site-specific and cassette mutagenesis for seamless protein, RNA and plasmid engineering": Figs S1-S13

#### Supplementary Figure Legends

##### Fig. S1 Characteristics of mutagenic primer pairs with 3′-overhangs

A. Schematic illustrating two partially complementary primers with 3′-overhangs (Fig. 1B) directing synthesis from nicked newly synthesized DNA strands. In stark contrast, completely complementary primers (Fig. 1A) cannot do so. One primer pair (K9R-F and K9R-R) with 3′-overhangs was used to engineer the K9R mutation in histone H3.

B. Sequences showing how two primer pairs were designed to engineer K9R and K27R/M. The codons of K9 and K27 have already been replaced with those for R9 and R27, respectively. Conventionally, for histone proteins, amino acid residue positions are numbered starting from the second residue (alanine), instead of the first methionine. As a result, K9 and K27 are encoded by the $10^{th}$ and $28^{th}$ codons, respectively.

C. Sequences showing how a single primer pair was designed to engineer G34A and G34R point mutations.

##### Fig. S2 P3a mutagenesis to engineer missense or nonsense mutants of epigenetic regulators

A. Efficiency of engineering six KAT6A mutants and five mutants of its lysine acetyltransferase (HAT) domain. Asterisks in three C-terminal truncation mutants, such as G814*, refer to stop codons.

B. Efficiency of engineering seven mutants and two deletion mutants of KAT6B. A vector encoding residues 364-810 of KAT6B (GenBank accession, AF113514) was used for the experiments. The protein encoded by AF113514 carries an F551L substitution, possibly due to a sequencing error during cDNA cloning and analysis. F551 is invariant among different KAT6A and KAT6B proteins, so the mutant F551L was tested to assess the importance of F551, as denoted by two asterisks (**).

C. Efficiency in engineering 13 KAT8 mutants. The notation ‘R136C;R136H’ refers to generating two mutants R136C and R136H, with a single pair of primers (see Fig. 1C).

D. Efficiency in engineering mouse CBP mutants. Asterisks in two C-terminal truncation mutants such as R1868* refer to stop codons.

E. Sequence chromatograms of four representative plasmids sequenced for engineering the CBP mutants R1446C and R1446H. The top chromatogram is the wild-type and the remaining three are mutants.

**Fig. S3 Deleting or converting an epitope tag by P3a cassette mutagenesis**

A. Sequence of the region encoding the HA tag fused to the C-terminus of mouse CBP. The HA-coding sequence to be deleted is indicated with a horizontal solid line. This expression vector is 13.4 kb in size.

B. Sequence of the region after deletion and the two partially complementary primers (mCBP-F and mCBP-R) that were used for deletion by P3a cassette mutagenesis.

C. Sequence of two partially complementary primers (3Flag-F and 3Flag-R) used to convert the coding sequence for a FLAG tag on pAW48 (a pcDNA3.1 derivative) to that for a 3xFLAG tag by P3a site-directed mutagenesis.

**Fig. S4 Replacing a FLAG tag with an HA tag by P3a cassette mutagenesis**

A. Sequence of the region encoding the FLAG tag fused to the C-terminus of human SIN3A. The tag needs to be replaced with an HA tag for subsequent co-immunoprecipitation with partners to be expressed as FLAG fusion proteins.

B. Sequence of the resulting mutant and the two partially complementary primers (HA-F4 and HA-SIN3A) that were used for P3a cassette mutagenesis.

C. Sequence chromatograms of three candidate plasmids sequenced for engineering the replacement of the FLAG tag with an HA tag in the SIN3A-FLAG mammalian expression vector. The results indicate that all three are correct.

D. Sequence chromatograms of three candidate plasmids sequenced for engineering the replacement of the FLAG tag with an HA tag in the YEATS2-FLAG mammalian expression vector. The results indicate that two are correct and the third possesses a T insertion and 5-bp deletion just upstream from the coding sequence for the HA tag.

**Fig. S5 Optimization of a 12.8-kb expression vector by deletion of a 1.9-kb fragment**

A. Map of the 12.6-kb mammalian expression vector for FLAG-p300. The 1.9-kb fragment to be removed by P3a mutagenesis is highlighted in blue. The original p300 expression vector from Addgene (Cat. 23252) is ~12.8 kb, whereas the one shown here results from introduction of the FLAG coding sequence, deletion of a 92-bp fragment and removal of the 6xHis coding sequence (see Fig. 5B).

B. Map of the mutated 10.7-kb expression vector for FLAG-p300. d2k, deletion of ~2 kb.

C. Sequence of the resulting deletion mutant and the two partially complementary primers (d2k-F and d2K-R) that were used for P3a cassette mutagenesis.

D. Sequence chromatograms of four candidate plasmids sequenced for engineering the ~2-kb deletion. The top one is the wild-type, containing the ~2 kb fragment, the middle two are the mutant with the fragment deleted and the fourth possesses a large unwanted deletion.

**Fig. S6 Optimization of a pcDNA3.1 vector by simultaneous deletion of two fragments**

A. Map of a 5.4-kb pcDNA3.1-based mammalian expression vector with the coding sequence for an HA tag. Two fragments (390- and 430-bp) to be removed by P3a mutagenesis are indicated.

B. Map of the resulting 4.6-kb expression vector pAW51s (where the letter s refers to the smaller size than pAW51) encoding an HA tag.

C-D. Sequence of the resulting mutant plasmid and two pairs of partially complementary primers that were used for P3a cassette mutagenesis. DelF1-F and DelF1-R were used for deletion of the 430-bp fragment, whereas d390-F and d390-R were for removal of the 390-bp fragment

E. Efficiency for engineering the deletions. Condition A refers to PCR mutagenesis that started with 15 ng of the parental plasmid and 20 amplification cycles. Condition B is the same as A but only 5 ng of the parental plasmid was used. Condition C is the same as B, but 23 PCR cycles were carried out. The efficiency refers to deletion of either of the 390- and 430-bp fragments, whereas simultaneous deletion of both is much less effective, with an overall success rate of 4/25 (16%).

**Fig. S7 P3a mutagenesis for engineering restriction sites on plasmids for subcloning**

A. Single or double restriction sites were engineered into (or eliminated from) the indicated vectors for subsequent subcloning. For pAW51, an NheI site upstream from the coding sequence for the HA tag was first eliminated to produce pAW51a, which was subsequently used for the

introduction of an XhoI site and an NheI site downstream form the HA coding sequence (panel B), for subcloning of open reading frames flanked by these two sites. For pCL36, an HindIII site was inserted to produce two vectors possessing different reading frames downstream from the coding sequence for the HA tag (see Fig. S8). Notably, for the histone H4 plasmid, two restriction sites were engineered upstream and downstream from its open reading frame, indicating that the method is applicable for introducing two distant mutations simultaneously.
B. Sequence chromatograms of four representative plasmids sequenced for engineering the XhoI and NheI sites into pAW51a. All were correct mutants, yielding an ideal efficiency of 100%.

**Fig. S8 P3a mutagenesis for engineering an HindIII site on a plasmid for subcloning**

Sequence chromatograms of four representative plasmids sequenced for engineering a HindIII site into the mammalian expression vector pCL36. The two primers (36H-F and 36H1-R) were designed to engineer the HindIII site at different reading frames upstream from the coding sequence for the HA tag, thereby resulting in two plasmids (pCL36H and pCL36H1, respectively).

**Fig. S9 P3a and P3 mutagenesis methods for engineering LoxP sites and introducing deletions**

A. Sequence chromatograms of four representative plasmids sequenced for engineering the LoxP site into the mammalian expression vector for HA-tagged BRPF1. The two primers (LoxP-F and LoxP-R) were designed to engineer the LoxP site upstream from the coding sequence for the HA tag. Clones C1 and C2 were from P3a mutagenesis, whereas clones C3 and C4 were from P3 mutagenesis. For each mutagenesis reaction, plasmids from 6 colonies were analyzed by sequencing. All those from P3a mutagenesis contained the LoxP site, resulting in the ideal efficiency of 100%. For those from P3 mutagenesis, 4 harbored the LoxP site, leading to an efficiency of 4/6 (66.7%).

B. Efficiency of P3 cassette mutagenesis to engineer deletion of 6 fragments and insertion of a LoxP site. The asterisk denotes the average efficiency from these 7 mutations. For comparison, the efficiency for insertion of a LoxP site via P3a cassette mutagenesis is also shown here.

C. Photo of bacterial colonies from P3 and P3a mutagenesis reactions to introduce the LoxP site. For each mutagenesis reaction, 2 μl of the mixture from DpnI digestion was used to transform 20 μl DH5α competent cells. After transformation, the bacterial cells from the two mutagenesis reactions were plated out on the two sides of one LB-agar plate containing ampicillin.

**Fig. S10 P3a mutagenesis of Cas9 and guide RNA vectors for genome editing**

A. The Cas9 expression vectors pX330, pX459 and some of their derivatives possess two 44-bp repeats in the sgRNA scaffold and its downstream region. The repeats are problematic to PCR primers designed to mutate the scaffold or insert sgRNA-coding sequences just from the scaffold.

B. Two primers (d47bp-F and d47bp-R) were designed to remove the second 44-bp repeat along with its downstream TTT sequence. Four colonies were analyzed and all of them contained the correct mutations, leading to the ideal efficiency of 100%.

C. Sequence chromatograms of three representative plasmids from P3a mutagenesis to replace the TTTT stretch in the sgRNA scaffold with TTTC. The corresponding downstream AAAA was changed to GAAA. Two primers (TTTC-F and TTTC-R) were designed to engineer these two point mutations. All three clones that were sequenced contained the correct mutations, resulting in the ideal efficiency of 100%.

**Fig. S11 P3a mutagenesis to repair the coding sequence of human HAT1**

A. Cartoon illustrating an Ultramer primer pair (HAT1-uF and -uR) to insert the coding sequence for the N-terminal 84 residues of human HAT1.

B. Two Ultramer primers (HAT1-uF and -uR) were designed to insert the coding sequence for the N-terminal 84 residues of human HAT1.

C. Sequence chromatograms of three representative plasmids from P3a mutagenesis to insert the coding sequence for the N-terminal 85 residues of human HAT1. Plasmids from 12 colonies were analyzed by restriction digestion and 6 were found to possess the correct insert. Three of them were sequenced and all were correct, leading to an overall efficiency of 6/12 (50%).

**Fig. S12 P3a mutagenesis to replace the receptor-binding domain of SARS-CoV-2 spike protein.**

A. Cartoon showing an Ultra primer pair (Omi430-F and Omi430-R) to replace the coding sequence for the receptor-binding domain of the spike protein.

B. Two Ultra primers (Omi430-F and Omi430-R) were designed to replace the coding sequence replace the coding sequence for the receptor-binding domain of the spike protein.

C. Sequence chromatograms of four representative plasmids from P3a mutagenesis to replace the coding sequence for the receptor-binding domain of the spike protein. All of them were correct, leading to the ideal efficiency of 100%.

**Fig. S13 Different primer design strategies used to introduce deletion and insertion.**

A-B. Two distinct strategies to design a primer pair with 3′-overhangs for engineering deletion. The strategy depicted in panel A, as employed in the previous study [1], assumes that the overlapping region (except for the deletion) is identical to the wild-type template. This is not required for the strategy depicted in panel B, employed in the current study; the sequence could be a redesigned junction flanking the deletion. For example, for a protein-encoding gene, the sequence could be codon-optimized or even an epitope tag. The strategy depicted in panel B is actually a special case of replacement mutagenesis (see panel F).

C-D. Two different strategies to design a primer pair with 3′-overhangs for engineering insertion. For the strategy depicted in panel C, as employed in a previous study [1], the entire insertion is located within the overlapping regions of the two primers, with one primer carrying the insertion in the middle and another harboring the insertion at the 5′-end. By contrast, for the strategy shown in panel D, the inserted sequence is encoded within the overlapping region and the two 3′-overhangs. For large insertion, this reduces the costs by almost a half. The strategy depicted in panel D is a special case of replacement mutagenesis (see panel F).

E. Sequential insertion mutagenesis of two pairs of Ultramer primers allows circumventing the size limit of mutagenesis with one pair of such primers. Megamer™ single-stranded DNA fragments from IDT are another viable alternative to circumvent the size limit, although they are more expensive than regular oligos or Ultramers.

F. Scheme showing replacement (i.e., ‘cassette’) mutagenesis, where a primer pair containing the sequence of fragment A is used to replace fragment B. Conceptually, deletion and insertion are special cases where the respective sequences to be inserted and deleted are zero bp. This is because replacement mutagenesis converts fragment A to fragment B. For deletion, fragment B is zero bp in size, whereas for insertion, fragment A is zero bp.

**References:**


1. H. Liu and J. H. Naismith (2008) An efficient one-step site-directed deletion, insertion, single and multiple-site plasmid mutagenesis protocol. *BMC Biotechnol* **8**, 91.

**Figure S1**

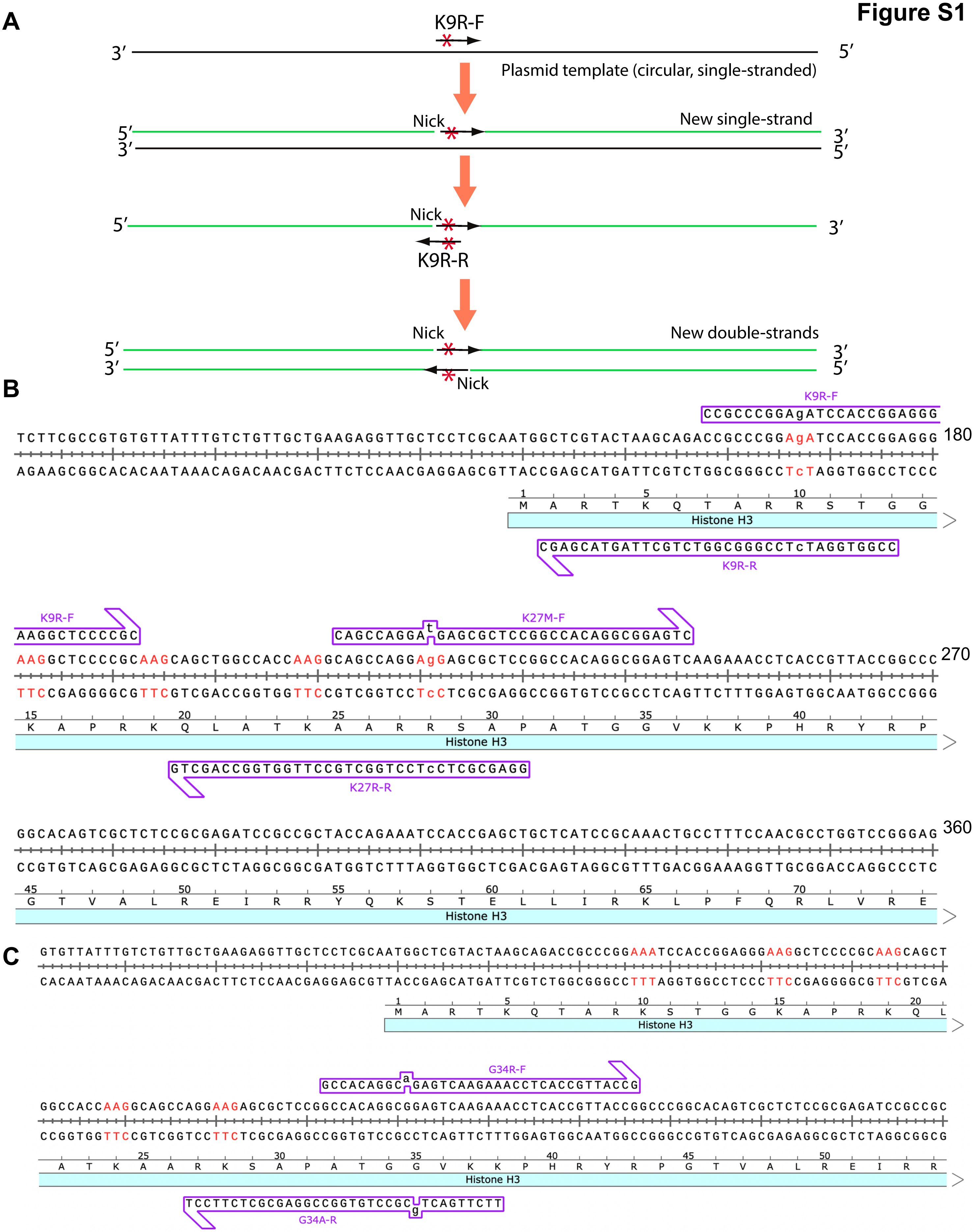
A
K9R-F
3′
5′
Plasmid template (circular, single-stranded)
Nick
New single-strand
5′
3′
3′
5′
Nick
5′
3′
K9R-R
Nick
New double-strands
5′
3′
3′
5′
Nick
B
K9R-F
CCGCCCGGAgATCCACCGGAGGG
TCTTCGCCGTGTGTTATTTGTCTGTTGCTGAAGAGGTTGCTCCTCGCAATGGCTCGTACTAAGCAGACCGCCCGGAgATCCACCGGAGGG
180
AGAAGCGGCACACAATAAACAGACAACGACTTCTCCAACGAGGAGCGTTACCGAGCATGATTCGTCTGGCGGGCCTcTAGGTGGCCTCCC
M A R T K Q T A R R S T G G
Histone H3
CGAGCATGATTCGTCTGGCGGGCCTcTAGGTGGCC
K9R-R
K9R-F
AAGGCTCCCCGC
K27M-F
CAGCCAGGA t GAGCGCTCCGGCCACAGGCGGAGTC
AAGGCTCCCCGCAAGCAGCTGGCCACCAAGGCAGCCAGGAgGAGCGCTCCGGCCACAGGCGGAGTCAAGAAACCTCACCGTTACCGGCCC
270
TTCCGAGGGGCGTTCGTCGACCGGTGGTTCCGTCGGTCCTcCTCGCGAGGCCGGTGTCCGCCTCAGTTCTTTGGAGTGGCAATGGCCGGG
K A P R K Q L A T K A A R R S A P A T G G V K K P H R Y R P
Histone H3
GTCGACCGGTGGTTCCGTCGGTCCTcCTCGCGAGG
K27R-R
GGCACAGTCGCTCTCCGCGAGATCCGCCGCTACCAGAAATCCACCGAGCTGCTCATCCGCAAACTGCCTTTCCAACGCCTGGTCCGGGAG
360
CCGTGTCAGCGAGAGGCGCTCTAGGCGGCGATGGTCTTTAGGTGGCTCGACGAGTAGGCGTTTGACGGAAAGGTTGCGGACCAGGCCCTC
G T V A L R E I R R Y Q K S T E L L I R K L P F Q R L V R E
Histone H3
C
GTGTTATTTGTCTGTTGCTGAAGAGGTTGCTCCTCGCAATGGCTCGTACTAAGCAGACCGCCCGGAAATCCACCGGAGGGAAGGCTCCCCGCAAGCAGCT
CACAATAAACAGACAACGACTTCTCCAACGAGGAGCGTTACCGAGCATGATTCGTCTGGCGGGCCTTTAGGTGGCCTCCCTTCCGAGGGGCGTTCGTCGA
M A R T K Q T A R K S T G G K A P R K Q L
Histone H3
G34R-F
GCCACAGGC a GAGTCAAGAAACCTCACCGTTACCG
GGCCACCAAGGCAGCCAGGAAGAGCGCTCCGGCCACAGGCGGAGTCAAGAAACCTCACCGTTACCGGCCCGGCACAGTCGCTCTCCGCGAGATCCGCCGC
CCGGTGGTTCCGTCGGTCCTTCTCGCGAGGCCGGTGTCCGCCTCAGTTCTTTGGAGTGGCAATGGCCGGGCCGTGTCAGCGAGAGGCGCTCTAGGCGGCG
A T K A A R K S A P A T G G V K K P H R Y R P G T V A L R E I R R
Histone H3
TCCTTCTCGCGAGGCCGGTGTCCGC g TCAGTTCTT
G34A-R

**Fig. S2**

**A**

| Mutation (KAT6A) | Mutants/colonies sequenced (%) |
|---|---|
| K604R | 3/4 (75.0%) |
| K350Kfs | 3/3 (100%) |
| R469* | 3/4 (75.0%) |
| G814* | 4/4 (100%) |
| L1061* | 3/4 (75.0%) |
| E1393* | 3/4 (75.0%) |
| L668F (HAT domain) | 2/3 (66.7%) |
| P682L (HAT domain) | 2/3 (66.7%) |
| Q743R (HAT domain) | 2/3 (66.7%) |
| Q740H;Q740L (HAT) | 2;1/3 (100%) |
| Total | 28/35 (80.0%) |

**B**

| Mutation (KAT6B) | Mutants/colonies sequenced (%) |
|---|---|
| F551L** | 3/3 (100%) |
| Q863H | 3/3 (100%) |
| E884G | 3/3 (100%) |
| S889C | 3/3 (100%) |
| D896N | 2/3 (66.7%) |
| R899H | 3/3 (100%) |
| E912k | 3/3 (100%) |
| del364-394 | 3/3 (100%) |
| del364-427 | 3/3 (100%) |
| Total | 26/27 (96.3%) |

**C**

| Mutation (KAT8) | Mutants/colonies sequenced (%) |
|---|---|
| E87D;E87Q | 2;1/3 (100%) |
| V90I | 2/3 (66.7%) |
| R136C;R136H | 1;2/3 (100%) |
| R140G;R140H | 2;1/4 (75.0%) |
| D190G | 2/6 (33.3%) |
| R224H | 2/3 (66.7%) |
| Y241C | 3/3 (100%) |
| P321L;P321H | 2;1/3 (100%) |
| R374W | 3/3 (100%) |
| Total | 24/31 (77.4%) |

**D**

| Mutation (CBP) | Mutants/colonies sequenced (%) |
|---|---|
| R1446C;R1446H | 2;1/4 (75.0%) |
| Y1503D;Y1503H | 1;2/4 (75.0%) |
| K1832E | 3/3 (100%) |
| R1869Q;R1869W | 2;1/3 (100%) |
| R1868* | 3/3 (100%) |
| R1881* | 3/3 (100%) |
| Total | 17/20 (85.0%) |

**E**

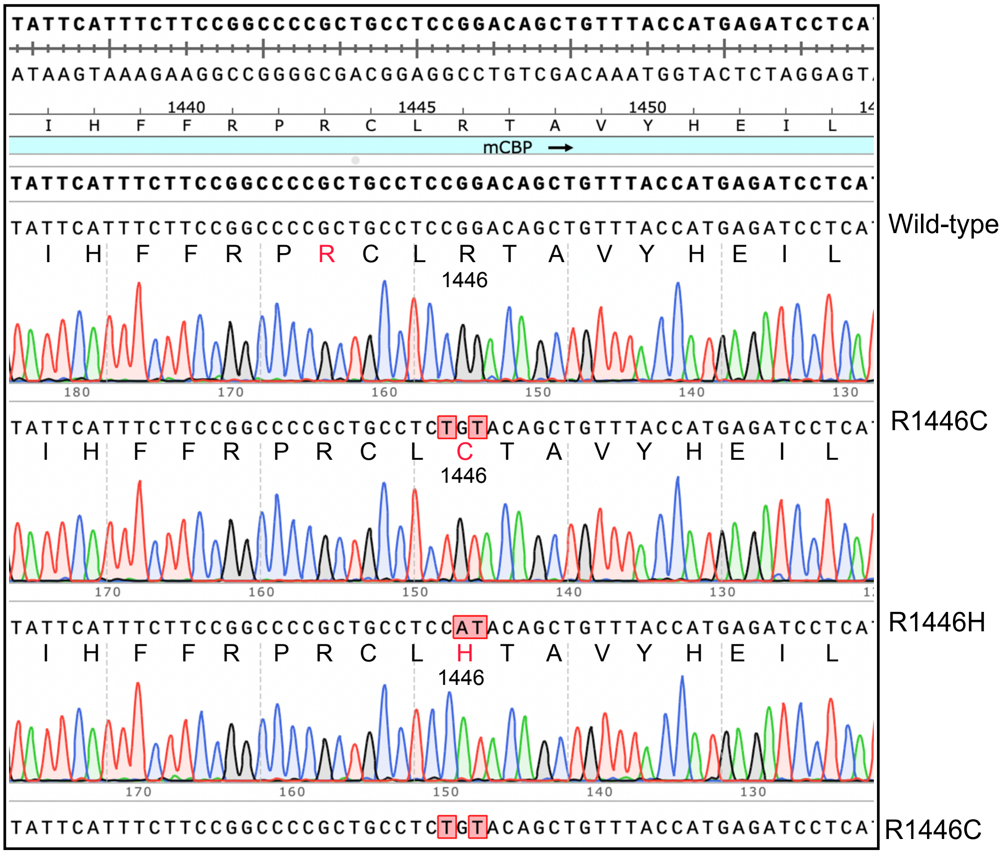

**Fig. S3**

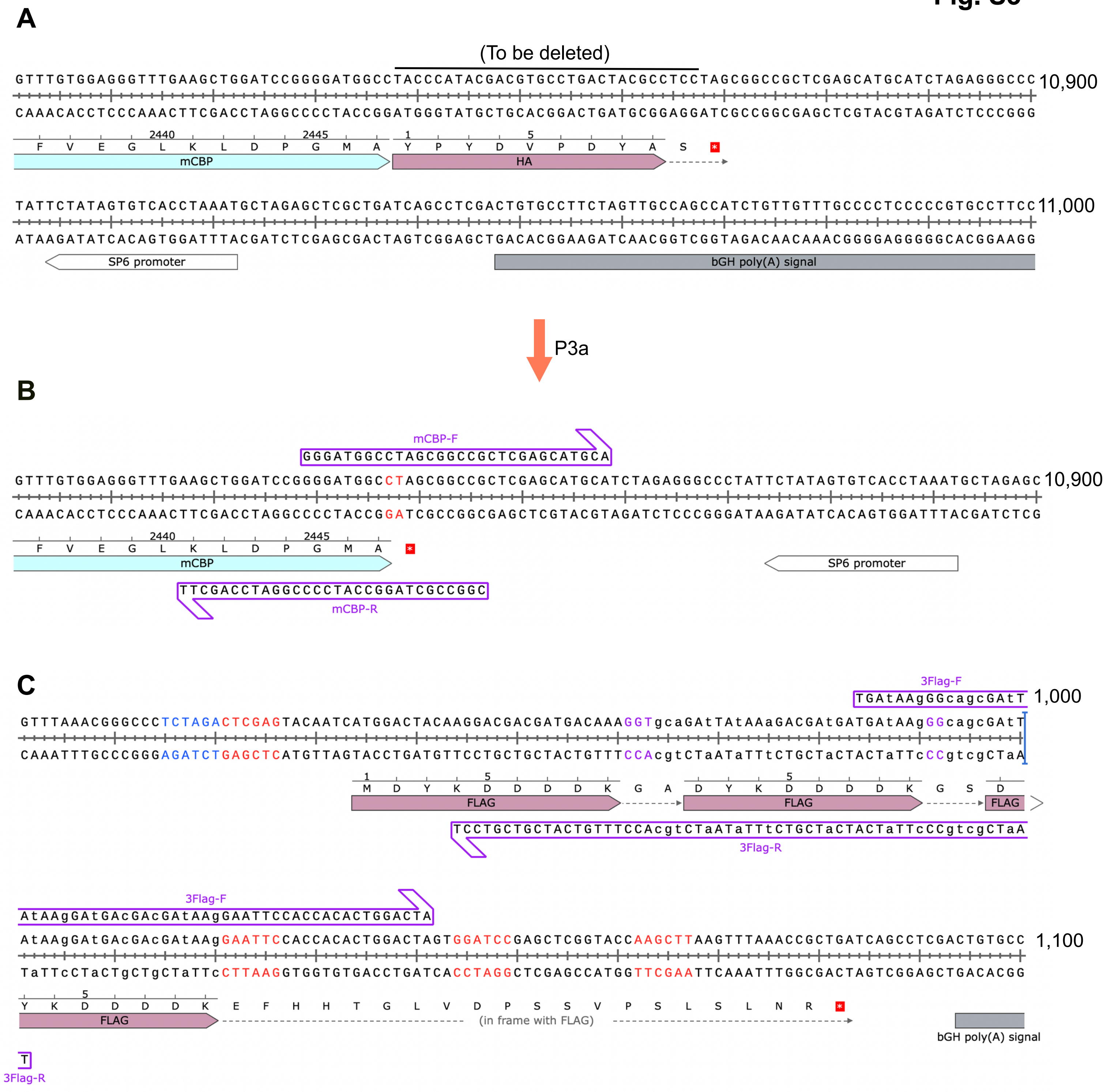
A
(To be deleted)
GTTTGTGGAGGGTTTGAAGCTGGATCCGGGGATGGCCTACCCATACGACGTGCCTGACTACGCCTCCTAGCGGCCGCTCGAGCATGCATCTAGAGGGCCC
10,900
CAAACACCTCCCAAACTTCGACCTAGGCCCCTACCGGATGGGTATGCTGCACGGACTGATGCGGAGGATCGCCGGCGAGCTCGTACGTAGATCTCCCGGG
2440
2445
F V E G L K L D P G M A Y P Y D V P D Y A S
mCBP
HA
TATTCTATAGTGTCACCTAAATGCTAGAGCTCGCTGATCAGCCTCGACTGTGCCTTCTAGTTGCCAGCCATCTGTTGTTTGCCCCTCCCCCGTGCCTTCC
11,000
ATAAGATATCACAGTGGATTTACGATCTCGAGCGACTAGTCGGAGCTGACACGGAAGATCAACGGTCGGTAGACAACAAACGGGGAGGGGGCACGGAAGG
SP6 promoter
bGH poly(A) signal
P3a
B
mCBP-F
GGGATGGCCTAGCGGCCGCTCGAGCATGCA
GTTTGTGGAGGGTTTGAAGCTGGATCCGGGGATGGCCTAGCGGCCGCTCGAGCATGCATCTAGAGGGCCCTATTCTATAGTGTCACCTAAATGCTAGAGC
10,900
CAAACACCTCCCAAACTTCGACCTAGGCCCCTACCGGATCGCCGGCGAGCTCGTACGTAGATCTCCCGGGATAAGATATCACAGTGGATTTACGATCTCG
2440
2445
F V E G L K L D P G M A
mCBP
SP6 promoter
TTCGACCTAGGCCCCTACCGGATCGCCGGC
mCBP-R
C
3Flag-F
TGAtAAgGGcagcGAtT
1,000
GTTTAAACGGGCCCTCTAGACTCGAGTACAATCATGGACTACAAGGACGACGATGACAAAGGTgcaGAtTAtAAaGACGAtGATGAtAAgGGcagcGAtT
CAAATTTGCCCGGGAGATCTGAGCTCATGTTAGTACCTGATGTTCCTGCTGCTACTGTTTCCAcgtCTaATaTTtCTGCTaCTACTaTTcCCgtcgCTaA
M D Y K D D D D K G A D Y K D D D D K G S D
FLAG
FLAG
FLAG
TCCTGCTGCTACTGTTTCCAcgtCTaATaTTtCTGCTaCTACTaTTcCCgtcgCTaA
3Flag-R
3Flag-F
AtAAgGAtGAcGAcGAtAAgGAATTCCACCACACTGGACTA
AtAAgGAtGAcGAcGAtAAgGAATTCCACCACACTGGACTAGTGGATCCGAGCTCGGTACCAAGCTTAAGTTTAAACCGCTGATCAGCCTCGACTGTGCC
1,100
TaTTcCTaCTgCTgCTaTTcCTTAAGGTGGTGTGACCTGATCACCTAGGCTCGAGCCATGGTTCGAATTCAAATTTGGCGACTAGTCGGAGCTGACACGG
Y K D D D D K E F H H T G L V D P S S V P S L S L N R
FLAG
(in frame with FLAG)
bGH poly(A) signal
T
3Flag-R

### Fig. S4

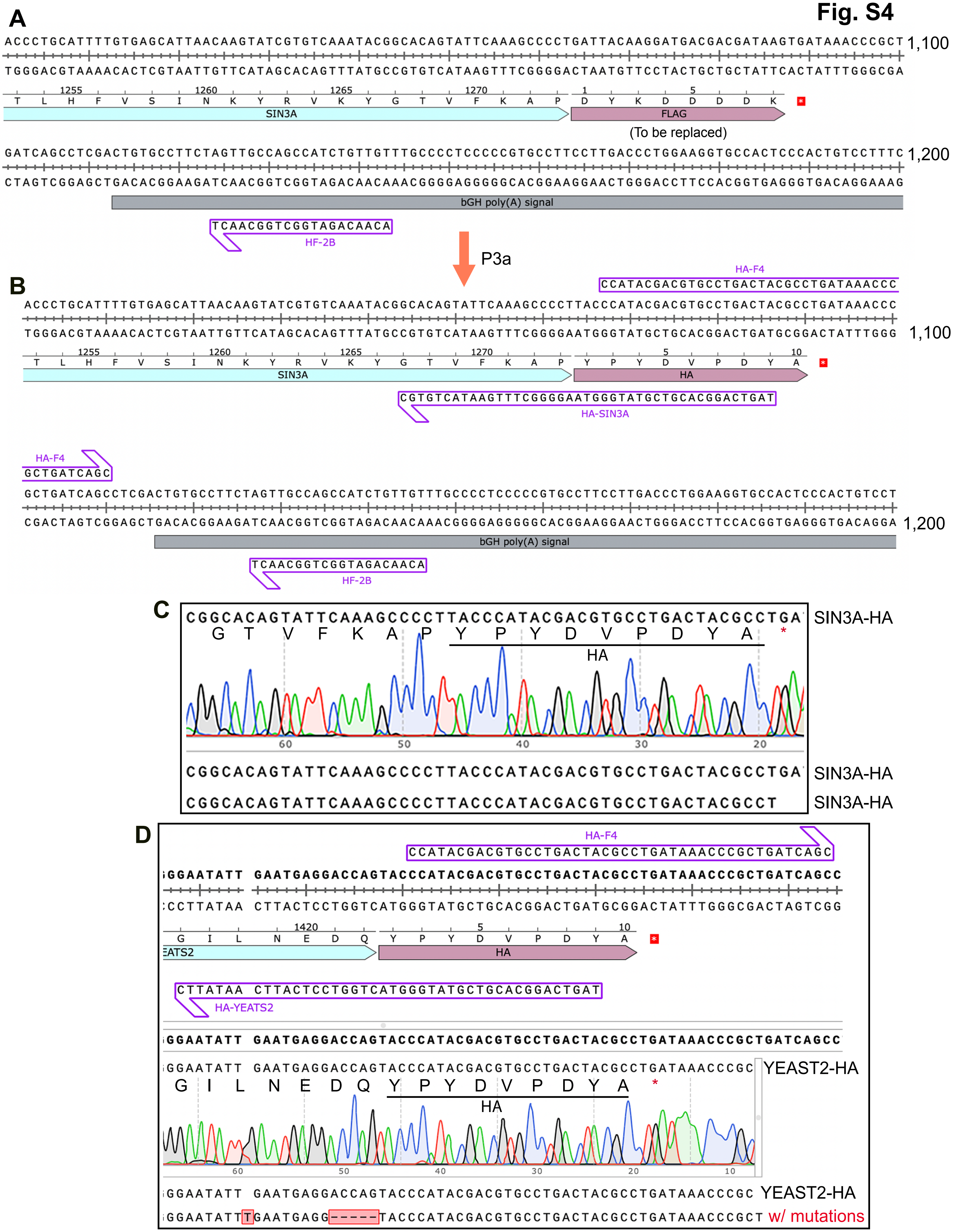

A
1,100
1255
1260
1265
1270
SIN3A
FLAG
(To be replaced)
1,200
bGH poly(A) signal
HF-2B
P3a
B
HA-F4
HA
HA-SIN3A
C
SIN3A-HA
D
EATS2
HA-YEATS2
YEAST2-HA
w/ mutations

**Fig. S5**

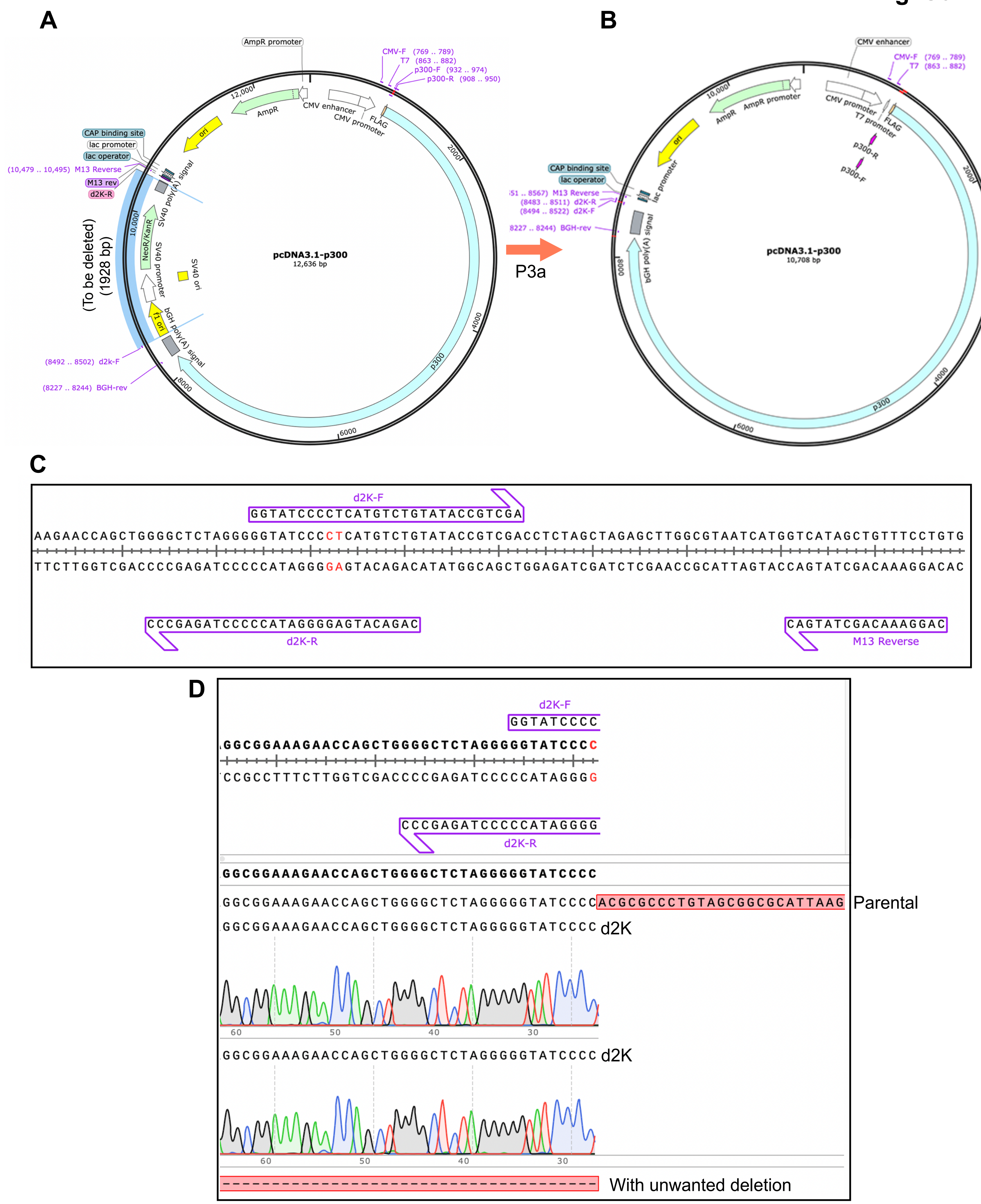
A
B
C
D
pcDNA3.1-p300
12,636 bp
AmpR promoter
CMV-F (769 .. 789)
T7 (863 .. 882)
p300-F (932 .. 974)
p300-R (908 .. 950)
CMV enhancer
CMV promoter
FLAG
AmpR
ori
CAP binding site
lac promoter
lac operator
(10,479 .. 10,495) M13 Reverse
M13 rev
d2K-R
SV40 poly(A) signal
NeoR/KanR
SV40 promoter
SV40 ori
f1 ori
bGH poly(A) signal
(To be deleted)
(1928 bp)
(8492 .. 8502) d2k-F
(8227 .. 8244) BGH-rev
p300
P3a
pcDNA3.1-p300
10,708 bp
(8483 .. 8511) d2K-R
(8494 .. 8522) d2K-F
8227 .. 8244) BGH-rev
p300-R
p300-F
d2K-F
GGTATCCCCTCATGTCTGTATACCGTCGA
AAGAACCAGCTGGGGCTCTAGGGGGTATCCCCTCATGTCTGTATACCGTCGACCTCTAGCTAGAGCTTGGCGTAATCATGGTCATAGCTGTTTCCTGTG
TTCTTGGTCGACCCCGAGATCCCCCATAGGGGAGTACAGACATATGGCAGCTGGAGATCGATCTCGAACCGCATTAGTACCAGTATCGACAAAGGACAC
CCCGAGATCCCCCATAGGGGAGTACAGAC
d2K-R
CAGTATCGACAAAGGAC
M13 Reverse
d2K-F
GGTATCCCC
GGCGGAAAGAACCAGCTGGGGCTCTAGGGGGTATCCCC
CCGCCTTTCTTGGTCGACCCCGAGATCCCCCATAGGGG
CCCGAGATCCCCCATAGGGG
d2K-R
GGCGGAAAGAACCAGCTGGGGCTCTAGGGGGTATCCCC
GGCGGAAAGAACCAGCTGGGGCTCTAGGGGGTATCCCCACGCGCCCTGTAGCGGCGCATTAAG
Parental
GGCGGAAAGAACCAGCTGGGGCTCTAGGGGGTATCCCC d2K
GGCGGAAAGAACCAGCTGGGGCTCTAGGGGGTATCCCC d2K
With unwanted deletion

Fig. S6

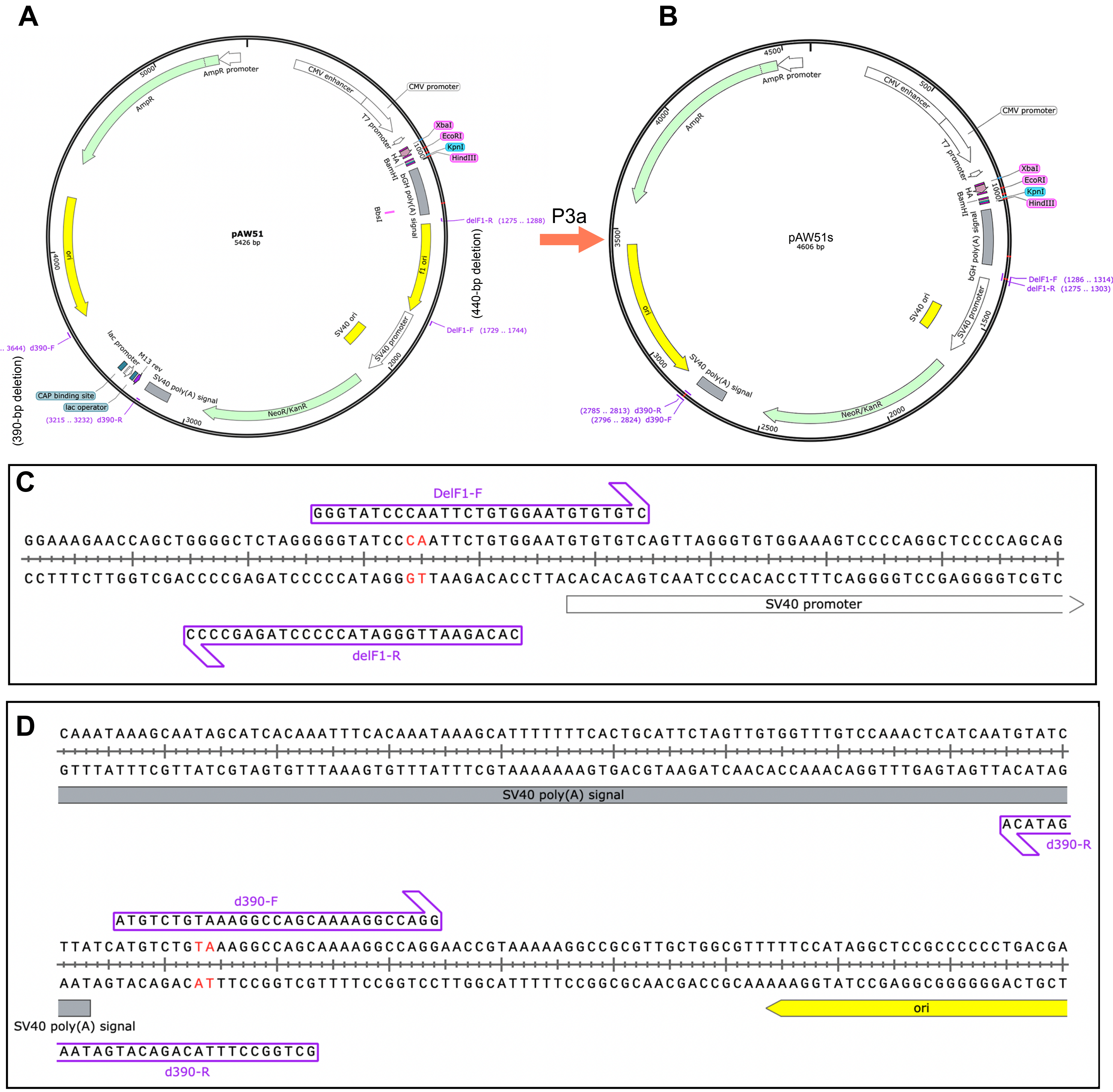


E

| Deletion | Condition A | B | C |
|---|---|---|---|
| 390 bp | 2 | 3 | 1 |
| 430 bp | 4 | 4 | 6 |
| Both | 1 | 3 | 0 |
| Wild-type | 3 | 1 | 1 |
| Total | 9 | 8 | 8 |
| Efficiency | 6/9 (66.7%) | 7/8 (87.5%) | 7/8 (87.5%) |

**Fig. S7**

**A**

| New restriction site | Mutants/colonies sequenced (%) |
| --- | --- |
| NheI elimination (pAW51) | 4/4 (100%) |
| XhoI/NheI (pAW51a) | 4/4 (100%) |
| HindIII (pCL36) | 4/4 (100%) |
| HindIII (USP10) | 3/4 (75%) |
| HindIII (SIN3B) | 4/9 (44.4%) |
| XhoI/BamHI (Histone H4) | 2/4 (50%) |
| Total | 21/29 (72.4%) |

**B**

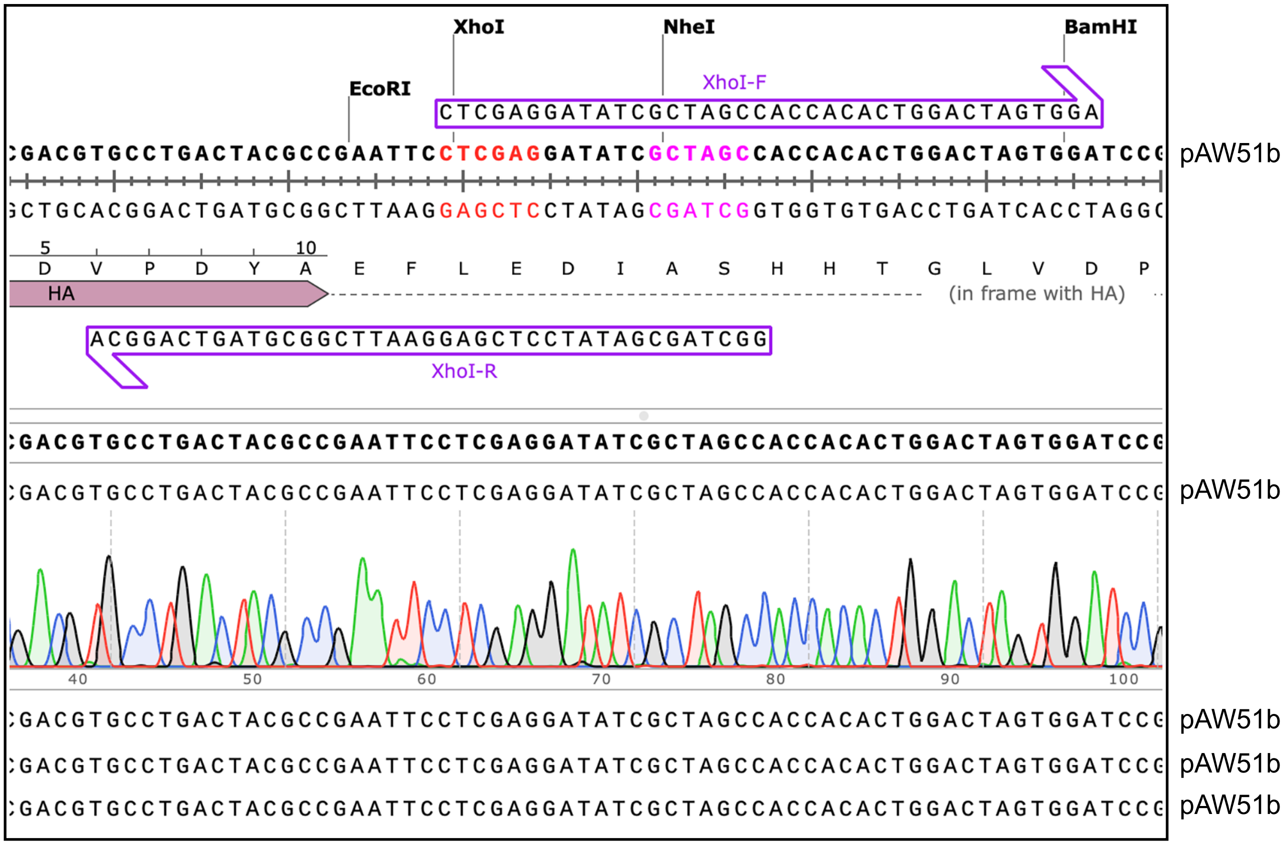


t

### Fig. S8

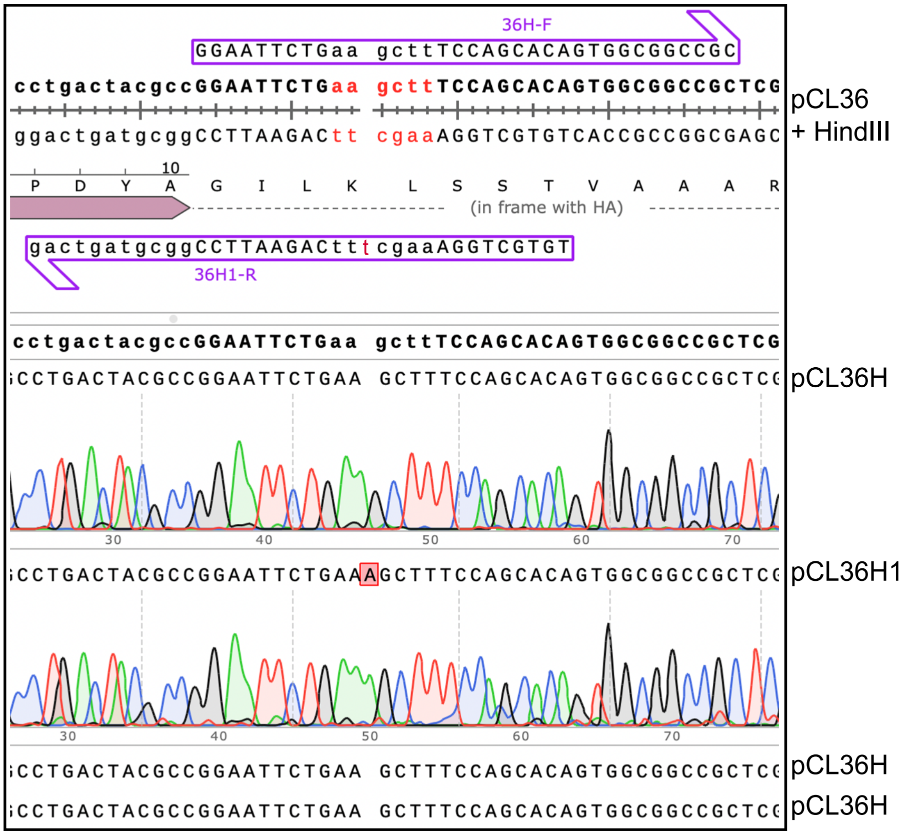
36H-F
GGAATTCTGaa gcttTCCAGCACAGTGGCGGCCGC
cctgactacgccGGAATTCTGaa gcttTCCAGCACAGTGGCGGCCGCTCG
ggactgatgcggCCTTAAGACtt cgaaAGGTCGTGTCACCGCCGGCGAGC
pCL36
+ HindIII
10
P D Y A G I L K L S S T V A A A R
(in frame with HA)
gactgatgcggCCTTAAGACttcgaaAGGTCGTGT
36H1-R
cctgactacgccGGAATTCTGaa gcttTCCAGCACAGTGGCGGCCGCTCG
CCTGACTACGCCGGAATTCTGAA GCTTTCCAGCACAGTGGCGGCCGCTCG
pCL36H
30
40
50
60
70
CCTGACTACGCCGGAATTCTGAAAGCTTTCCAGCACAGTGGCGGCCGCTCG
pCL36H1
30
40
50
60
70
CCTGACTACGCCGGAATTCTGAA GCTTTCCAGCACAGTGGCGGCCGCTCG
pCL36H
CCTGACTACGCCGGAATTCTGAA GCTTTCCAGCACAGTGGCGGCCGCTCG
pCL36H

### Fig. S9

A

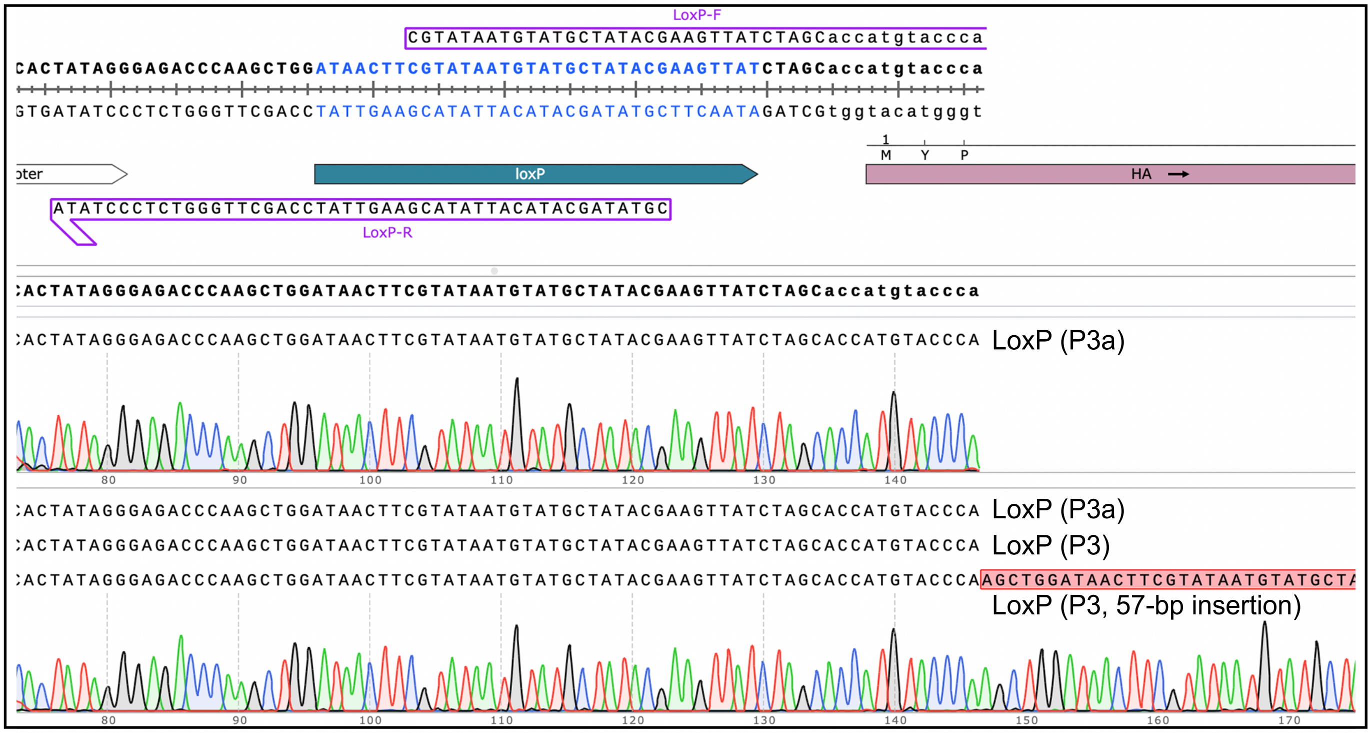


B

Efficiency of P3 cassette mutagenesis in egnieering deletion and insertion

| Deletion Mutants | Plasmids Sequenced (%) | Deletion (bp) |
|---|---|---|
| BRPF1-dN131 | 4/6 (66.7%) | 393 |
| BRPF1-dN204 | 3/6 (50%) | 612 |
| BRPF2-d31-80 | 2/6 (33.3%) | 150 |
| p300-dN1301 | 2/5 (40%) | 3,903 |
| CBP-dN1068 | 6/7 (85.7%) | 3,204 |
| p300 vector_d2K | 3/6 (50%) | 1,928 |
| LoxP (P3) | 4/6 (66.7%) | 34 bp (insertion) |
| LoxP (P3a) | 6/6 (100%) | 34 bp (insertion) |
| Total | 24/42 (57.1%; 56.1±16.7%)* | n/a |

C

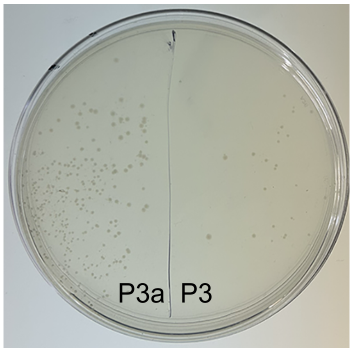

Fig. S10

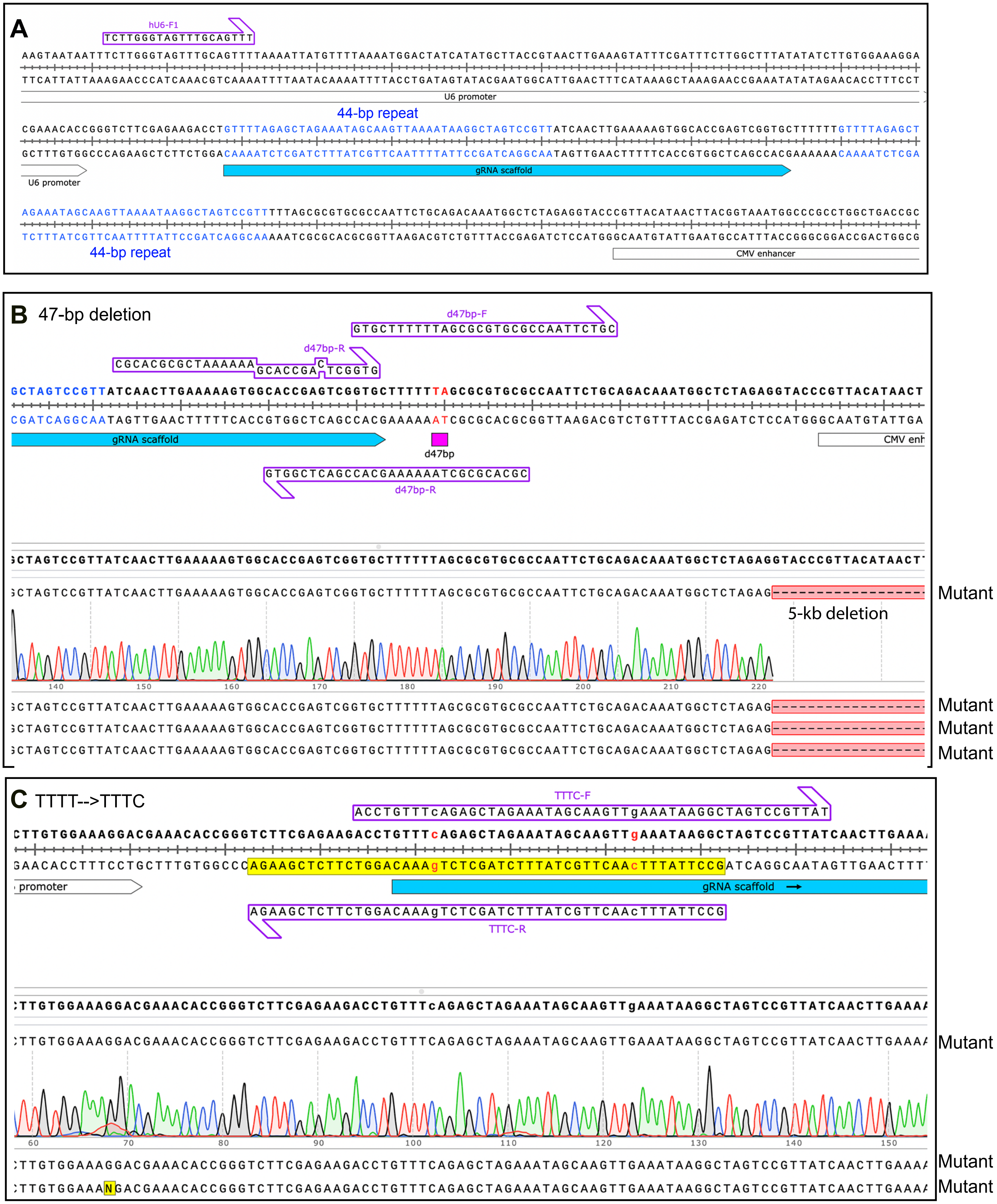
A
hU6-F1
TCTTGGGTAGTTTGCAGTTT
U6 promoter
44-bp repeat
gRNA scaffold
U6 promoter
44-bp repeat
CMV enhancer
B 47-bp deletion
d47bp-F
GTGCTTTTTTAGCGCGTGCGCCAATTCTGC
d47bp-R
gRNA scaffold
d47bp
CMV enh
GTGGCTCAGCCACGAAAAAATCGCGCACGC
d47bp-R
Mutant
5-kb deletion
Mutant
Mutant
Mutant
C TTTT-->TTTC
TTTC-F
ACCTGTTTcAGAGCTAGAAATAGCAAGTTgAAATAAGGCTAGTCCGTTAT
promoter
gRNA scaffold
AGAAGCTCTTCTGGACAAAgTCTCGATCTTTATCGTTCAAcTTTATTCCG
TTTC-R
Mutant
Mutant
Mutant

### Fig. S11

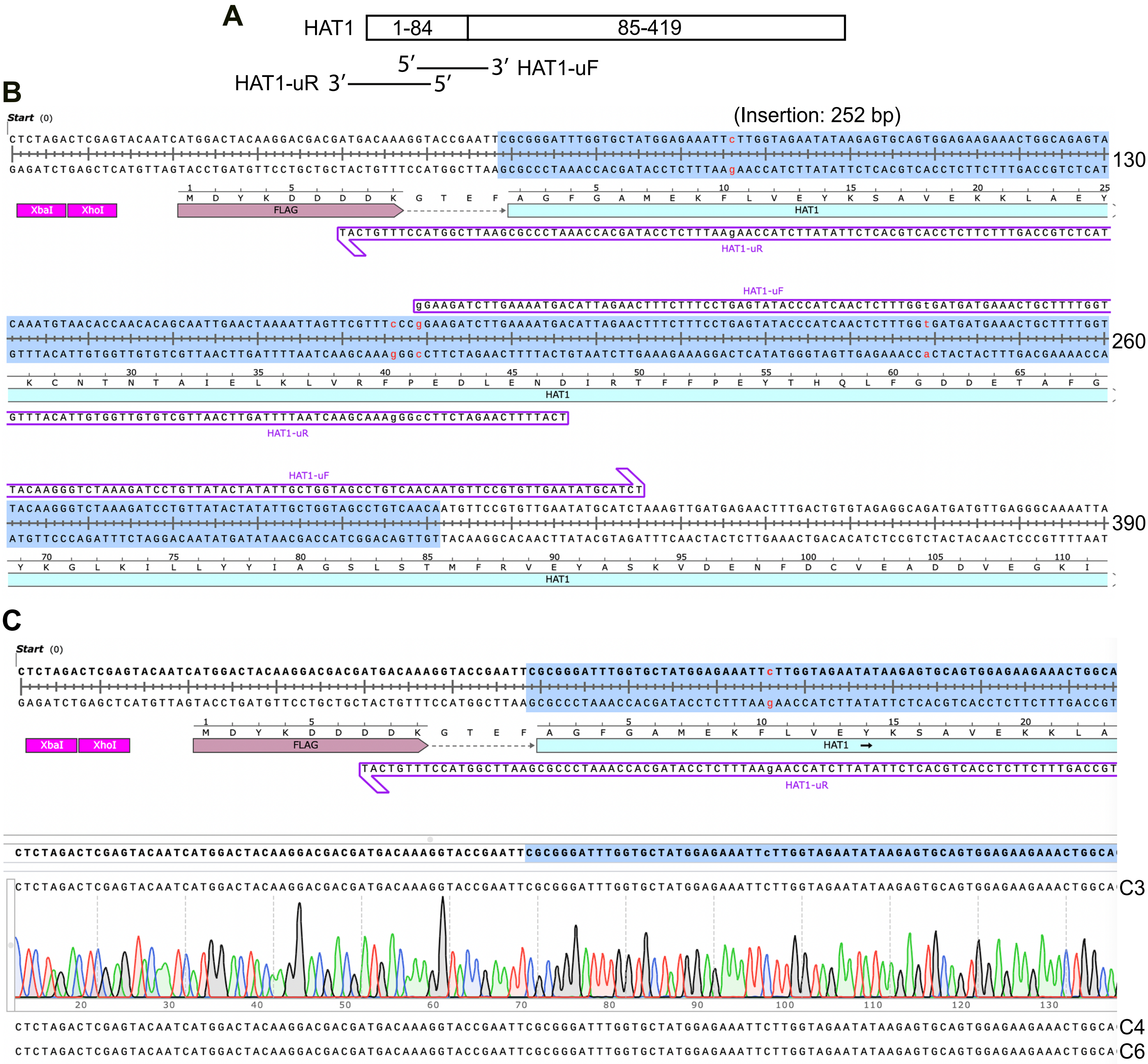
A
HAT1
1-84
85-419
5′ 3′ HAT1-uF
HAT1-uR 3′ 5′
B
Start (0)
(Insertion: 252 bp)
130
260
390
XbaI
XhoI
FLAG
HAT1
HAT1-uR
HAT1-uF
C
Start (0)
XbaI
XhoI
FLAG
HAT1
HAT1-uR
C3
C4
C6

Fig. S12

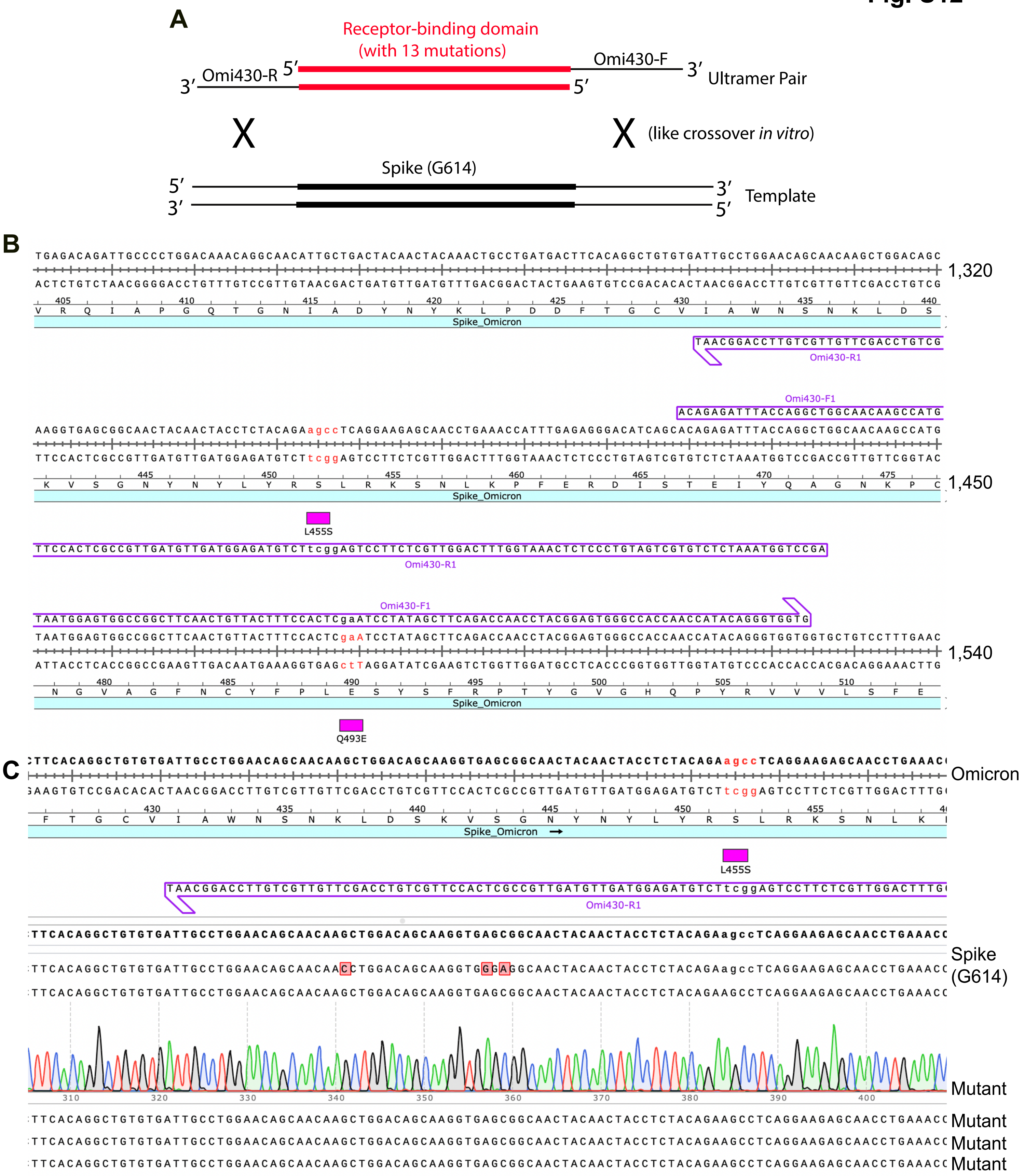

A
Receptor-binding domain
(with 13 mutations)
Omi430-R
Omi430-F
Ultramer Pair
(like crossover *in vitro*)
Spike (G614)
Template
B
1,320
Spike_Omicron
Omi430-R1
Omi430-F1
1,450
L455S
1,540
Q493E
C
Omicron
Spike_Omicron
L455S
Omi430-R1
Spike
(G614)
Mutant
Mutant
Mutant
Mutant

Fig. S13

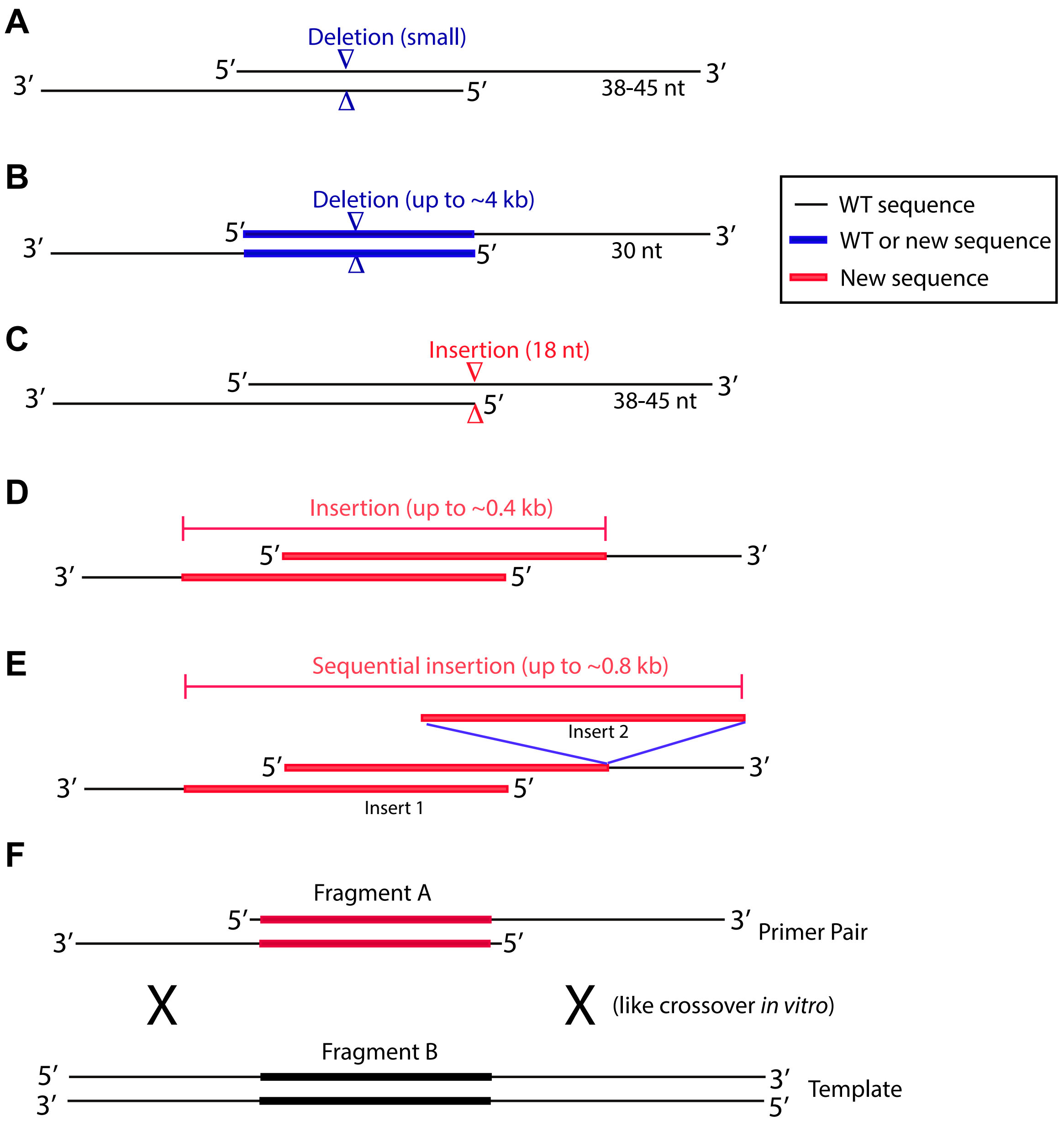
A
Deletion (small)
5′
3′
3′
5′
38-45 nt
B
Deletion (up to ~4 kb)
5′
3′
3′
5′
30 nt
WT sequence
WT or new sequence
New sequence
C
Insertion (18 nt)
5′
3′
3′
5′
38-45 nt
D
Insertion (up to ~0.4 kb)
5′
3′
3′
5′
E
Sequential insertion (up to ~0.8 kb)
Insert 2
5′
3′
3′
5′
Insert 1
F
Fragment A
5′
3′
Primer Pair
3′
5′
X
X
(like crossover *in vitro*)
Fragment B
5′
3′
Template
3′
5′
